# Schwann cells promote sensory neuron excitability during development

**DOI:** 10.1101/2022.10.31.514415

**Authors:** Husniye Kantarci, Pablo D. Elvira, Arun P. Thottumkara, Manasi Iyer, Lauren J. Donovan, Micaela Quinn Dugan, Nicholas Ambiel, Emma M. O’Connell, Alejandro Granados, Hong Zeng, Nay L. Saw, Amanda Brosius Lutz, Steven A. Sloan, Erin E. Gray, Khanh V. Tran, Aditi Vichare, Ashley K. Yeh, Alexandra E. Münch, Max Huber, Aditi Agrawal, Maurizio Morri, Mehrdad Shamloo, Thomas Anthony Anderson, Vivianne L. Tawfik, J. Du Bois, J. Bradley Zuchero

## Abstract

Excitability—the ability to fire action potentials—is a signature feature of neurons. How neurons become excitable during development, and whether excitability is an intrinsic property of neurons or requires signaling from glial cells, remain unclear. Here we demonstrate that Schwann cells, the most abundant glia in the peripheral nervous system, promote somatosensory neuron excitability during development. We find that Schwann cells secrete prostaglandin E_2_, which is necessary and sufficient to induce developing somatosensory neurons to express normal levels of voltage-gated sodium channels and fire action potential trains. Inactivating this signaling pathway specifically in Schwann cells selectively impairs the maturation of nociceptor and proprioceptor somatosensory neuron subtypes, leading to corresponding sensory defects in thermoception, inflammatory pain, and proprioception. Our studies thus reveal a cell non-autonomous mechanism by which glia regulate neuronal excitability to enable the development of normal sensory functions.

## INTRODUCTION

Excitability is a defining feature of all neurons. Dysregulation of excitability underpins diverse pathologies of the nervous system including chronic pain, epilepsy, and neuropsychiatric disease^1–8^. During development, neuronal maturation to achieve proper electrical function of the nervous system is vital to human health. However, the mechanisms that promote neuronal excitability during development are poorly understood.

Dorsal root ganglia (DRG) somatosensory neurons innervate peripheral tissues to transmit sensory inputs to the central nervous system (CNS). After specification during embryonic development, DRG neurons are initially electrically inexcitable and only become excitable once they begin to express voltage-gated sodium channels (Na_V_s)—at around embryonic (E) day 12- 14 in mouse (E16 in rat)^9–14^. From late embryogenesis until after birth, DRG neurons further mature to become fully responsive to physiological sensory stimuli^14^. Developmentally, initiation and maturation of DRG neuron excitability coincides with the appearance of Schwann cell precursors in peripheral nerves (∼E12) and their subsequent differentiation into Schwann cells (E14-E16)^15, 16^. Given that glia actively regulate several important aspects of nervous system development and function (e.g., synaptogenesis)^17^, we wished to investigate whether glia may also regulate the initiation and/or maturation of neuronal excitability. While glia can indirectly affect neuronal excitability (e.g., by regulating ion homeostasis and synapse strength^18–24^), whether glia directly regulate intrinsic neuronal excitability is unknown. A major unanswered question is whether neurons require signaling from glia to become excitable during development, and if so, what are the mechanism(s) that underly this process.

Here we reveal an unexpected, instructive role for glia in inducing neuronal excitability during development. We found that highly purified embryonic DRG neurons were hypoexcitable and largely failed to fire action potential trains in the absence of glia. Adding back media containing Schwann cell-secreted signaling molecules was sufficient to induce neurons to express voltage-gated sodium channels (Na_V_s) and to generate action potentials. We identified a single signaling molecule released by Schwann cells—the prostaglandin PGE_2_—that was necessary and sufficient to promote Na_V_ expression and neuronal excitability. Genetically ablating this signaling pathway caused a profound neurodevelopmental defect in somatosensory neurons, including reduced induction of Na_V_ expression and neuronal hypoexcitability. Single cell RNA-seq revealed that mice lacking PGE_2_ production in Schwann cells had specific defects in the development of two important classes of sensory neurons— nociceptors and proprioceptors—and exhibited corresponding behavioral defects in these sensory modalities. Collectively, our findings reveal that excitability—the most fundamental property of neurons—requires active signaling from glial cells, and raise broader questions about the developmental role of glia in disorders of the nervous system.

## RESULTS

### Schwann cells increase neuronal excitability by inducing ion channel expression

To test whether glia regulate DRG neuron excitability during development, we first required a method to obtain highly pure cell cultures of embryonic DRG neurons that lack glia. In vivo, DRG neurons are closely associated with myelinating or nonmyelinating Schwann cells, which together cover nearly the entire surface of all peripheral axons^25, 26^. We developed a rapid (∼5 hr) immunopanning method to isolate embryonic rat DRG neurons to greater than 97% purity, devoid of contaminating Schwann cells, fibroblasts, and blood cells (Figures 1A and S1A)^27, 28^. Previous studies used unpurified DRG neurons contaminated with glia^29, 30^ or antimitotic drugs to kill non-neuronal cells over 1-2 weeks^31^, confounding efforts to directly measure the effect of glial cells in regulating neuronal excitability. The ability to rapidly isolate embryonic DRG neurons in pure neuronal cultures allowed us to test whether DRG neurons were normally excitable in the absence of glial cells. Unexpectedly, in response to injected current, immunopanned DRG neurons exhibited a high threshold for firing an action potential (77 ± 6 pA) and rarely propagated multiple action potential spikes (Figures 1B–1D). This result is in stark contrast to the normal ability of DRG neurons to fire repetitive trains of action potentials in culture^30, 32, 33^ and in vivo^34–38^. Thus, in the absence of glial cells, cultured DRG neurons are hypoexcitable.

**Figure 1.**
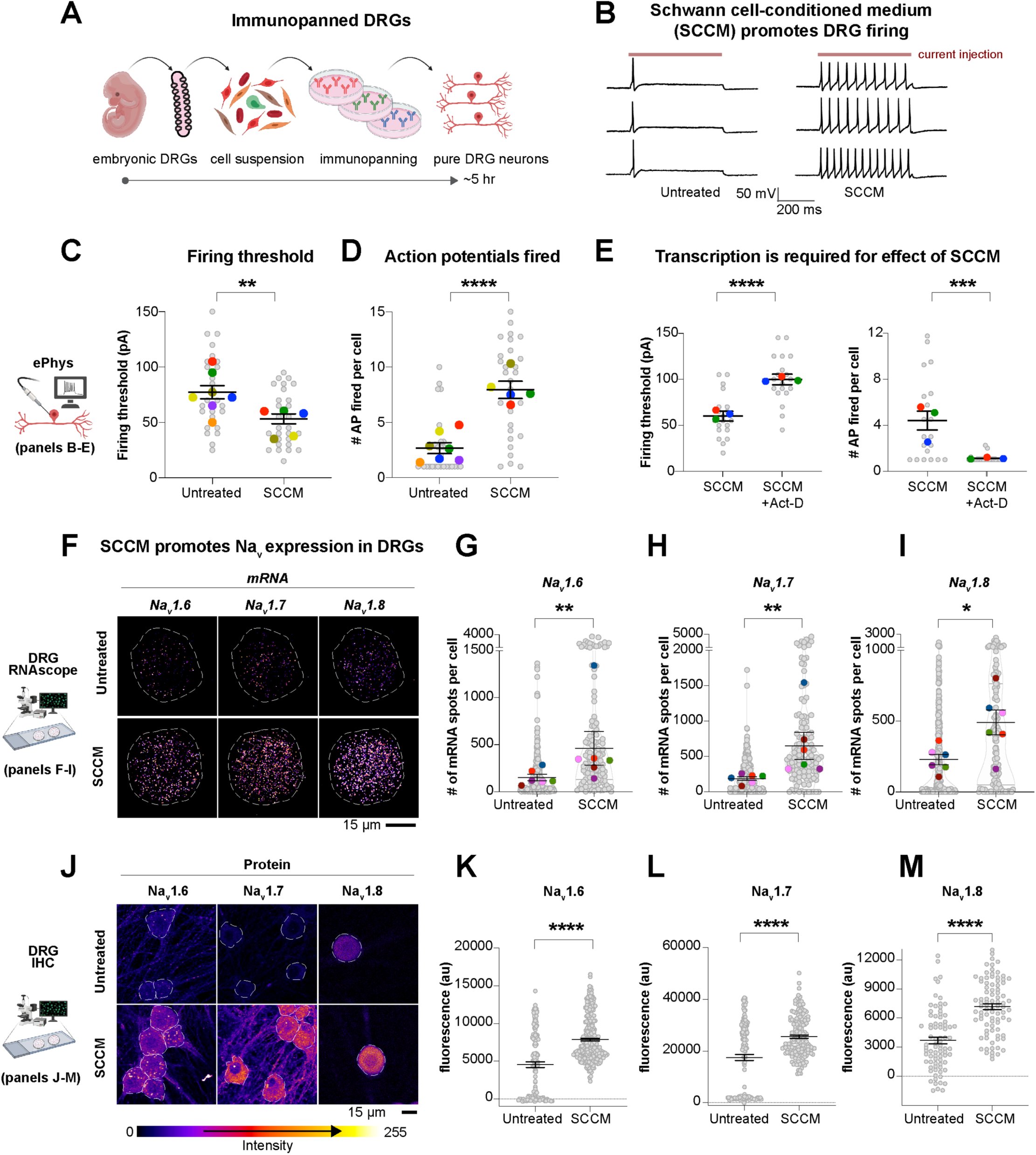
Schwann cells promote neuronal excitability and Na_V_ expression. (**A**) Cartoon depicting dissection, dissociation, and panning steps of the DRG neuron immunopanning method (see Methods). (**B**) Schwann cell-conditioned medium (SCCM) increased action potential firing frequency. Representative recordings of untreated and SCCM-treated DRG neurons. Traces show activity over 500 ms with 150 pA of current injection (see also Figure S4 for the raster plots summarizing the activity of all analyzed cells). (**C**) DRG neurons treated with SCCM required significantly lower levels of current injections for action potential propagation. (**D**) DRG neurons treated with SCCM fired significantly more action potentials at suprathreshold current injections. (**E**) Transcription inhibitor actinomycin-D (Act-D) blocked the excitability-inducing effect of SCCM. DRG neurons typically fired no more than one action potential and exhibited a higher firing threshold when Act-D was added to SCCM media. *n* = 20 total cells from 3 distinct biological replicates. For C-E: gray circles, response of an individual DRG neuron; colored circles, the average response of cells in each biological replicate. Mean ± SEM of all cells is shown; p values compare cells in unpaired t-test. (**F**) SCCM treatment enhanced *Na_V_* transcription in DRG neurons. Images show RNAscope for indicated genes in single representative DRG neurons that were either untreated (top) or treated with SCCM overnight (bottom). (**G** to **I**) Quantification of mRNA spots using FishQuant. Gray circles, mRNA number of an individual DRG neuron for the indicated genes; colored circles, the average # of mRNAs per cell in each biological replicate. *n* = 6 distinct biological replicates per treatment group. Mean ± SEM is shown for biological replicates. p values compare biological replicates in a paired t-test. (**J**) SCCM treatment enhanced Na_V_ protein levels in DRG neurons. Images show immunohistochemistry for indicated genes in single representative DRG neurons that were either untreated (top) or treated with SCCM overnight (bottom). (**K** to **M**) Quantification of fluorescence levels using Cellpose. Gray circles, fluorescent intensity of an individual DRG neuron for the indicated genes. *n* = 80 to 328 cells from 1-4 distinct biological replicates per treatment group. Mean ± SEM is shown for cells. p values compare cells in a one-way ANOVA and Tukey test (see also Figure S3B). *p<0.05, **p<0.01, ***p<0.001, ****p<0.0001.

Are Schwann cells required to promote the excitability of DRG neurons? To test this idea, we cultured acutely isolated rat Schwann cells and collected secreted factors as Schwann cell-conditioned medium (SCCM). Treating DRG neurons with SCCM for 16–28 hours decreased the current threshold of activation (Figure 1C; 53 ± 4 pA) and enhanced the ability of neurons to fire multiple action potentials at suprathreshold current injections (Figures 1B–1D). We measured no significant change in neuronal resting membrane potential, which could indirectly affect DRG neuron firing properties^39^ (Figure S1B). Schwann cells isolated from postnatal rats expressed immature Schwann cell markers but not inflammatory genes associated with disease or injury states^40, 41^ (Figure S2), suggesting the excitability-inducing effect did not result from an inflammatory form of Schwann cells. Together, these results demonstrated that Schwann cells secrete molecule(s) that increase the excitability of sensory neurons.

The effect of SCCM on excitability was dependent on the transcription of new mRNA, as treating DRG neurons with the transcription inhibitor actinomycin-D blocked the ability of SCCM to decrease the current threshold of activation and increase action potential firing (Figures 1E and S1C). Voltage-gated sodium channels (Na_V_s) mediate excitability of DRG neurons. Transcriptional increases in Na_V_s are induced by Schwann cells in culture^31^ and drive DRG neuron hyperexcitability following injury and in pain models^8, 42^. Thus, we hypothesized that Schwann cell-derived molecule(s) increase transcription of Na_V_s in DRG neurons to promote excitability. We used multiplexed fluorescence RNA in situ hybridization (RNAscope) to visualize and quantify expression of the major DRG Na_V_ subtypes (*Na_V_1.6*, *Na_V_1.7*, and *Na_V_1.8*)^8^ in DRG neurons cultured alone or treated with SCCM. Without SCCM, DRG neurons expressed few Na_V_ transcripts, consistent with their inability to fire action potential trains (Figures 1F–1I). SCCM treatment increased the expression of *Na_V_1.6*, *Na_V_1.7*, and *Na_V_1.8* channels by ∼2-4-fold (Figures 1F–1I). The effect of SCCM required molecules secreted by Schwann cells, as Schwann cell growth media alone was not sufficient to promote Na_V_ expression (Figure S1D). SCCM treatment did not alter expression of control housekeeping genes, DRG neuron soma size, or neurofilament staining (Figures S1E, S1F and S3), suggesting that the effect of SCCM on excitability is due to increased Na_V_ expression. Indeed, immunostaining experiments using knockout-validated antibodies demonstrated that SCCM treatment also increased Na_V_1.6, Na_V_1.7, and Na_V_1.8 protein levels by ∼1.5-2-fold, indicating that the increase in transcription levels results in a corresponding increase in protein concentrations (Figures 1J–1M; see also Figure S4B). Among other Na_V_ isoforms, Na_V_ 1.1, Na_V_ 1.2, Na_V_ 1.5, and Na_V_ 1.9 (which show low expression in DRG neurons^8^), SCCM only increased protein levels of Na_V_ 1.5 (Figure S4C). Overall, these results suggested that Schwann cells increase DRG neuron excitability by secreting molecule(s) that specifically promote expression of Na_V_1.6, Na_V_1.7, and Na_V_1.8.

### PGE_2_ is the excitability-inducing factor secreted by Schwann cells

Following our initial discoveries, we aimed to identify the Schwann cell-secreted molecule(s) that induce Na_V_ expression and excitability in DRG neurons. Size-based fractionation of SCCM with 5 kDa, 10 kDa, and 20 kDa molecular weight cutoff filters showed that the excitability- inducing effect remained in the 5 kDa flow-through fraction (Figure 2A; see Methods), implicating a small molecule or peptide. Mass spectrometry analysis of the 5 kDa media flow- through revealed several m/z signals, including a peak of 351.2182 ± 1.6 ppm. This value is consistent with the molecular weight of three prostaglandins (PGD_2_, PGE_2_, and PGI_2_), members of a family of endogenous lipid signaling molecules (Figure S4A).

**Figure 2.**
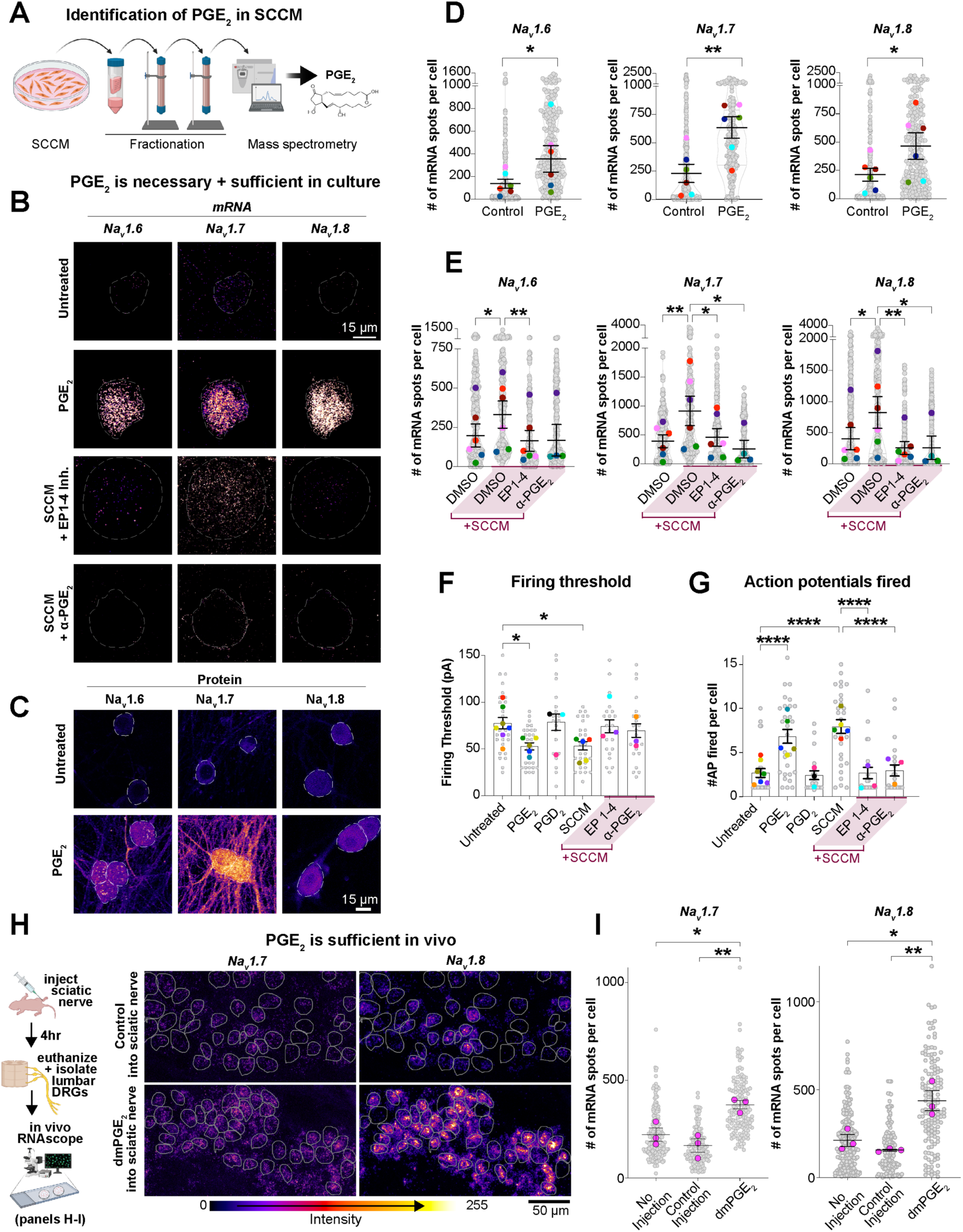
Identification of PGE_2_ as the Schwann cell-derived signal promoting neuronal Na_V_ expression and excitability. (**A**) Cartoon depicts size fractionation and mass spectrometry steps that identified PGE**_2_** as the excitability-inducing molecule in SCCM. (**B**) PGE_2_ treatment (1 μM, 16-28 hr) increased *Na_V_* expression in DRG neurons, similar to SCCM. *n* = 6 distinct biological replicates per treatment group. Bottom images: Incubation with neutralizing PGE_2_ antibody or PGE_2_ receptor (EP1-EP4) antagonists blocked the SCCM-induced transcriptional increase in *Na_V_*s (see also representative images for DMSO-treated DRG neurons in Figure S3E). *n* = 4-6 distinct biological replicates per treatment group. (**C**) PGE_2_ treatment enhanced Na_V_ protein levels in DRG neurons. Images show immunohistochemistry for indicated genes in single representative DRG neurons that were either untreated (top) or treated with PGE_2_ overnight (bottom) (see also quantification in Figure S4B). (**D** and **E**) Quantification of the RNAscope results shown in (B), using FishQuant. Controls were either untreated cells or cells treated with DMSO vehicle (solvent for PGE**_2_**). Mean ± SEM is shown for biological replicates. p values compare biological replicates in a paired t-test (D) or mixed-effect analysis (E). (**F** and **G**) DRG neurons treated with PGE_2_ (1 μM, 16-28 hr) fired significantly more action potentials at suprathreshold current injections and exhibited a decrease in the firing threshold, similar to SCCM. Incubation with neutralizing PGE_2_ antibody or PGE_2_ receptor (EP1–EP4) antagonists blocked the excitability-inducing effect of SCCM. DRG neurons treated with 1 μM PGD_2_, a constitutional isomer of PGE_2_, remained hypoexcitable (see also Figure S5). Mean ± SEM is shown for cells, p values compare cells in a one-way ANOVA and Tukey test. *n* = 20-29 total cells from 3–7 biological replicates. Untreated and SCCM conditions are replotted from Figs. 1C and 1D, as these experiments were done concurrently. (**H-I**) Injection of dmPGE_2_ into the P0 sciatic nerve increased *Na_V_1.7* and *Na_V_1.8* transcript levels in lumbar DRG neurons compared to controls (the third fluorescence channel was reserved for *Neun*; see Figure S7C). Results were quantified using FishQuant (I). Gray circles, *Na_V_* mRNA count in a single DRG neuron; colored circles, the average of cells in each biological replicate. Mean ± SEM is shown for biological replicates. p values compare biological replicates in cells in a one-way ANOVA and Tukey test. *n* = 3 mice per treatment group (at least 45 DRG neurons per mouse). *p<0.05, **p<0.01, ***p<0.001, ****p<0.0001.

Of the putative prostaglandins detected in SCCM, prostaglandin E_2_ (PGE_2_) is a signaling molecule with diverse functions in development, repair, and inflammation^43–48^. Intriguingly, PGE_2_ has been shown to affect several aspects of Na_V_ function in mature sensory neurons, including ion conduction, phosphorylation, trafficking, and expression^49–58^. This established role of PGE_2_ in neuronal excitability led us to hypothesize that PGE_2_ is the excitability-inducing molecule in Schwann cell-conditioned media. To test this hypothesis, we first treated immunopanned DRG neurons with PGE_2_ (1.0 μM, 16–28 hr). Similar to SCCM, PGE_2_ treatment specifically increased transcript and protein levels of Na_V_1.6, Na_V_1.7, and Na_V_1.8 (Figures 2B–2D, S4B and S4C), decreased action potential firing threshold (Figure 2F), and increased the number of action potentials fired (Figures 2G and S5). In marked contrast, prostaglandin D_2_ (PGD_2_, 1 μM), a structural isomer of PGE_2_, did not show this excitability-inducing effect (Figures 2F, 2G and S5).

Given the role of PGE_2_ in inflammation, we tested whether SCCM or PGE_2_ treatments altered inflammatory or metabolic pathways that could indirectly affect neural excitability. We used bulk RNA sequencing (RNAseq) analysis of control, SCCM-, or PGE_2_-treated DRG neurons and analyzed altered pathways using Ingenuity Pathway Analysis (IPA). IPA analysis indicated no statistically significant changes in neural inflammation or metabolism pathways in SCCM- or PGE_2_-treated neurons (Figure S6A). However, we detected significant increases in established PGE_2_ downstream pathways such as G beta gamma signaling and PKA signaling (Figures S6B–D). We also measured increases in the expression of several voltage-gated sodium channels (Figure S6E), consistent with our earlier experiments. Overall, these results suggested that SCCM and PGE_2_ treatments activate canonical PGE_2_ downstream signaling pathways to promote the excitability of DRG neurons—and that neuronal inflammation and/or changes to metabolism are not responsible for the observed effects.

We next investigated whether PGE_2_ is the essential component of SCCM that gives rise to DRG excitability. PGE_2_ mediates its effects through four related G-protein coupled receptor subtypes, EP1-4, all of which are expressed by DRG neurons^59–61^. First, we used a cocktail of EP1-4 receptor antagonists (see Methods) to block PGE_2_ signaling in DRG neurons. Second, we incubated SCCM with a specific PGE_2_-neutralizing antibody to sequester PGE_2_ molecules and inhibit their binding to EP receptors^62, 63^. Both approaches prevented SCCM from increasing Na_V_ expression (Figures 2B and 2E), lowering the firing threshold (Figures 2F and S5), and increasing the number of fired action potentials (Figure 2G and Figure S5). Treatment with individual EP inhibitors also reduced the excitability-inducing effect of SCCM (Figures S7A and S7B). These results established that PGE_2_ signaling is required for SCCM to promote Na_V_ expression and excitability.

To test whether PGE_2_ signaling also induces Na_V_ expression in DRG neurons in vivo, we injected 16,16-dimethyl PGE_2_ (dmPGE_2_)—a PGE_2_ derivative that is resistant to degradation by 15-hydroxy PGDH and thus has a prolonged circulating half-life in vivo^45, 64^—into the sciatic nerve where Schwann cells wrap and ensheath DRG axons (Figure 2H). *Na_V_1.7* and *Na_V_1.8* transcript levels are low in lumbar DRG neurons at birth (postnatal day 0; P0), but their expression increases through postnatal development^12^. Local dmPGE_2_ injection into sciatic nerves substantially upregulated *Na_V_1.7* and *Na_V_1.8* in lumbar DRG neurons compared to controls (∼2-3-fold induction, Figures 2H and 2I) without affecting *Neun* expression (a marker of mature DRG neurons; Figure S7C) or causing inflammation (Figure S8). Since EP receptors localize to peripheral DRG axons in vivo^65, 66^, these data suggested that Schwann cell-secreted PGE_2_ could be sensed locally by DRG axons to induce Na_V_ expression.

### Ptges3 is necessary for the synthesis of PGE_2_ by Schwann cells

Do Schwann cells make and secrete PGE_2_ in vivo? In cells, Cox1/2 enzymes convert arachidonic acid to a prostaglandin precursor molecule, PGH_2_. A PGE_2_ synthase subsequently catalyzes the conversion of PGH_2_ to PGE_2_ (Figure 3A)^67^. To examine whether Schwann cells express genes in the PGE_2_ synthesis pathway, we used RNAseq of Schwann cells acutely isolated from P18 rat sciatic nerve^68^. Analysis of the resulting Schwann cell transcriptome indicated that only one of four homologous PGE_2_ synthases^67^, Ptges3, was expressed at appreciable levels (Figure 3B). This finding is consistent with recent single-cell RNAseq analysis of the mouse sciatic nerve, which showed expression of Ptges3 in the entire Schwann cell lineage beginning embryonically in Schwann cell precursors (Figure S9A)^69^. We also confirmed expression of Ptges3 in Schwann cells in vivo in the sciatic nerve using RNAscope (Figure 3C).

**Figure 3.**
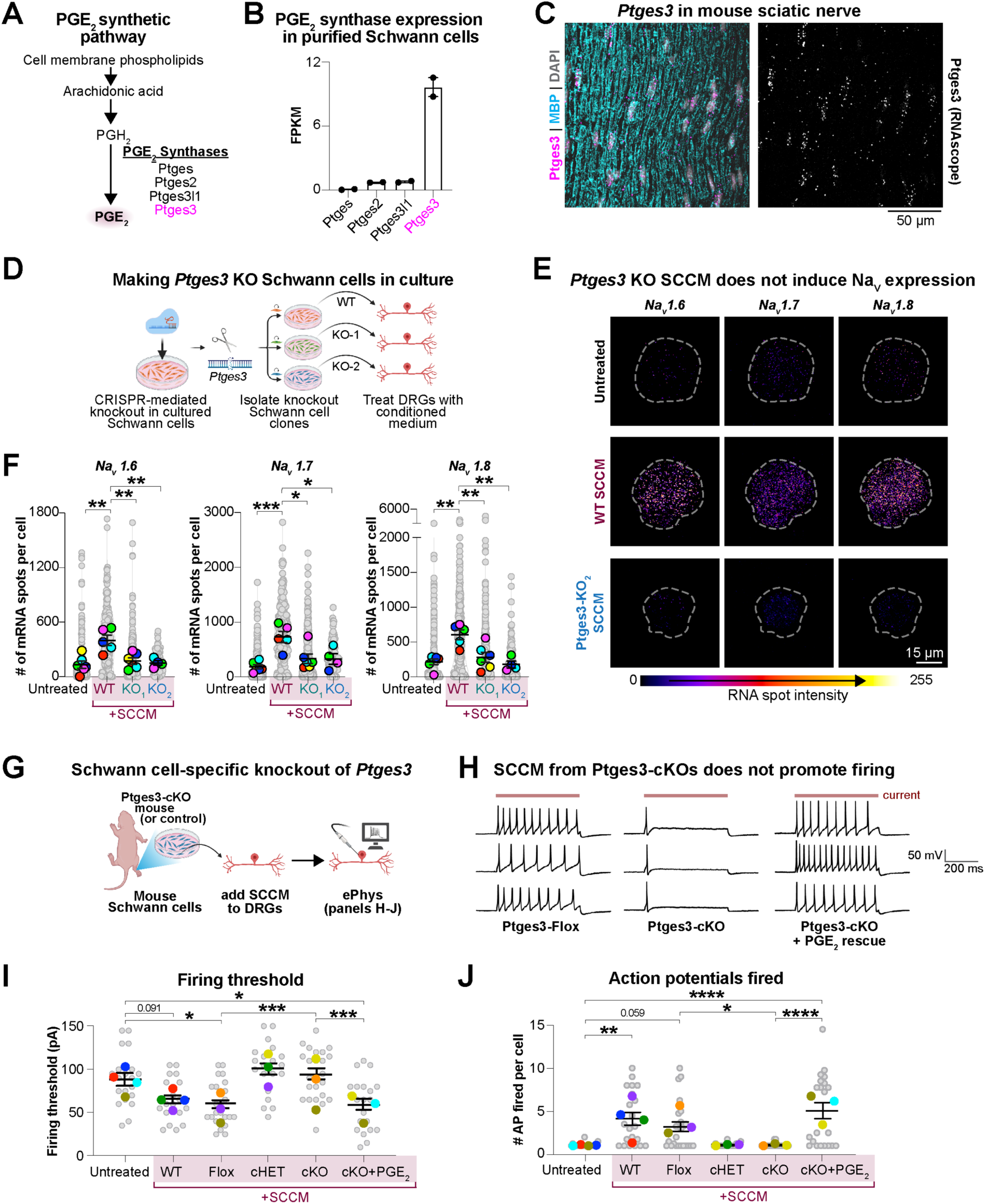
Ptges3 is essential for Schwann cells to produce PGE_2_ and promote neuronal excitability. (**A**) Diagram summarizes the PGE_2_ synthesis pathway. (**B**) RNAseq of Schwann cells purified from mature rat sciatic nerves showed expression of *Ptges3* (mRNA). *n* = 2 biological replicates, each replicate representing >10 pooled sciatic nerves. (**C**) RNAscope confirmed *Ptges3* expression in Schwann cells of the adult sciatic nerve, identified by nuclei aligned with Myelin Basic Protein (MBP)-expressing fibers. *n* = 3 mice (see also Figure S9A). (**D**) Generation of *Ptges3* knockout rat Schwann cell colonies. (**E**) SCCM from Ptges3-KO colonies failed to enhance DRG neuron *Na_V_* transcript levels. (**F**) Results were quantified using FishQuant. Gray circles, mRNA number of an individual DRG neuron for the indicated genes; colored circles, the average # of mRNAs per cell in each biological replicate. Mean ± SEM is shown for biological replicates, compared in a one-way ANOVA and Tukey test. *n* = 4–6 distinct biological replicates per group. (**G** to **J**) Mouse Schwann cells were acutely isolated from *Ptges3* conditional knockout mice (Ptges3-cKO: *Ptges3^fl/fl^;Dhh^CRE^*) or littermates (Ptges3-flox: *Ptges3^fl/fl^*and Ptges3-cHet: *Ptges3^fl/+^;Dhh^CRE/+^*). All SCCM was filtered through a 5 kDa filter before adding to DRG neurons. Representative traces (H) indicate activity over 500 ms with 150 pA of current injection. Adding PGE_2_ to cKO SCCM rescued the loss of excitability-inducing effect in cKO SCCM (see also Figure S11 and 12). Mean ± SEM is shown for cells, compared in a one-way ANOVA and Tukey test. *n* = 20–25 total cells from 3–4 distinct biological replicates per group. *p<0.05, **p<0.01, ***p<0.001, ****p<0.0001.

In principle, knocking out Ptges3 in Schwann cells would eliminate their ability to produce PGE_2_ and thus prevent their effect on DRG neuron excitability. We first tested this hypothesis in culture by using CRISPR/Cas9 genome editing to knock out *Ptges3* in Schwann cells immunopanned from rat sciatic nerves. We isolated single Schwann cells and grew them into genetically-identical colonies (Figure 3D). We identified two different Schwann cell colonies with biallelic deletion or insertion events (INDELs) predicted to cause complete loss of Ptges3 function (Figure S9B). We also isolated one colony with no INDELs in the *Ptges3* locus as a wild-type control to test for possible artifacts of prolonged culture or off-target mutations caused by CRISPR/Cas9. *Ptges3* knockout Schwann cells lost Ptges3 protein expression compared to wild-type control cells (Figures S9C–S9E).

We next tested whether SCCM collected from *Ptges3* knockout colonies increased the excitability of DRG neurons to the same extent as wild-type SCCM. As before (Figures 1F and 1I), SCCM collected from either acutely-isolated Schwann cells or the wild-type colony increased transcript levels of *Na_V_1.6*, *Na_V_1.7*, and *Na_V_1.8*. In contrast, SCCM from both *Ptges3* knockout colonies failed to induce Na_V_ expression (Figures 3E and 3F). Overall, these results established that Ptges3 is essential for Schwann cell production of PGE_2_ and provided further support that PGE_2_ signaling from Schwann cells promotes DRG neuron excitability.

### Schwann cell-secreted PGE_2_ promotes Na_V_ expression and DRG neuron excitability in vivo

Identifying *Ptges3* as an essential gene for Schwann cell synthesis of PGE_2_ enables studies to determine whether this putative glia-to-neuron signaling pathway regulates sensory neuron excitability in vivo. Global knockout of *Ptges3* causes perinatal lethality^70–72^; thus, we generated a conditional knockout mouse line for *Ptges3* to determine the role of Ptges3 and PGE_2_ synthesis specifically in Schwann cells. Previous studies showed that targeted disruption of exon 2 and 3 of the *Ptges3* locus leads to complete loss of function^71^. Using CRISPR/Cas9 genome editing, we targeted loxP sequences flanking exons 2 and 3 to enable Cre-mediated excision of the intervening DNA (Figure S10A). We confirmed the site-specific integration of both loxP sites and the absence of any silent mutations by sequencing. We crossed *Ptges3^fl/fl^* mice to *Dhh^Cre/+^* mice to conditionally knock out *Ptges3* in embryonic Schwann cell precursors^73, 74^, and we confirmed recombination in the *Ptges3* locus in Schwann cells purified from these mice (Figure S10B). As expected, Cre-mediated excision of exons 2 and 3 caused a significant decrease in *Ptges3* mRNA levels in the sciatic nerve (∼75% decrease by RNAscope; Figure S10C). *Ptges3* conditional knockout (“Ptges3-cKO”, genotype: *Dhh^Cre/+^; Ptges3^fl/fl^*) and conditional heterozygous (“Ptges3-cHet”, genotype: *Dhh^Cre/+^; Ptges3^fl/+^*) mice showed no gross abnormalities and had comparable weight to wild-type (“Ptges3-Flox”, genotype: *Ptges3^fl/fl^*) littermates (Figures S10D and S10E).

We first tested whether *Ptges3* conditional knockout in Schwann cells impairs their ability to increase the excitability of immunopanned DRG neurons in culture. Similar to wild-type rat SCCM (Figures 1B to 1I), conditioned media collected from Ptges3-Flox Schwann cells lowered the firing threshold of DRG neurons, increased their ability to fire multiple action potentials, and increased the transcript levels of *Na_V_1.6*, *Na_V_1.7*, and *Na_V_1.8* in DRG neurons (Figures 3G–3J, Figure S11, and Figure S12). In contrast, treatment with SCCM collected from Ptges3-cKO or Ptges3-cHet Schwann cells failed to promote this excitability-inducing effect: action potential firing threshold remained high, DRG neurons rarely fired more than one action potential, and Na_V_ transcript levels remained low (Figures 3G–3J, Figure S11, and Figure S12). Adding PGE_2_ to Ptges3-cKO SCCM rescued the loss of the excitability-inducing effect (Figure 3G–3J, Figure S11, and Figure S12). These results indicated that Ptges3-cKO cells lack the ability to increase the excitability of sensory neurons specifically due to reduced levels of secreted PGE_2_.

We next used RNAscope and immunohistochemistry to examine whether loss of Schwann cell-secreted PGE_2_ impairs the expression of Na_V_s in DRG neurons in vivo during development. Strikingly, transcript levels of *Na_V_1.7* and *Na_V_1.8* in Ptges3-cKO mice were less than half of those measured in control embryos at embryonic day 16 (E16) (Figure S13). At the day of birth (P0), expression levels of *Na_V_ 1.7* and *Na_V_1.8* remained lower than half of control expression levels (Figures 4A–4B, and Figure S14A). Protein levels of Na_V_1.7 and Na_V_1.8 were also reduced in Ptges3-cKO pups at P0 (Figure 4C–4D). We did not find significant changes in the expression levels of *Na_V_1.1*, *Na_V_1.2*, *Na_V_1.3,* or *Na_V_1.6* at E16 or P0, indicating that compensatory upregulation of these Na_V_ subtypes did not occur in cKO mice (Figures S13 and S14). Interestingly, expression of the neuron differentiation marker *Neun* was also significantly lower in Ptges3-cKO mice at E16 and P0 (Figures S13 and S14), raising the possibility that neuronal maturation is delayed in these animals (see also Figure 5).

**Figure 4.**
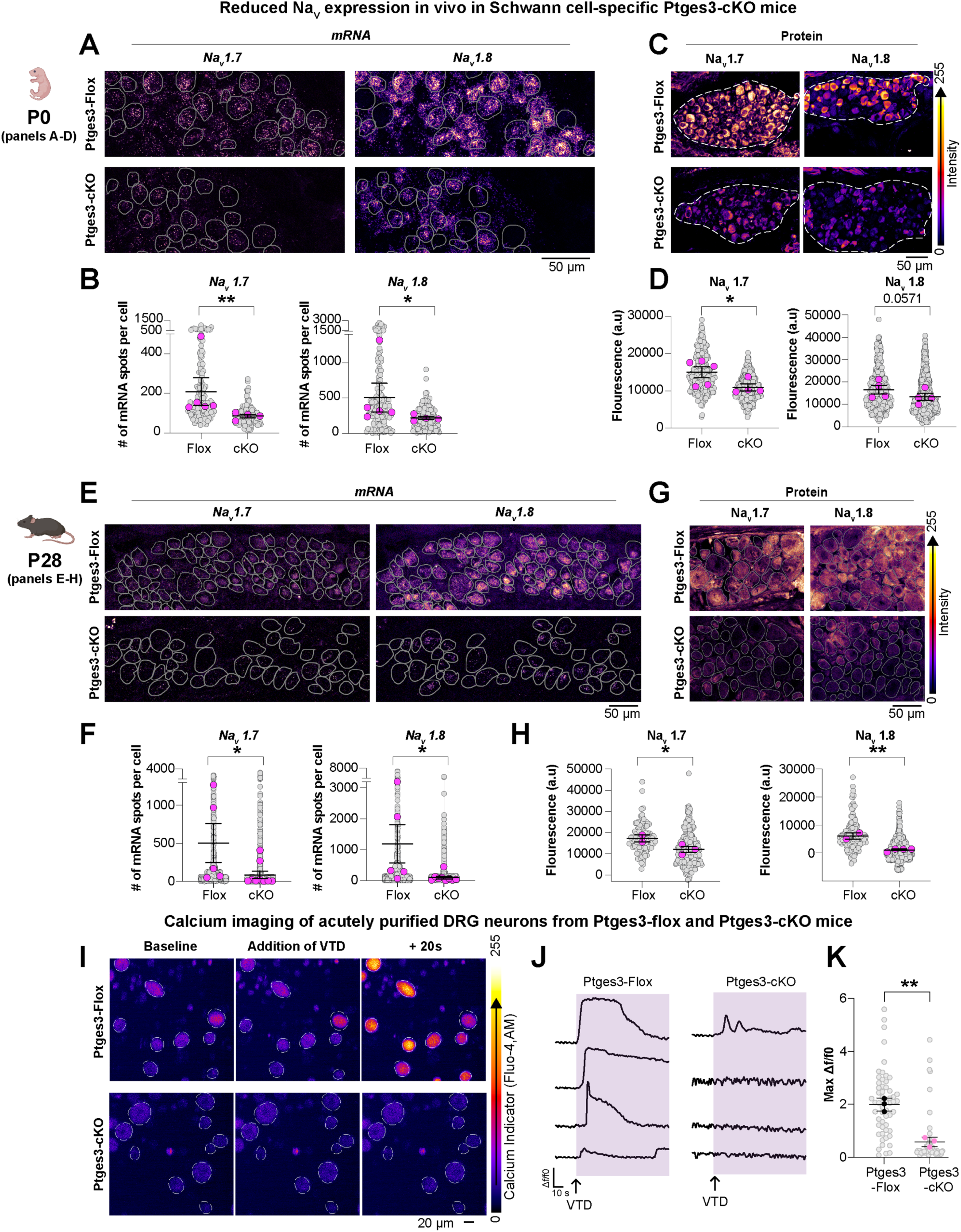
Schwann cell-secreted PGE_2_ promotes Na_V_ expression and DRG neuron excitability in vivo. (**A** to **H**) RNAscope and immunostaining of Na_v_ expression in vivo in Ptges3-Flox or Ptges3-cKO mice. Left panels: RNAscope images show expression of *Na_V_1.7* and *Na_V_1.8* transcripts in lumbar DRG neurons at P0 (A) and P28 (E) (see also figs. S13, S14, and S15 for E16 time point and supporting data). Number of *Na_V_1.7* and *Na_V_1.8* mRNAs per cell quantified using FishQuant from P0 (B) and P28 (F). *n* = 4-9 mice per group. Right panels: immunohistochemistry for Na_V_1.7 and Na_V_1.8 in lumbar DRG neurons at P0 (C) and P28 (G). Cellular fluorescence was quantified using Cellpose. Gray circles, mRNA count or fluorescence intensity in a DRG neuron; pink circles, the average of cells in each mouse. *n* = 2-5 mice per group. Mean ± SEM is shown for biological replicates (mice). p values compare biological replicates in Mann-Whitney test (B, D and F) or an unpaired t-test (H). (**I** to **K**) Calcium imaging of acutely purified DRG neurons from Ptges3-Flox or Ptges3-cKO mice. Images show fluorescent intensity of the calcium indicator (Fluo-4,AM) in DRG neurons acutely isolated (within 2 hours of purification) from Ptges3-Flox (top) or Ptges3-cKO (bottom) mice at baseline, during the addition of VTD and 20 seconds (s) after VTD addition (I). Representative neuron traces (J) and the maximum difference between the florescence after stimulation and during baseline (max delta F/F) (panel K) are shown. n = 3 distinct biological replicates (mice) per group, 8-26 cells per mouse. Mean ± SEM is shown for biological replicates (mice). p values compare biological replicates in an unpaired t-test. *p<0.05, **p<0.01, ***p<0.001, ****p<0.0001.

**Figure 5.**
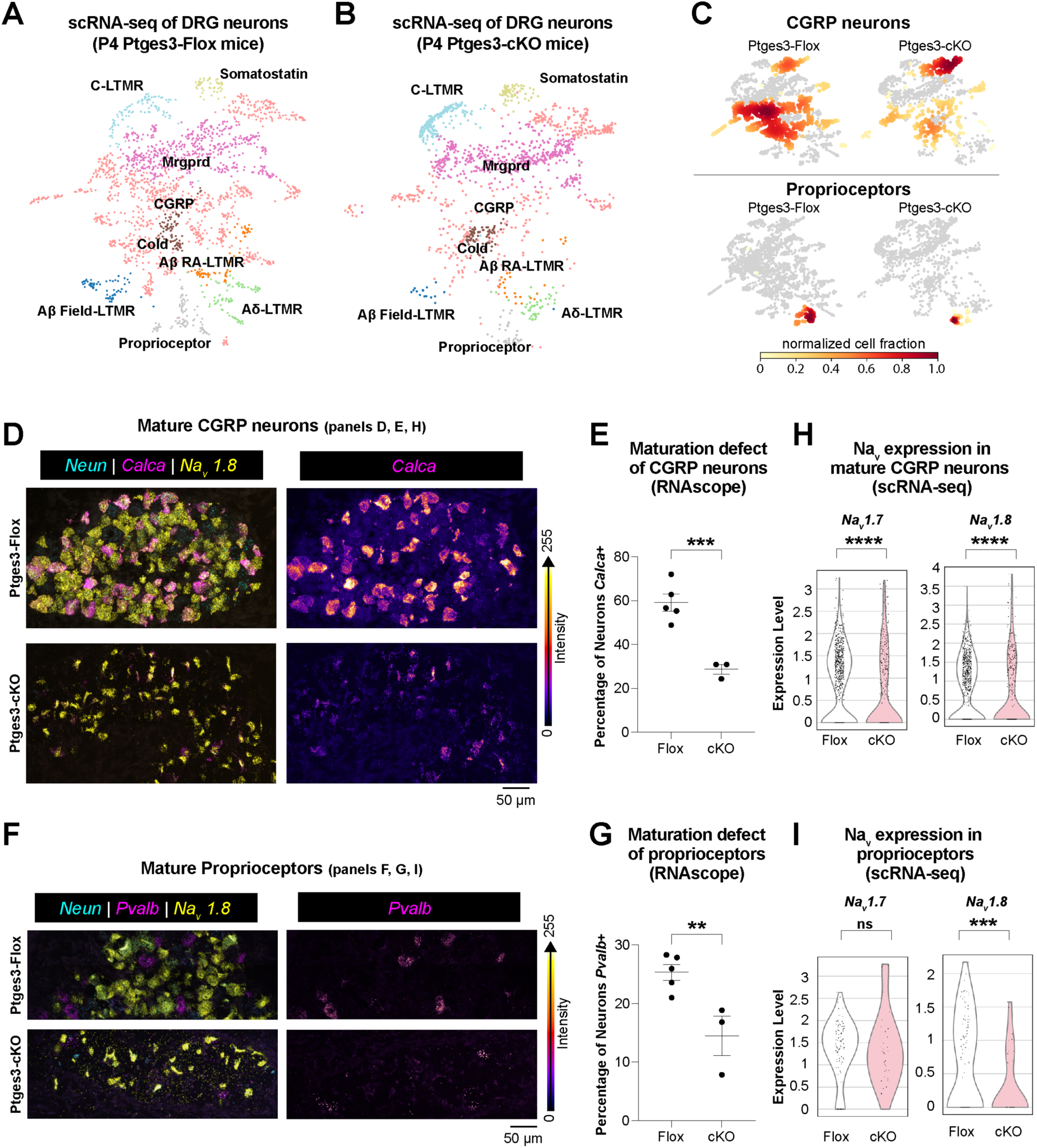
Schwann cells regulate the development and function of pain-sensing and proprioceptive DRG neuron subtypes. (**A** and **B**) UMAP visualization of DRG scRNA-seq data in Ptges3-Flox and Ptges3-cKO mice at P4. (**C**) Cell identity composition heatmaps of CGRP and proprioceptor DRG neuron subtype populations. Dark red indicates high relative cell density. (**D** to **G**) RNAscope shows expression of *Neun, Pvalb, or Calca,* and *Na_V_1.8* transcripts in lumbar DRG neurons at P4 to label proprioceptor or CGRP neurons, respectively. *Pvalb*+ or *Calca*+ neuron numbers are normalized to *Neun*+ cells to calculate the percentage of proprioceptor or CGRP DRG neurons respectively. *n* = 3-5 mice, p values compare biological replicates in an unpaired t-test. (**H** and **I**) Bar plots indicate expression of *Na_V_ 1.7* and *Na_V_1.8* in CGRP and proprioceptor DRG subpopulations, from scRNA-seq data. p values compare cells in an unpaired t-test. *p<0.05, **p<0.01, ***p<0.001, ****p<0.0001.

Does the impaired sodium channel expression we observed in early development persist postnatally? Expression of sodium channels in DRG neurons normally increase during postnatal development, reaching adult levels by P28^9, 12, 75^. However, similar to earlier developmental stages, P28 Ptges3-cKO mice had a ∼5-12-fold reduction in *Na_V_1.6*, *Na_V_1.7*, and *Na_V_1.8* transcript levels in lumbar DRG neurons compared to control littermates (Figures 4E, 4F, S15B and S15C). Protein levels of Na_V_1.7 and Na_V_1.8 were also 1.5- and 5-fold lower, respectively (Figures 4G and 4H). Consistent with impaired neuronal activity, the neuronal activity marker cFos was reduced ∼4-fold in lumbar DRG neurons in Ptges3-cKO mice compared to control mice (Figures S15C and S15D). Electron microscopy of sciatic nerves showed that DRG axonal development, Schwann cell myelination, and Remak bundle formation were ultrastructurally normal in Ptges3-cKO and Ptges3-cHet mice (Figure S16), indicating that reduced Na_V_ expression in DRG neurons is unlikely to be a secondary effect of gross axonal or myelin defects. Together, these results suggested that PGE_2_ secretion from Schwann cells is required in vivo for the normal expression of Na_V_ channels and, perhaps, more broadly for neuronal function.

To assess the functional consequences of Ptges3 conditional knockout on DRG neuron excitability during development, we acutely isolated lumbar DRG neurons from postnatal (P18/P19) Ptges3-Flox or Ptges3-cKO mice in the presence of transcriptional blockers (to prevent ex vivo gene expression changes induced by the isolation procedure^76^) and acutely measured their excitability within 2 hours of purification. Veratridine (VTD) is a voltage-gated sodium channel agonist that binds to open Na_V_s and prevents them from entering an inactivated state^77^. We therefore used VTD to activate DRG neurons acutely isolated from Ptges3-Flox or Ptges3-cKO mice and measured calcium responses as a readout of Na_V_-dependent neuronal activity. Addition of 5 μM VTD elicited rapid calcium responses in DRG neurons isolated from Ptges3-Flox controls (max f/f0= 1.99), whereas calcium responses were minimal in DRG neurons isolated from Ptges3-cKO mice (max f/f0= 0.58, Figures 4I to 4K). This result showed that Na_V_-induced neuronal responses are severely dampened in Ptges3-cKO mice and that Schwann cell-secreted PGE_2_ is crucial for development of normal neuronal activity in DRG neurons in vivo.

### Schwann cell-secreted PGE_2_ promotes the development and function of pain-sensing and proprioceptive DRG neuron subtypes

Distinct DRG neuron subtypes innervate peripheral tissues and transduce mechanical, thermal, and chemical signals to the CNS^36, 78–80^. During embryonic development, DRG neurons emerge in an unspecialized state and are thought to require extrinsic cues from the ganglia and axonal environment to mature into specific subtypes^13, 14^. Our finding that Schwann cell-secreted PGE_2_ promotes the development of sensory neurons into mature and excitable cells prompted us to test if all DRG subtypes and sensory modalities are equally impaired by the loss of Schwann cell-secreted PGE_2_.

First, we compared DRG neurons from Ptges3-Flox and Ptges3-cKO mice using single-cell RNA sequencing (scRNA-seq) of dorsal root ganglia from all axial levels at postnatal day 4, when DRG subtypes have normally been fully specified^13^. Using principal component analysis (PCA) and uniform manifold approximation and projection (UMAP), we clustered DRG neurons from Ptges3-Flox mice based on the similarity of gene expression profiles (Figure 5A). We then classified each cluster as a subtype based on marker genes described in previous studies^13, 36, 80^ (Figures 5A, 5B, S17A, and S17B). Comparison of the scRNA-seq transcriptome of DRG neurons isolated from Ptges3-cKO mice by projecting onto the UMAP of Ptges3-Flox controls revealed that there were fewer mature CGRP and proprioceptor DRG neurons in Ptges3-cKO mice (Figure 5C). We also measured a ∼30% decrease in the number of NeuN+ cells in the lumbar DRG neurons of Ptges3-cKO mice at P4 (Figure S17D; see also reduced *Neun* expression in cKO mice in Figures S13B and S14A). RNAscope analysis of *Calca+* (marks CGRP neurons) and *Pvalb+* (marks proprioceptors) DRG neurons confirmed a 54% and 40% decrease in the number of neurons expressing these markers, respectively (Figure 5D–5G). Examination of the transcriptomes of CGRP and proprioceptive neurons in Ptges3-cKO mice revealed that these subtypes also expressed fewer *Na_V_1.7* and *Na_V_1.8* transcripts (Figures 5H and 5I) and showed reduced expression of mature neuronal markers (such as *Tubb2b* and *Prph*, also see Figure S18). The reduction in numbers of mature neurons can be attributed to a delay in their maturation rather than cell death, as we observed no cell death in lumbar DRG neurons in Ptges3-cKo mice at P0 using the TUNEL assay (Figure S19A) and no upregulation of apoptosis-related genes via transcriptomic analysis (Figure S19B). Moreover, by P28 there was no longer a difference in *Neun* expression (Figure S15D) or in the number of Neun+ cells between genotypes (Figure S19C), consistent with a maturation delay rather than neuronal death. Overall, these results confirmed that functional maturation of CGRP and proprioceptor DRG neuron subtypes is impaired in Ptges3-cKO mice.

Next, we recorded compound action potentials (CAPs) from the sciatic nerve of adult Ptges3-Flox and Ptges3-cKO mice ex vivo to directly measure nerve excitability. CAP recordings allowed the simultaneous measurement of excitability and conduction velocities of both Aα/β and Aδ fibers (containing axons from motor neurons, proprioceptors and mechanoceptors) and C fibers (containing axons from nociceptors). Consistent with our scRNA-seq results (Figure S17C), we measured normal nerve excitability in Aα/β and Aδ fibers, which are enriched in DRG subtypes that were not strongly perturbed in Ptges3-cKO mice (Figure S20). In contrast, C-fibers exhibited a higher threshold stimulus voltage (voltage at which a CAP is detectable), indicating that C fibers in Ptges3-cKO nerves were hypoexcitable (Figures 6A and 6B). We measured no differences in peak amplitude or latency of the largest C-fiber CAP recorded for Ptges3-Flox and Ptges3-cKO nerves, evidence that the number and overall health of C fibers were not impaired by loss of Ptges3 in Schwann cells (Figures 6C and 6D). Together, these data showed that a major phenotype in Ptges3-cKO mice was reduced excitability of small diameter, unmyelinated nociceptive neurons. We were unable to resolve proprioceptor-specific signals from the other Aα/β fibers in our recordings.

**Figure 6.**
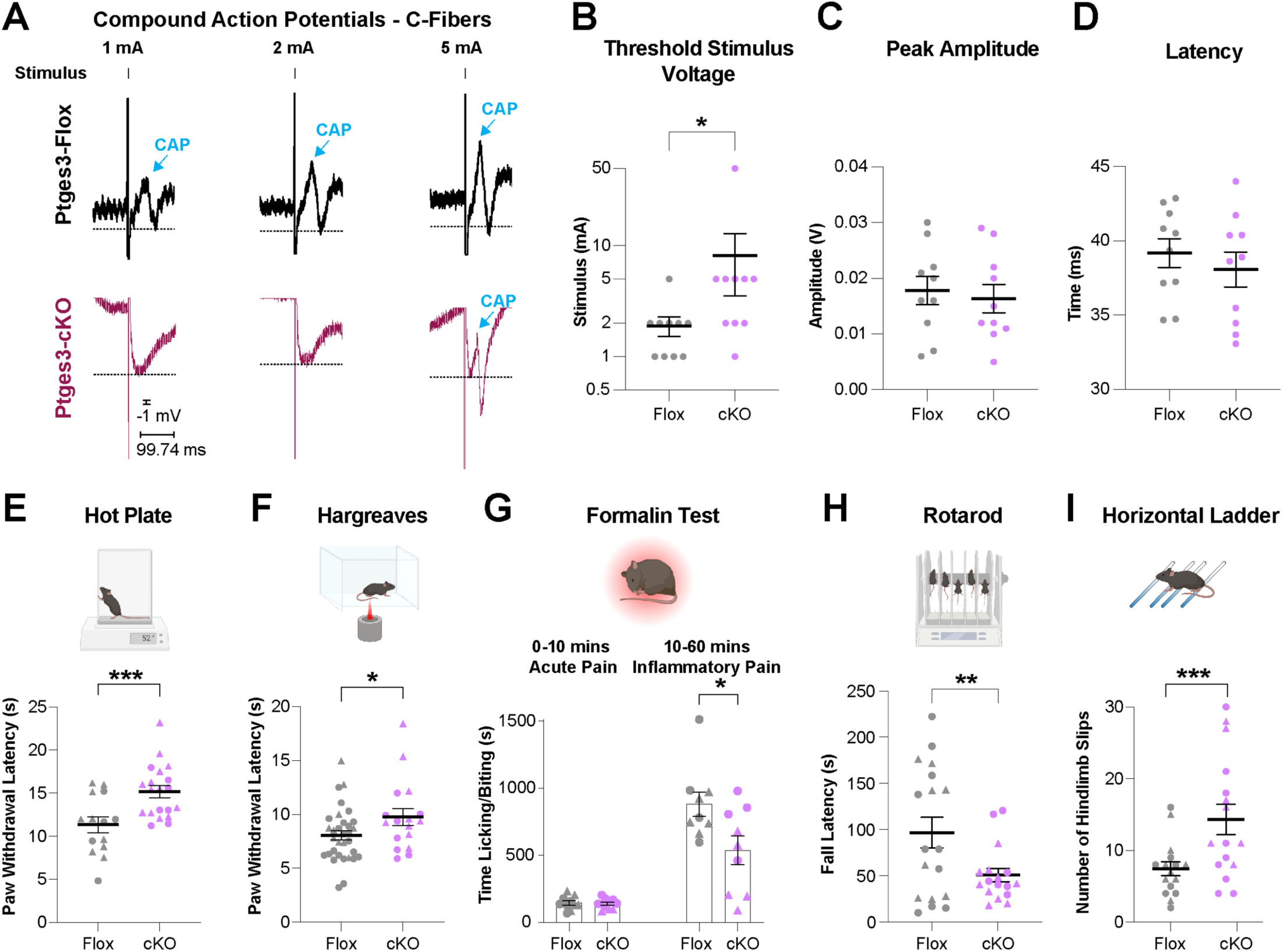
Schwann cell-secreted PGE_2_ is required for normal nerve excitability and sensory function. (**A**) C fiber compound action potentials in Ptges3-Flox and Ptges3-cKO sciatic nerve in response to indicated stimulus applications. (**B**) Threshold stimulus voltage of C fibers is decreased in Ptges3-cKO sciatic nerve, consistent with reduced excitability. y axis is plotted in log2 scale. (**C** and **D**) Peak amplitude and the latency of the largest CAP did not reveal significant changes between Ptges3-cKO and Ptges3-Flox animals. n=10 sciatic nerves from 5-7 male mice for each group. p values compare sciatic nerves in Mann-Whitney test. (**E** and **F**) Hot plate and Hargreaves tests indicate that paw withdrawal latencies in response to noxious heat was significantly lengthened in Ptges3-cKO mice. *n* = 14, 20 mice (hot plate); 32, 17 mice (Hargreaves). Control littermates in (F) were either *Ptges3^fl/fl^* or *Ptges3^fl/+^* mice. (**G**) Pain response following 1% PFA injection into the hind paw indicated the typical biphasic response. The first phase (acute pain) was unaffected, but the second phase (inflammatory pain) was reduced by ∼40% in Ptges3-cKO mice. *n* = 9-11 mice (also see Figure S21). (**H**) Rotarod assay (32 RPM, constant speed) indicates decreased fall latency in Ptges3-cKO mice. *n* = 18, 17 mice. (**I**) Precise foot placement is severely impacted in Ptges3-cKO mice recorded in the horizontal ladder test with unevenly placed rungs. *n* = 16 mice for each group. p values compare biological replicates (triangles-females, circles-males) in an unpaired t-test. *p<0.05, **p<0. 01, ***p<0.001, ****p<0.0001.

Are CGRP and proprioceptor DRG neuron functions impaired in Ptges3-cKO mice? Selective ablation of CGRP neurons in mice leads to a profound reduction in thermoreception, whereas cold and mechanical sensitivity remain unaffected^81^. In addition, loss of Na_V_1.7 and Na_V_1.8 channels also leads to impaired thermal sensitivity in mice^82–84^. These findings prompted us to test heat responsivity in mice lacking Ptges3 in Schwann cells. Similar to mice lacking CGRP nociceptors or Na_V_1.7 and Na_V_1.8 channels, Ptges3-cKO mice displayed impaired thermal sensitivity as measured by a ∼35% (∼4 s) increase in paw withdrawal latencies in the hot plate test (Figure 6E). Ptges3-cKO mice also displayed a 20% increase in paw withdrawal latency to localized infrared heat application in the Hargreaves test (Figure 6F). The magnitudes of these effects were comparable to withdrawal latencies recorded in nociceptor-specific knockout of Na_V_1.7 or global knockout of Na_V_1.8 ^82–84^. Similar to single and double Na_V_ knockout mice^82–84^ or genetic ablation of CGRP neurons^81^, Ptges3-cKO mice had normal mechanical sensitivity to Von Frey fibers (Figure S21A).

CGRP neurons^81^ and Na_V_1.7 and Na_V_1.8^82–84^ also contribute to inflammatory pain. To test whether inflammatory pain is impaired in Ptges3-cKO mice, we injected formalin into the hind paw, which induces a characteristic biphasic pain response with acute pain (0-10 min) and inflammatory pain (10-60 min). Acute pain response was unaffected in Ptges3-cKO mice, whereas inflammatory pain was reduced by ∼40% (Figure 6G). Interestingly, the response to inflammatory pain was more pronounced in females (Figures S21B and S21C). These results indicated that Schwann cell-secreted PGE_2_ is required for the normal development of fully functional CGRP neurons in vivo.

Finally, we investigated the functional consequences of the improper maturation of proprioceptive neurons (as predicted by our scRNA-seq and histology data) in Ptges3-cKO mice. Catwalk gait analysis revealed that Ptges3-cKO mice had significant defects in several locomotion parameters, including a dramatic loss of interpaw coordination in females (Figures S22A, S22C, and S23), decreased frequency of normal step patterns in males (Figure S22B, S22D, and S23), and general effects on gait (Figure S23). Moreover, Ptges3-cKO mice showed a decreased fall latency in the rotarod test (Figure 6H) but did not exhibit any signs of gross motor defects (Figure S21A). Together, these results were consistent with proprioceptive deficits in both sexes. To further test loss of proprioceptive sensory function in Ptges3-cKO mice, we performed a skilled motor behavior test and examined walking on a horizontal ladder test with nonuniformly placed rungs^85, 86^. Ptges3-cKO mice exhibited a 92% increased foot drop error rate compared to control littermates, measured as foot slips between the rungs (Figure 6I, also see Figure S23C). These results phenocopied mice lacking proprioceptive sensory feedback^85, 87^, suggesting that proprioceptive functions are impaired in Ptges3-cKO mice. Overall, our data support a model in which Schwann cell-secreted PGE_2_ is required for specific subtypes of DRG neurons to become fully excitable and that this glia-neuron signaling interaction regulates sensory neuron function.

## DISCUSSION

How do neurons first become excitable? This remains a fundamental question in development and neurobiology and has broad implications for understanding how dysregulation of excitability emerges in many diseases of the nervous system. Here, we discovered that somatosensory neurons require a secreted signal from Schwann cells to express voltage-gated sodium channels (Na_V_s) and achieve normal excitability. We identified the Schwann cell-derived signal as PGE_2_. Inactivating PGE_2_ synthesis specifically in Schwann cells (Ptges3-cKO) reduced expression of several key Na_V_s in DRG neurons and reduced neuronal activity. Single-cell RNA-seq, histology, and behavioral studies of Ptges3-cKO mice revealed that this signaling pathway selectively drives the maturation of pain-sensing and proprioceptive neuron subtypes. Together, our studies identify a previously undescribed role for PGE_2_ in promoting neural excitability during development (Figure 7). Although glia have been found to indirectly modulate the excitability of mature neurons (e.g., by strengthening synaptic connections or ion buffering^18–24^), our discovery establishes that glia also developmentally regulate intrinsic neuronal excitability.

**Figure 7.**
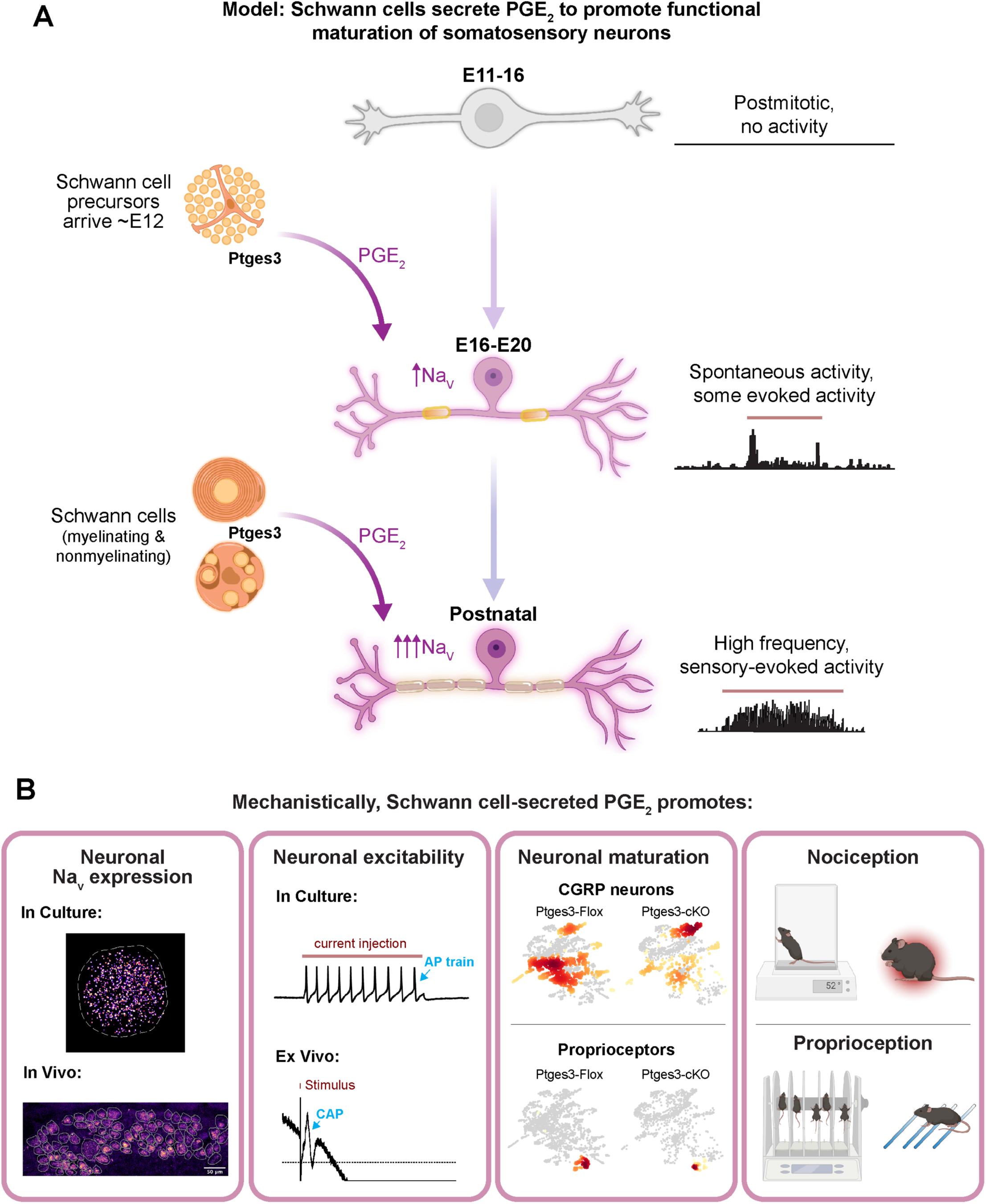
Model figure: Schwann cells promote sensory neuron excitability and normal sensory behavior via secreted PGE_2_. (**A**) During sensory neuron development, DRG neurons coalescence into ganglia at around E11 and remain inactive until E16. Schwann cell precursors arrive in the ganglia at around E12 and begin to differentiate into myelinating and Remak Schwann cells. Our model proposes that Schwann cell precursors and mature Schwann cells secrete PGE_2_ to initiate excitability and electrical activity in DRG neurons, allowing them to mature (graphics adapted from previous studies^9, 75^). (**B**) Mechanistically, our studies reveal that Schwann cell-secreted PGE_2_ promotes excitability of DRG neurons during development by inducing expression of neural Na_V_s (left box) to increase neuronal excitability (center-left), leading to maturation of nociceptors and proprioceptors (center-right) and, accordingly, for the normal development of related sensory modalities (right).

In the adult nervous system, a well-known role of PGE_2_ is as an inflammatory mediator that maladaptively promotes sensory neuron hyperexcitability following injury and in pain or disease states^8, 56, 88^. However, PGE_2_ is also known to have a diverse set of non-inflammatory functions across a variety of cell types, including essential signaling roles in embryonic development, regulating blood flow, and promoting tissue regeneration^43–48^. Complementary to these prior studies, our work shows that PGE_2_ secreted from Schwann cells has an important signaling function in the developing, healthy PNS—distinct from its role in neuroinflammation in adults. As an inflammatory mediatory, PGE_2_ primarily regulates neuronal excitability through Na_V_ phosphorylation and by promoting Na_V_ trafficking^56^ rather than regulating gene expression. Here, we identify a more fundamental, noninflammatory mechanism by which PGE_2_ regulates neuronal excitability—by inducing the basal transcription of Na_V_s.

Our findings raise several questions. First, what is the mechanism by which Schwann cell-derived PGE_2_ promotes excitability in somatosensory neurons? PGE_2_ is known to function through activation of its G protein-coupled receptors EP1-EP4, all four of which are expressed by embryonic and mature DRG neurons^59–61^. Activation of either EP2 or EP4 receptors has been shown to elevate intracellular levels of the second messenger cAMP in DRG neurons^89, 90^. cAMP, in turn, stimulates survival of embryonic DRG neurons^91^ and can induce activation of the cAMP response element binding protein (CREB), which regulates transcription of numerous target genes essential for neuronal growth, survival, and differentiation^92^. Of note, elevating cAMP signaling (e.g., by forskolin or db-cAMP) is a common strategy for promoting neuronal survival in defined medium and for inducing neuronal differentiation from iPSCs^27, 28, 93–95^. Future work will test whether PGE_2_ regulates DRG neuron maturation through cAMP and CREB signaling as predicted by this model.

Our single-cell RNA-seq experiments revealed that loss of PGE_2_ signaling from Schwann cells broadly affects gene expression in developing DRG neurons beyond Na_V_s. These data also revealed that PGE_2_ is required for functional maturation of select subtypes of DRG neurons in vivo. How does transcriptional regulation of Na_V_s by PGE_2_ relate to these more widespread effects on neuronal differentiation? One possibility is that the neuronal differentiation defects we observed in Ptges3-cKO mice are a direct consequence of hypoexcitability. Neuronal excitability and activity have long been known to influence the differentiation of many types of neurons, including DRG neurons^96–100^. For example, using potassium chloride to mimic neuronal activity is sufficient to induce progenitor cells to become responsive to neurotrophins^96^ or mature into DRG neurons^97^. Our ability to rapidly purify embryonic DRG neurons away from glia—and thus, to have control over their ability to express Na_V_s—may enable us to resolve whether the broader transcriptional changes we observed in Ptges3-cKO mice are a downstream consequence of reduced neuronal excitability. Moreover, our development of a Ptges3-Flox mouse will allow us to determine whether continued PGE_2_ secretion from Schwann cells is required to maintain Na_V_ expression and/or DRG identity in adults.

How broadly throughout the nervous system does this glial-neuron signaling pathway extend? For example, does glial-derived PGE_2_ also regulate normal neuronal excitability in the CNS? PGE_2_ has been implicated in other facets of CNS neurodevelopment, including neural proliferation, synaptogenesis, and synaptic plasticity^88, 101–103^. Enzymes for PGE_2_ synthesis including Ptges3 are expressed by oligodendrocytes and other CNS glia in both mice and humans^104, 105^. Whether glial-derived PGE_2_ directly initiates or modulates neuronal excitability in the healthy or diseased CNS remains an open question for future studies.

In summary, our findings establish a novel function of glial cells for promoting the development of neurons into excitable cells. This discovery opens up new research directions to explore the influence and ramifications of glial PGE_2_ signaling on neural development, function, plasticity, and disease.

## ACKNOWLEDGMENTS

We thank the members of the Zuchero and Du Bois laboratories for feedback and support; Nirao Shah, Michelle Monje, and Ivan Soltesz for their critical reading of the manuscript; Dies Meijer and Kelly Monk for DhhCre/+ mice; Livia Eberlin for support in mass spectrometry experiments; John J. Perrino and Ibanri Phanwar-Wood at the Stanford University Cell Sciences Imaging Core Facility for assistance with electron microscopy; the Stanford Functional Genomic Facility, the Stanford Behavioral and Functional Neuroscience Laboratory, and the Stanford Transgenic, Knockout and Tumor model Center; and Ben A. Barres for mentorship and support in the early stages of this project. Figure schematics were created with BioRender.com and Adobe Illustrator. We gratefully acknowledge support from The Shurl and Kay Curci Foundation (JBZ), The McKnight Endowment Fund for Neuroscience McKnight Scholar Award (JBZ), National Institutes of Health grant R01GM117263-05 (JD), National Institutes of Health grant R35GM137906 (VLT), Stanford ChEM-H Postdocs at the Interface seed grant (HK and EEG), Walter V. and Idun Berry Postdoctoral Fellowship Program (HK), Stanford Dean’s Fellowship (HK), Stanford Jump Start Award (HK), Stanford ChEM-H Chemistry/Biology Interface Predoctoral Training Grant (PDE), National Institutes of Health training grant T32GM120007 (PDE), Center for Molecular Analysis and Design at Stanford University (PDE), Stanford Wu Tsai Neurosciences Institute Interdisciplinary Scholar Award (EEG), and the Stanford Graduate Fellowship in Science and Engineering (MI). Research reported in this publication was supported by the National Institute on Drug Abuse of the National Institutes of Health under Award Number T32DA035165 (LJD). The content is solely the responsibility of the authors and does not necessarily represent the official views of the National Institutes of Health. Electron microscopy studies were supported, in part, by ARRA Award Number 1S10RR026780-01 from the National Center for Research Resources (NCRR); their contents are solely the responsibility of the authors and do not necessarily represent the official views of the NCRR or the National Institutes of Health. The Stanford Behavioral and Functional Neuroscience Laboratory for behavioral testing was supported by the NIH S10 Shared Instrumentation for Animal Research grant 1S10OD030452-01. The Stanford Transgenic, Knockout and Tumor model Center was partially supported by National Institutes of Health grant P30CA124435.

## AUTHOR CONTRIBUTIONS

HK, PDE, AT, JD, and JBZ conceptualized the project. HK, PDE, AT, LJD, MQD, HZ, MH, AA, MM, MS, TAA, VLT, JD, and JBZ developed methodologies for experiments. HK, PDE, AT, LJD, MQD, NA, NLS, ABL, SAS, EEG, KVT, AV, AKY, AEM, MI, MH, AA, MM, TAA, and JBZ conducted experiments. HK, PDE, AT, MH, AA, MM, and JBZ contributed to data visualization. HK, LJD, EEG, JD, MM, MS, VLT, and JBZ acquired funding for the project. HK, PDE, JD, and JBZ wrote and edited the manuscript; all authors reviewed the manuscript. HK, HZ, MS, VLT, JD, and JBZ supervised and mentored contributors. JD and JBZ oversaw all aspects of the work.

## Data and materials availability

The data that support the findings of this study and step-by-step protocols are available from the lead author or first authors upon request. Raw fastq files resulting from the bulk RNA-seq studies were deposited in the National Center for Biotechnology Information (NCBI) Gene Expression Omnibus (GEO, accession number GSE177037). Single-cell RNA-seq data will be deposited at NCBI GEO prior to publication.

## SUPPLEMENTAL INFORMATION

Supplemental Information document includes the following:

- Methods
- Supplemental References
- Supplemental Figures S1—S23

## Supplemental Materials for Schwann cells promote the development of sensory neuron excitability

### Methods

#### Animals and ethics statement

All procedures involving animals were approved by the Institutional Administrative Panel on Laboratory Animal Care (APLAC) of Stanford University and followed the National Institutes of Health guidelines. Animals were group-housed in a Stanford University animal facility with a 12:12 hour light/dark cycle and with free access to water and food. All mice received regular monitoring from veterinary and animal care staff and were not involved in prior procedures or testing. *Dhh^CRE/+^*mice were obtained from Dr. Kelly Monk (OHSU/Vollum) and Dr. Dies Meijer (The University of Edinburgh). *Ptges3^fl/fl^* mice were generated at the Stanford University Transgenic, Knockout, and Tumor model Center and maintained by crossing to C57BL/6 mice. Sprague-Dawley rats and C57BL/6 mice were from Charles River Laboratories. Both male and female mice were studied for all in vivo experiments, including behavioral assays, and we kept track of sexes for all in vivo experiments. For cell culture studies, embryos or sciatic nerves of both sexes were pooled in order to obtain sufficient cell numbers.

#### Purification of primary embryonic rat dorsal root ganglion (DRG) neurons

DRG neuron isolation was performed following a method adapted from our previous report ^1, 2^. DRGs were dissected from E15 embryos using timed pregnant Sprague-Dawley rats. 10–18 embryos were pooled for each experiment, and each pregnant rat was considered one biological replicate. Following the enzymatic digestion of DRGs, dissociated cells were passed serially over a BSL-1 plate (to remove blood and endothelial cells) and a CD9 plate (to remove glial cells). This protocol yields a single-cell suspension of approximately 500,000 DRGs devoid of contaminating glia, blood cells, and fibroblasts. Additional details of the DRG isolation protocol are indicated below.

##### Preparation of coverslips

1 mg/mL of poly-D-lysine hydrobromide stock (Sigma-Aldrich, P6407) was diluted 100-fold in sterile ddH_2_O and added to coverslips for 30 minutes at room temperature. Coverslips were rinsed with sterile ddH_2_O three times and allowed to dry off completely. Coverslips were coated with 1 mg/mL laminin (R&D Systems, 3400-010-02) diluted 1:200 in Neurobasal media (Thermo Fisher Scientific, 21103049) at 37 °C for 4 to 24 hours. Laminin was removed from the plates immediately before plating neurons, and extra care was taken during plating to prevent the coverslips from drying.

##### Preparation of immunopanning plates

*BSL-1 plate*: One 15-cm petri dish was coated with 40 μL of 2 mg/mL BSL-1 (Vector Labs, L-1100) diluted in 20 mL PBS overnight at 4 °C. The plate was rinsed three times with sterile PBS and blocked with 9 mL of 0.2% BSA (Sigma-Aldrich, A-8806) at room temperature for at least 2 hours before using. *CD9 plate*: One 15-cm petri dish was coated with 90 μL of goat-anti-mouse IgG+ IgM (H+L) secondary antibody (Jackson ImmunoResearch, 115-005-044) diluted in 20 mL of Tris-HCl (pH 9.5) overnight at 4 °C. The plate was rinsed three times with sterile PBS and coated with anti-rat CD9 antibody (Thermo Fisher Scientific, BDB551808) diluted 1:200 in 0.2% BSA.

##### Dissections

One or two pregnant Sprague-Dawley rats with E15 embryos were euthanized by CO_2_ inhalation. The placentas were dissected out and placed in Leibovitz’s L15 dissection medium (Thermo Fisher Scientific, 11415114) supplemented with 10% FCS (Thermo Fisher Scientific, A3160401) and penicillin-streptomycin solution (1:100, Sigma-Aldrich, P0781). Embryos were removed from the amniotic sacs. Following decapitation, all organs and the spinal column on the ventral side of the embryos were removed until the spinal cord was entirely exposed along the anteroposterior axis. Next, the skin above the spinal cord was peeled off with two forceps, revealing DRGs lateral to the spinal cord. DRG (intact ganglia) were then cut out at their bases using small scissors.

##### Immunopanning

DRG immunopanning protocol was performed as previously described ^1, 2^. Briefly, intact DRGs were enzymatically dissociated by Papain+DNase digestion and serial trituration, then filtered through a 20 μm Nitex filter to obtain a single-cell suspension. Dissociated cells were passed serially over a BSL-1 plate (to remove blood and endothelial cells) and a CD9 plate (to remove glial cells).

##### Plating and culturing

Isolated DRG neurons were resuspended at 1000 cells/μL. DRG neurons were plated in densities ranging from 3,000–5,000 cells per coverslip for 5–10 minutes to allow for proper adherence. Coverslips were then immersed in defined DRG base medium ^1, 2^ supplemented with 100 ng/mL NGF (Neuromics, GT15057), 50 ng/mL BDNF (PeproTech, 450-02), and 1 ng/mL NT3 (PeproTech, 450-03). 20 mL of DRG base media was prepared with the following reagents: 9 mL Dulbecco’s Modified Eagle Medium (Gibco, 11960069), 9 mL Neurobasal Medium (Sigma, I2643), 200 μL insulin (Sigma, I2643), 200 μL penicillin-streptomycin (Gibco, P0781), 200 μL SATO solution (Bovine Serum Albumin [Sigma, A416], apo-transferrin human [Sigma, T1147], putrescine dihydrochloride (Sigma, P5780), progesterone (Sigma, P8783), sodium selenate [Sigma, S5261]), 400 μL NA-21 max [R&D systems, AR008], 200 μL 4 µg/mL 3,3′,5-triiodo-L-thyronine sodium salt [Sigma, T6397], 200 μL 200 mM L-glutamine [Gibco, 25030081], and 20 μL 5mg/mL N-acetyl-L-cysteine [Sigma, A8199]). The media was sterile filtered using a DMEM-rinsed 0.22 μm filter unit. Half-volume media change was done every 2–3 days. Forskolin was omitted from DRG neuron cultures since we determined that it is not required for neuronal survival or health and due to its possible effects on neuronal physiology through elevating cAMP levels (see Discussion).

#### Purification of rat or mouse Schwann cells

Sciatic nerves from one litter of P2 rats were dissected for purification of Schwann cells by immunopanning as described previously ^3^. For mouse Schwann cell isolation, sciatic nerves were isolated from P12 or P28 mice that had been previously genotyped by Transnetyx. Schwann cell isolation steps are outlined briefly below:

##### Dissociation

Sciatic nerves were incubated in a collagenase/dispase cocktail composed of 1 mg/mL collagenase (Roche, 10103578001) and 2.9 mg/mL dispase (Roche, 101103578001) in 1X HEPES buffered saline solution (Sigma-Aldrich, 51558) at 37 °C for 1.5 hours. A single-cell suspension was prepared by titrating nerves with a Pasteur pipette and passing the solution through a 20 μm Nitex filter.

##### Preparation of immunopanning plates

*CD45 plate*: One 10-cm petri dish was coated with 30 μL of goat-anti-rat IgG (H+L) secondary antibody (Jackson ImmunoResearch, 112-005-167) diluted in 10 mL of Tris-HCl (ThermoFisher Scientific, 15506017, pH 9.5) overnight at 4 °C. The plate was rinsed three times with sterile PBS and coated with rat anti-mouse CD45 antibody (BD Biosciences, 550539) diluted 1:600 in 6 mL 0.2% BSA. *Thy-1.2 plate for mouse prep*: One 10-cm petri dish was coated with 30 μL of goat-anti-mouse IgG+ IgM (H+L) secondary antibody (Jackson ImmunoResearch, 115-005-044) diluted in 10 mL of Tris-HCl (ThermoFisher Scientific, 15506017, pH 9.5) overnight at 4 °C. The plate was rinsed three times with sterile PBS and coated with anti-mouse Thy 1.2 antibody (BIO-RAD, MCA02R) diluted 1:600 in 6 mL 0.2% BSA. *Thy-1.1 plate for rat prep*: One 10-cm petri dish was coated with 30 μL of goat-anti-mouse IgM µ chain specific secondary antibody (Jackson ImmunoResearch, 115-005-020) diluted in 10 mL of Tris-HCl (ThermoFisher Scientific, 15506017, pH 9.5) overnight at 4 °C. The plate was rinsed three times with sterile PBS and coated with 6 mL of undiluted T11D7 hybridoma. *O4 plate*: One-10 cm petri dish was coated with 30 μL of goat-anti-mouse IgM µ chain specific secondary antibody (Jackson ImmunoResearch, 115-005-020) diluted in 10 mL of Tris-HCl (ThermoFisher Scientific, 15506017, pH 9.5) overnight at 4 °C. The plate was rinsed three times with sterile PBS and coated with O4 hybridoma diluted 1:6 in 6 mL 0.2% BSA.

##### Immunopanning

Briefly, dissociated cells were passed serially over a CD45 plate to remove macrophages, a Thy1.1 (for rat cells) or Thy1.2 (for mouse cells) plate to remove fibroblasts, and an O4 plate to recover Schwann cells. Schwann cells were obtained from the final plate by trypsinization.

##### Plating and Culturing

Isolated Schwann cells were plated in 10-cm dishes in Schwann cell base media supplemented with 10 ng/mL Fibroblast Growth Factor-basic (Peprotech, 450-33), 4.2 µg/mL forskolin (Sigma-Aldrich, F6886), and 10 ng/mL neuregulin 1-β1 EGF domain (R&D, 396-HB-050) growth factors and allowed to proliferate and reach confluency. Schwann cell-conditioned medium (SCCM) was collected from confluent plates every three days for 7–10 days, and cultures were transferred into new tissue culture plates. In some experiments, SCCM was collected from passaged cultures or was frozen at –80 °C until use. Where indicated, SCCM was passed through a 5 kDa molecular weight cutoff spin concentrator, and the flow through enriched for small molecules (e.g. PGE_2_) were collected.

##### Purification and culture of mouse Schwann cells

All steps were the same as from rat, above, with the following modifications. For isolation of Schwann cells from mouse sciatic nerves, reagents were downscaled to match the available number of mouse pups, and smaller plates were used accordingly for panning steps and plating. To promote proliferation and survival, mouse Schwann cells were treated with fetal calf serum (FCS, Gibco, 10437-028, 1:10) and conditioned media from confluent wild-type mouse Schwann cell cultures (1:4) until they reached confluency. After confluency, FCS and wild-type SCCM was removed from the cultures, and plates were gently washed with PBS and supplemented only with minimal Schwann cell growth medium ^3^ with no FCS; 3–4 days later, conditioned media from genotypes indicated in Figure 3 was collected.

#### Generation of rat Ptges3 knockout Schwann cell colonies

We developed and optimized the following CRISPR-based protocol for generation of clonal *Ptges3* knockout rat Schwann cell lines. Following purification from P2 rat sciatic nerves, Schwann cells were allowed to reach confluency and trypsinized from the plates for transfection experiments. Ptges3 rat Schwann cell knockout colonies were generated by transfection of synthetic sgRNAs (CRISPRevolution sgRNA EZ Kit) and Cas9 (SpCas9 2NLS Nuclease, Synthego, Menlo Park, CA) ribonucleoprotein (RNP) complexes. Three guide RNAs were designed using the Synthego CRISPR design tool to cut at exon 2 to eliminate the catalytic residue of Ptges3 (tyrosine 9): GGCAGCCUGCUUCUGCAAAG, UGCAAAGUGGUACGACCGAA, and CAAUGAAUACAUAGUCCCUU. Individual sgRNAs were allowed to form RNP complexes per manufacturer’s recommendations at 9:1 sgRNA/ Cas9 ratio and were mixed to generate a cocktail. 4 µL of RNP cocktail (including total 108 pmol sgRNA) was transfected into 250,000 Schwann cells in 20 µL primary cell transfection solution using EC134 program on an Amaxa 4D. Transfected cells were plated in 10-cm dishes, allowed to proliferate for a week, and transfected again to increase the percentage of cells with INDELs in the *Ptges3* locus. At the end of the third transfection, INDEL efficiency of the Schwann cell pool was predicted as 58% using the TIDE estimation tool ^4^. Approximately 30 Schwann cells were plated in a 10-cm tissue culture dish and were allowed to proliferate into genetically identical colonies. The proliferation of single cells was stimulated by adding conditioned media from confluent wild-type rat Schwann cell cultures (1:4 ratio to fresh Schwann cell growth medium); we found this step to be critically important for survival and proliferation of single Schwann cell clones. Schwann cell colonies were isolated using cloning cylinders and were propagated into cultures, using increasing sizes of tissue culture plates as the cells proliferated (i.e. 24-well, then 6-well). Rat *Ptges3* loci were amplified with the primers ATGATAGGTCTTAGTCCCTGTTAGC (forward) and CATAAGCCCAATAATCCATTTCT (reverse). INDELs in Schwann cell colonies were estimated with TIDE ^4^ and ICE tools (2019. v2.0. Synthego).

#### Single-cell electrophysiology

Data were measured using the patch-clamp technique in the whole-cell configuration with an Axon Axopatch 200B amplifier (Molecular Devices, San Jose, CA). Pulse stimulation and data acquisition used Molecular Devices Digidata 1322A or 1550B controlled with pCLAMP software version 10.4 or 11.1, respectively (Molecular Devices, San Jose, CA). Borosilicate glass micropipettes (Sutter Instruments, Novato, CA) were fire-polished to a tip diameter yielding a resistance of 2.0–10.0 MΩ in working solutions. All measurements were done at room temperature (20–25 °C). Data were collected from DRG neurons at least 5 minutes after establishing a seal to allow for proper equilibration of the neurons. Data were analyzed using Clampfit (Molecular Devices, San Jose, CA) and Prism 9.1 (GraphPad Software, LLC).

##### Current-clamp recordings

Data were collected on DIV7 immunopanned DRG neurons firing action potentials with overshoots > 10 mV and resting membrane potentials between –40 mV and –60 mV with < 10% variation throughout the course of the experiment. The micropipette was filled with an internal solution composed of 140 mM KCl, 0.5 mM EGTA, 5 mM HEPES, and 3 mM Mg-ATP. The pH of the internal solution was adjusted to 7.3 with KOH and the osmolarity was adjusted to 300 mOsm/L with glucose ^5^. The internal solution was aliquoted and stored at – 20 °C until needed. The external solution was composed of 140 mM NaCl, 3 mM KCl, 2 mM MgCl_2_, 2 mM CaCl_2_, and 10 mM HEPES. The pH of the external solution was adjusted to 7.3 with NaOH and the osmolarity was adjusted to 315 mOsm/L with glucose ^5^. The external solution was stored at 4 °C until needed.

DRG neurons were clamped at –80 mV. For above threshold firing patterns, action potentials were elicited by 4 x 500 ms injections of 150 pA with 5 s rest periods in between pulses. The number of action potentials fired was averaged over the four current injections to yield a single number per DRG neuron. For the firing threshold, DRG neurons were injected with 0–150 pA of current, in 5 pA intervals, until the first action potential was elicited. The firing threshold was determined to be the minimum amount of current necessary to elicit a DRG neuron to fire one action potential. Resting membrane potential (RMP) was recorded directly from the Axon Axopatch 200B amplifier.

#### DRG neuron treatments

DIV6 immunopanned DRG neurons were incubated with different treatment conditions at least 16 hours before data were recorded in electrophysiology experiments. DIV13 immunopanned DRG neurons were treated with the following conditions for 16–24 hours and fixed on DIV14 for RNAscope. The treatment conditions are as follows: **Untreated** were incubated in DRG media (DRG base medium supplemented with NGF, BDNF, and NT-3, but without forskolin; see “Plating and culturing” above); **SCCM** were incubated in half rat SCCM, half DRG media; **PGE_2_** were incubated in DRG media with 1 µM PGE_2_ (Cayman Chemicals, 14010); **Schwann cell growth media** were incubated with half Schwann cell growth media (containing bFgf and Neuregulin) and half DRG media; **Act-D** were incubated with 3 µM actinomycin-D (Sigma-Aldrich, A9415) containing half rat SCCM, half DRG media; **α-PGE2+SCCM** were incubated in half rat SCCM that had been pre-incubated with 1:5000 anti-PGE_2_ antibody (2 mg/mL, Cayman Chemicals, 10009814) and half DRG media; **EP1–EP4 antagonists** were incubated in half DRG media with concentrations as follows: 100 nM ONO-8711 (EP1 inhibitor, Cayman Chemicals, 14070), 1 µM PF-04418948 (EP2 inhibitor, Cayman Chemicals, 15016), 20 nM L-798106 (EP3 inhibitor, Cayman Chemicals, 11129), 14 µM AH-23848 (EP4 inhibitor, Cayman Chemicals, 19023). Half rat SCCM with the same drug concentrations were added after 2.5 hours of preincubation. The concentrations of the EP1– EP4 inhibitors were determined according to their dissociation constants and available empirical data ^6–8^. **DMSO** were incubated with 0.24% DMSO in DRG media; **PGD_2_** were incubated in DRG media with 1 µM PGD_2_ (Cayman Chemicals, 12010); **WT SCCM** were incubated in half wild-type rat Schwann cell wild type colony conditioned media, half DRG media; **KO_1_ SCCM** were incubated in half *Ptges3* knockout rat Schwann cell colony 1 conditioned media, half DRG media; **KO_2_ SCCM** were incubated in half *Ptges3* knockout rat Schwann cell colony 2 conditioned media, half DRG media; **WT** were incubated in half mouse SCCM, half DRG media; **CKO SCCM** were incubated in half *Ptges3^fl/fl^;Dhh^CRE/+^*mouse SCCM, half DRG media; **CKO SCCM+PGE2** were incubated in half *Ptges3^fl/fl^;Dhh^CRE/+^*mouse SCCM, half DRG media with 1 µM PGE_2_; **cHet SCCM** were incubated in half *Ptges3^fl/+^;Dhh^CRE/+^*mouse SCCM, half DRG media; **Dhh^CRE^** were incubated in half Dhh^CRE^ mouse SCCM, half DRG media; **Flox** were incubated in half Ptges3^fl/fl^ mouse SCCM, half DRG media.

#### dmPGE_2_ injection into mouse sciatic nerve

Neonatal mice (at postnatal day 0) were anesthetized by immersing in crushed ice and water (2–3 °C) in a latex glove/sleeve. 2 µL of injection solution (∼7.61 ng/µL [20 µM] dmPGE_2_ (Cayman, 14750) or 0.2% ethanol vehicle control in half PBS and half Trypan Blue [0.4%, ThermoFisher Scientific, 15250061]) solution) was injected slowly using a 10 µL Hamilton syringe. Injections were done into the sciatic nerve (into the upper thighs, between gluteus superficialis and biceps femoris muscles). Following sciatic nerve injections, upper sciatic nerve fragments were found to be completely bathed in Trypan blue. After the injection, mice were placed onto a warming blanket for recovery (approximately 2–3 minutes). Once pups were ambulatory, they were returned to the dam in their home cage. Mice were euthanized 4 hours following injection. Spinal columns were dissected and frozen on dry ice for RNAscope analysis.

#### Preparation of tissues for RNAscope

Mice were euthanized by decapitation with (P28) or without (P0) isofluorane anaesthesia to allow for rapid collection of tissues for RNAscope. The lumbar portion of the spinal column (spinal column region between the ribs and hip bones) were cut and frozen on dry ice. Sciatic nerves were dissected and placed in O.C.T. compound (Thermo Fisher Scientific, 23-730-571) to properly position and freeze on dry ice. Tissues were kept at –80 °C until sectioning.

For tissue sectioning, spinal columns or sciatic nerve were mounted in O.C.T. compound (Thermo Fisher Scientific, 23-730-571) until sectioning. 10–12 μm thick sagittal sections were collected with a Lecia CM3050 S cryostat preset to –20 °C. Sections were immediately transferred to cold Superfrost Plus (VWR, 48311-703) microscopy slide and stored at –80 °C until staining assays were performed.

#### RNA fluorescence in-situ hybridization (RNAscope)

RNAscope multiplex fluorescence (ACDbio) assays were completed according to the manufacturer’s sample preparation manuals. Multiplex fluorescence kits (ACDbio, 320851) and the HybEZ^TM^ II Hybridization system (ACDbio, 321710) were used in all RNAscope assays. Fresh frozen tissue slices were fixed for 15 minutes in 4% paraformaldehyde solution (EMS, 15710) and serially dehydrated in ethanol washes immediately before assaying. Cultured cell samples were fixed for 15 minutes in 4% paraformaldehyde solution, washed with PBS, serially dehydrated with ethanol washes, and stored in 100% ethanol at –20 °C until assaying.

After dehydration steps, both sample types were dried at room temperature for 20 minutes and then treated with pretreat IV (ACDbio, 320842) for 30 minutes. RNAscope probes were hybridized to target mRNAs, and the samples were treated with hybridization reagents Amp-FL 1-4 and counterstained with DAPI. Tissue samples were covered with microscopy cover glass (Fisherbrand, 12-544-E) and mounted with Prolong Gold antifade mount (Invitrogen, 10144). Cultured cell samples were mounted on Superfrost microscopy slides (Fisherbrand, 12-550-143) with Prolong Gold antifade mount.

##### RNAscope Probe List

| **Probe ID** | **Species** | **ACDbio Cat#** |
| --- | --- | --- |
| positive control (Polr2a, PPIB, UBC) | rat | 320891 |
| positive control (Polr2a, PPIB, UBC) | mouse | 320881 |
| negative control (Dapb) | *Bacillus subtilis* | 320871 |
| Na<sub>v</sub>1.3 | mouse | 502641 |
| Na<sub>v</sub>1.6 | rat | 411321 |
| Na<sub>v</sub>1.6 | mouse | 434191 |
| Na<sub>v</sub>1.7 | rat | 511741 |
| Na<sub>v</sub>1.7 | mouse | 313341 |
| Na<sub>v</sub>1.8 | rat | 403971 |
| Na<sub>v</sub>1.8 | mouse | 426011 |
| Rbfox3 | mouse | 453771 |
| Ptges3 | mouse | 536671 |
| Na<sub>v</sub>1.1 | mouse | 556181 |
| Na<sub>v</sub>1.2 | mouse | 423641 |
| Na<sub>v</sub>1.9 | mouse | 403531 |
| Calca | mouse | 578771 |
| Pvalb | mouse | 421931 |
| II1b | mouse | 316891 |
| II6 | mouse | 315891 |

#### RNAscope imaging and analysis

Slides and cultured cells were imaged with a Zeiss LSM 800 laser scanning confocal microscope using Zeiss Zen Blue software. A 63X oil objective was used to obtain Z-stack images of cells and tissues. Specimens were blinded in the experiments following the initial pilot experiment when possible. Samples were unblinded after imaging. Cells were chosen randomly for imaging based on DAPI or Neurofilament staining without viewing RNAscope channels. Quantification of the culture or in vivo RNAscope experiments was performed automatically using FishQuant ^9^. Except for cases in which a significant background signal was noted in a coverslip or slide, the same threshold analysis settings were applied to all specimens within an experiment. Tissues without positive control *NeuN* RNAscope signal were considered degraded and were discarded from RNAscope analysis—this was a pre-determined exclusion criterion. Cell boundaries were determined using DAPI staining and mRNA localization.

#### Immunostaining

Antibodies used in this study include the following: Neurofilament (Sigma-Aldrich, N4142, 1:1000), goat anti-rabbit IgG polyclonal antibody (CF^tm^ 405M, 20181, 1:1000), Ptges3 (knockout validated, Origene technologies, TA803433, 1:1000), Mbp (knockout validated ^10^, Abcam, ab7349, 1:100), Nav1.8 (knockout validated, Neuromab, SKU 75-166, 1:1000 for cultured cells, 1:200 for tissues [Figure 4G]), Nav1.7 (knockout validated ^11^, Neuromab, SKU 75-103, 1:200 for tissues [Figure 4G]), Nav1.1 (knockout validated ^12^, Alomone Labs, Asc-001, 1:1000), Nav1.2 (knockout validated ^13^, Alomone Labs, Asc-002, 1:1000), Nav1.6 (knockout validated ^14^, Alomone Labs, Asc-009, 1:1000), Nav1.7 (knockout validated ^15^, Alomone Labs, Asc-008, 1:1000 for cultured cells, 1:200 for tissues [Figure 4C]), Nav1.8 (knockout validated ^16^, Alomone Labs, ASC-016, 1:200 for tissues [Figure 4C]), Nav 1.5, knockout validated ^17^, Alomone Labs, ASC-005, 1:1000), Nav 1.9, Alomone Labs, AGP-030, 1:1000), and highly cross-absorbed Alexa Fluor 488-, 594-, or 647-labeled secondary antibodies (Thermo Fisher).

Antibody staining was performed on cells and tissues as previously described ^10, 18^. Briefly, cells (Schwann cells or DRG neurons) on coverslips were incubated with 3% BSA (Sigma-Aldrich, A2153) and 0.1% Triton (Sigma, T8787) containing PBS for 20 minutes. Primary antibody incubation was performed at 4 °C overnight. Coverslips were rinsed in PBS 3 times. Coverslips were then incubated with the secondary antibodies for 1–2 hours at room temperature. Coverslips were rinsed again with PBS three times, counterstained with DAPI and mounted with Prolong Gold antifade agent (Invitrogen, 10144) on Superfrost microscopy slides (Fisherbrand, 12-550-143). Alternatively, CellMask Blue stain (Fisher Scientific #H32720) was used to stain DRG neuron cell bodies. Briefly, a stock solution of CellMask Blue stain (2 mg/mL in DMSO) was diluted 1:2000 in PBS and applied to DRG neurons on coverslips for 15 minutes. Coverslips were rinsed again with PBS three times before mounting as described above. Tissues on slides were incubated with 10% goat serum (Thermo Scientific, PCN5000) in 0.1% Triton (Sigma, T8787) containing PBS for 1 hour following RNAscope. Mbp primary antibody (knockout validated ^10^, Abcam, ab7349, 1:100) incubation was done in 1% goat serum in PBS overnight; slides were rinsed for 30 minutes with 0.1% Triton containing PBS; this process was repeated four times. Secondary antibody solution (Alexa Fluor 488, 1:500) was performed at room temperature for 1-2 hours. Slides were rinsed with 0.1% Triton containing PBS for 30 minutes; this process was repeated four times. Slides were counterstained with DAPI and mounted with Prolong Gold antifade mount (Invitrogen, 10144).

Following adaptations were implemented to increase the signal/noise ratio from Nav antibodies:

##### For DRG neurons on coverslips

Following fixation, coverslips were incubated with 10% goat (Thermo Scientific, PCN5000) or donkey serum (Abcam, Ab7475) and 1% Triton (Sigma, T8787) containing PBS for 1 hour before primary antibody incubation.

##### For DRG tissues

For immunostaining of neonatal DRG tissues, a light fixation of 4%PFA in PBS solution was performed on fresh-frozen tissue sections at room temperature for 15 minutes. Following fixation an antigen retrieval step was performed by incubating slides in 0.2% hydrogen peroxide, 0.3% Triton and 20% methanol solution containing PBS for 25 minutes at room temperature. The blocking step was done with 0.3% Triton, 5% donkey serum (Abcam, Ab7475), 5% goat serum (Thermo Scientific, PCN5000) and 50 mM glycine containing PBS for one hour at room temperature. Primary antibody incubations were done in 0.1% Triton and 5% donkey serum containing PBS (Abcam, Ab7475).

For immunostaining of postnatal DRG tissues, a light fixation of 4% PFA in PBS solution was performed on fresh-frozen tissue sections at room temperature for 15 minutes. A more stringent permeabilization step was applied with 2.5% Triton and 10% goat serum (Thermo Scientific, PCN5000) containing PBS at room temperature for 1 hour. Primary antibody incubations were done in 0.1% Triton and 5% goat serum containing PBS.

#### TUNEL assay

TUNEL assay was performed with In Situ Cell Death Detection Kit, TMR Red (Millipore Sigma, # 12156792910, Lot # 58382200) based on manufacturer’s protocol. Briefly, fresh-frozen tissue sections were fixed with 4% PFA in PBS solution at room temperature for 20 minutes. Tissue sections were then washed with PBS for 1 hour and permeabilized with protease IV (ACDbio, 320842) for 30 minutes. 50 μl of TUNEL reaction mixture (5 μl enzyme solution + 45 Label Solution) was applied to slides for 1 hour at 37 °C. Coverslips were rinsed again with PBS three times, counterstained with DAPI and mounted using VECTASHIELD antifade mounting medium with DAPI (Vector Laboratories, #H-1200).

#### Western immunoblotting

Schwann cells were lysed in RIPA buffer (25 mM Tris–HCl, 150 mM NaCl, 1% Triton X-100, 1% sodium deoxycholate, 0.1% SDS) supplemented with protease inhibitor cocktail tablets (Roche, 04693159001). Clarified lysates were mixed with NuPAGE LDS Sample Buffer (Invitrogen, NP0007) and NuPAGE Sample Reducing Agent (Invitrogen, NP0009) and boiled for 10 min. Lysed samples were subjected to electrophoresis using a XCell SureLock Mini-Cell Electrophoresis System (Invitrogen, EI0002) and a Mini-PROTEAN system (Bio-Rad, 1658026FC). After separation, protein was transferred to a nitrocellulose membrane (Life Technologies, LC2001), followed by standard blotting procedure. Primary antibodies were obtained from Abcam and OriGene: Anti-alpha Tubulin (Abcam, ab52866, 1:1000) and Ptges3 (knockout validated, Origene technologies, TA803433, 1:1000). HRP-conjugated secondary antibodies (CST, 7074P2 and 7076P2, 1:10000) were used for protein band detection. Protein bands were visualized by chemiluminescence (Thermo Scientific, 34075) using a ChemiDoc imaging system.

#### (DESI)#Mass spectrometry of SCCM

Briefly, CH2Cl2 was infused through fused capillary tubing (100 mm i.d., 360 mm o.d.) of a custom DESI source ^19^ a rate of 50 mL/min (0 or 5 kV spray voltage). An N2 sheath gas (0.6 L/min) generated a microdroplet spray directed toward a paper surface onto which a solution containing SCCM extract (flow-through from 5 kDa molecular weight cutoff filter) in CH2Cl2 was deposited. The secondary microdroplets were sampled by an LTQ Velos Orbitrap mass spectrometer (Thermo Fisher Scientific) in the negative ion mode. The PEEK tubing connected to the outlet of the mixing tee was connected to an ESI source. DRG neuron media was used as a control medium. Peaks that appeared in the chromatogram of the purified 5 kDa FT SCCM, but were missing from the DRG neuron media, were identified as of interest. The tallest peaks at 351.2182, 353.1976, 353.3398, and 354.9219 were analyzed against known metabolites using the metabolomics database METLIN with ± 75 ppm mass accuracy. The majority metabolites with mass 351.2182 ± 75 ppm are products of the arachidonic acid pathway. A similar DESI-MS experiment with a sample commercial PGE_2_ in CH2Cl2 provided a mass spectrum that closely matches the data generated from the 5 kDa FT media sample.

#### RNAseq of Schwann cells from the rat sciatic nerve

RNAseq of acutely isolated rat Schwann cells is described fully elsewhere (A.B.L. & S.A.S., submitted). Briefly, immunopanning from uninjured P18 rat sciatic nerve was used to acutely purify Schwann cells as previously described ^3, 20^. A minimum of 10 sciatic nerves was used for each sample of purified cells. RNA from purified Schwann cells was isolated at the end of the immunopanning procedure by scraping the O4 positive panning dish according to the instructions of the RNeasy MICRO kit. RNA quality was verified through Bionalyzer analysis. All samples submitted for RNASeq had a RNA integrity number of 8 or above.

For RNAseq library construction and sequencing, total RNA that passed QC standards proceeded to library synthesis using the Ovation^®^ RNA-seq system V2 (Nugen 7102). First- and second-strand cDNA synthesis and SPIA amplification were performed following the manufacturer’s instructions, and cDNA was fragmented with a sonicator (Covaris S2) using the following parameters: duty cycle 10%, intensity 5, cycles/burst 100, time 5 minutes. A Next Ultra RNA-seq library prep kit for Illumina (NEB E7530) and NEBNext^®^ multiplex oligos for Illumina® (NEB E7335 E7500) was then used to perform end repair, adaptor ligation, and 5–6 cycles of PCR enrichment according to the manufacturer’s instructions. The quality of the libraries was then assessed by bioanalyzer and qPCR, and high quality libraries were sequenced by an Illumina NextSeq sequencer to obtain 150 bp pair-end reads.

For bioinformatic processing of RNAseq data, FASTQ files were first groomed using a FASTQ groomer and then mapped using TopHat2, which invokes Bowtie as an internal read mapper. The paired-end option was selected, and rat genome Rnor_6.0 was used as the reference genome. Cufflinks was run to assemble transcripts and estimate expression level as fragments per kilobase of transcript sequence per million mapped fragments (FPKM). All subsequent data analysis was performed using R software (https://www.r-project.org/). All raw fastq files from the RNA-seq data were deposited in the National Center for Biotechnology Information (NCBI) Gene Expression Omnibus (GEO, accession number GSE177037).

#### Conditional knockout Ptges3 mouse line generation

To generate cell-type-specific knockout of the *Ptges3* gene, a *Ptges3^fl/fl^* mouse was generated using CRISPR-mediated genome recombination (Stanford Transgenic, Knockout and Tumor model Center, Stanford, CA). Two loxP sites were introduced at intron 1 and intron 3 to flox exons 2 and 3. After Cre recombination, *Ptges3* exon 2 and exon 3 will be excised, resulting in a 62 amino acid deletion, shift of the downstream reading frame, and knockout of Ptges3 (note that this deletion strategy causes loss of the critical catalytic residue for PGE_2_ production, and has been successfully utilized in a global knockout to ablate Ptges3 function ^21–23^. Four guide RNAs (gRNA) were initially designed, and single-strand guide RNAs (sgRNA) were purchased from Synthego (Menlo Park, CA). After in vitro validation by DNA cutting efficiency, two gRNAs were chosen for generating *Ptges3* floxed mice. Guide RNA1 (GGCGTGCTGCTGTTATCTTA) cut at intron 1, 210 bp upstream of exon 2. Guide RNA4 (CCCAGAATAAACTCCCTATT) cut at intron 3, 67 bp downstream of exon 3. Donor DNAs were designed for the insertion of the loxP sites. Donor DNAs contained a 1 kb homologous arm upstream of intron 1 loxP, exon 2, exon 3, and a 1 kb homologous arm downstream of intron 3 loxP (Figure S10E). The two loxP sites inserted at the gRNA cutting sites also destroyed gRNA recognition sequences to prevent the gRNA-Cas9 complexes from cutting recombinants. Donor DNAs were cloned and amplified in a vector plasmid, a 2.7 kb double-strand linear DNA fragment that was cut off from vector DNA and purified for mouse zygote pronuclear injection.

To generate Ptges3^fl/fl^ mice, a mixture of gRNA (10 ng/µL), Cas9 protein (30 ng/µL) (IDT, Iowa) and donor DNA (3 ng/µL) were injected into C57Bl/6J mouse zygote pronuclei. A total of 150 zygotes were injected. Of 15 pups born, three pups were identified by genotyping to have loxP sites at the designed location.

The genotyping strategy to confirm intron 1 loxP integration by PCR included using a forward primer at the upstream of the homologous arm (GGCCACCTCAAGTAGTAGAGCTTGGCTACATAC) and one primer at intron 3 loxP (CTCACAGCAACAAACCCATAACTTCGTATAGC). The same strategy for confirming the intron 3 loxP was used: one PCR primer at the downstream of the homologous arm (AGCAAAGGGGCAAGAGGCAAGCG) and one primer at the intron 1 loxP (CCTCATCTCCCAGTTTAAACTCCCTATAACTTCG). The amplified fragments were sequenced to confirm the integration of loxP sequences and exons.

*Ptges3^fl/fl^* founder mice were bred with C57Bl/6J for heterozygotes; backcross breeding can eliminate possible mosaics during CRISPR mediated recombination. Heterozygote mice were screened using primers—forward (GCTGTACTGGAACTCACTCTGTAGATG) and reverse (GCAAATTACCTGAAAGTAAGTTTGGATTTTTCA)—and sequenced to confirm the presence of a loxP site using ICE tool (2019. v2.0. Synthego). Heterozygous (backcrossed) mice were intercrossed to generate homozygous *Ptges3^fl/fl^* mice. Homozygous *Ptges3^fl/fl^* mice were also screened for correct integration of loxP sites and lack of silent background mutations that could be caused by CRISPR/Cas9 genome editing. The genotyping strategy to confirm intron 1 loxP by PCR included a forward primer at the upstream of the homologous arm (GGCCACCTCAAGTAGTAGAGCTTGGCTACATAC) and one primer at the downstream of intron 3 loxP (GCAAATTACCTGAAAGTAAGTTTGGATTTTTCA). The amplified fragment was sequenced by PCR to confirm the integration of loxP sequences and the integrity of exon 2. A similar strategy for confirming the intron 3 loxP was used: one primer at the upstream of exon 3 (GCTCCTGTCTCTAGGATTGTGTAGAATTG) and one PCR primer targeting the homologous arm (GTAGTTTAAATGAAAGTGGCCTCACAGG) was used. The PCR product was sequenced to confirm the integration of intron 3 loxP sequence and the integrity of exon3.

*Ptges3^fl/+^* mice were bred to *Dhh^CRE/+^* mice to generate *Ptges3^fl/+^;Dhh^CRE/+^* mice. *Ptges3^fl/+^;Dhh^CRE/+^* mice were bred to *Ptges3^fl/fl^* mice to generate Schwann cell specific knockout of Ptges3. Recombination in Schwann cells of heterozygous and full conditional knockout animals (*Ptges3^fl/+^;Dhh^CRE/+^* or *Ptges3^fl/fl^;Dhh^CRE/+^)* was confirmed using primers that amplify a 336 bp-long recombined DNA (1226 bp long before recombination, not amplified with the settings used): forward (CCTCATCTCCCAGTTTAAACTCCCTATAACTTCG) and reverse (GTAGTTTAAATGAAAGTGGCCTCACAGG). In addition, control primers detecting a 168 bp region of the *Mbp* locus were used to test the integrity of the DNA samples: forward (AGCTCTGGTCTTTCTTGCAG) and reverse (CCCCGTGGTAGGAATATTACATAAC). After the verification of the line with the studies outlined above, mice were genotyped by Transnetyx.

#### Transmission electron microscopy sample preparation

Transmission electron microscopy (TEM) was completed in the Stanford Cell Sciences Imaging Facility. Samples were prepared according to previously published protocols ^24^. Samples were initially washed in chilled Karlsson-Schultz fixative (2.5% glutaraldehyde, 4% PFA in phosphate buffer, pH 7.3) and incubated in 2% OsO_4_ for four hours at 4 °C. Samples were then serially dehydrated at 4 °C and embedded in EmBed812 (EMS, 14120). 80 nm sections were taken using an UC7 (Leica, Wetzlar, Germany) and were collected on formvar/Carbon coated 100 mesh Cu grids. Sections were then stained for 40 seconds in 3.5% uranyl acetate in 50% acetone followed by staining in Sato’s lead citrate for 2 minutes. TEM images were obtained using a JEOL JEM-1400 120kV with a Gatan OneView 4k X 4k digital camera. Quantification of TEM images was performed manually using Fiji/Image-J.

#### Mouse behavior assays

Mice were older than 2 months at the time of behavioral testing. Male and female mice were assessed separately; the results were displayed for both sexes, and the sex of the mice is indicated in the graphs and figure legends. One cohort of mice was used in activity chamber, hot plate, Hargreaves, Von-Frey, horizontal ladder, rotarod and CatWalk tests. An additional independent cohort was used for Hargreaves test to increase power (based on power analysis using pilot data; power=0.8, alpha=0.05).

##### Activity chamber assay

The assessment took place in an Open Field Activity Arena (Med Associates Inc., St. Albans, VT. Model ENV-515) mounted with three planes of infrared detectors within a specially designed sound-attenuating chamber (Med Associates Inc., St. Albans, VT. MED-017M-027). The arena was 43 cm (L) × 43 cm (W) × 30 cm (H), and the sound-attenuating chamber was 74 cm (L) × 60 cm (W) × 60 cm (H). The mice were placed in the corner of the testing arena and allowed to explore the arena for 10 minutes while being tracked by an automated tracking system. Parameters, including distance moved, velocity, rearing, and times spent in the periphery and center of the arena were analyzed. Periphery was defined as the zone 5 cm away from the arena wall. The arena was cleaned with a 1% Virkon solution at the end of each trial. Experimenters were blinded to mouse genotype during the experiment.

##### Hot plate assay

The hot plate apparatus (IITC Inc. Model 39) was set to a temperature of 52 ± 0.2 °C. Mice were placed on the surface of the hot plate and covered by a transparent glass cylinder, 25 cm high and 12 cm diameter. A 30 second cutoff time is assigned in this protocol. The application of cutoff time minimizes the exposure of animals to the hot plate while protecting them from distress. A remote foot-switch pad is used to control the start/stop/reset function. The latency time is recorded when the first hind paw licking, removal, or jumping-off occurs. Experimenters were blinded to mouse genotype during the experiment.

##### Hargreaves test

Mouse hind paw heat thresholds were determined using the Hargreaves test ^25^. Mice were acclimated in the testing room over 2 days for 15 min/day on a wire rack in-cylinder plastic tubes followed by 45 minutes in Hargreaves boxes (Stoelting). On testing day, mice were initially acclimated in the testing room on wire racks for 15 minutes followed by 15 minutes in Hargreaves boxes. Mice were then subjected to 3–5 trials of radiant heat application to the hind paw using a Hargreaves apparatus set to 25% of infrared generator intensity (25 mW/cm^3^) with the duration of the stimulation set at a maximum of 30 seconds to avoid skin injury. Paw withdrawal latency was measured by an automated system that shut off when the mouse withdrew its hind paw from the heat source ^26^. Mice were tested for only 3 total trials if no outliers between the first three trials were found (outlier defined as > 6 seconds difference from nearest withdrawal time). If outlier measurements were recorded, or if there was a need for tiebreakers, an extra trial (trial #4 and/or #5) was done. At least 5 minutes were allowed to pass between each trial for any given mouse. At the end of testing, outlier measurements were omitted, and remaining trial data were averaged to generate each withdrawal time for each mouse. Experimenters were blinded to mouse genotypes during the experiment.

##### Von-Frey test

Von-Frey test is used to evaluate baseline mechanical nociception. Mice were initially acclimated on a wire rack for at least 40 minutes, followed by application of a series of 8 von-Frey filaments (Stoelting) ranging in gram force from 0.007 to 6.0 g to the plantar hind paw, starting with the 0.16 g filament. Positive responses were recorded when the applied force-induced a rapid withdrawal of the paw away from the stimulus filament within 4 seconds. The up-down statistical method ^27^ was used to calculate the 50% withdrawal mechanical threshold score for each mouse. These baseline scores were then averaged for each experimental group across. Experimenters were blinded to mouse genotype during the experiment.

##### CatWalk Gait analysis

Gait analysis was done using the video-based CatWalk XT system (v9.1, Noldus). Mice were placed on a linear track, and multiple trials were performed until each mouse had completed three compliant runs, traversing the track continuously without stopping. The software automatically assigned the positioning of the paws, but the experimenter corrected the errors manually. The experimenter was blinded to the genotypes while doing the experiment and analysis.

#### Rotarod Test

Mice were placed on a rotarod apparatus to test sensorimotor behavior. Mice were acclimated to the rotarod at a constant speed of 4 RPM for 1 min. Each animal was given three trials, each 5 minutes, with the rotarod at 32 RPM constant speed. There were at least 15 minutes in between trials. Latency to fall was recorded and three trials were averaged. Mice that fall within 5 seconds were given a second trial. If mice hold onto the rod for more than 2 consecutive turns, this was considered a fall. Experimenters were blinded to mouse genotypes during the experiment.

##### Formalin Injection Assay

50 μl of 1% formalin was injected to the right hind paw of the mice. Pain responses (licking and biting the injected paw) was recorded for 1 hour, video tracking and analysis were performed using EthoVision XT. For one Ptges3-Flox and two Ptges3-cKO males, video tracking was interrupted at 40 minutes due to hardware errors, these animals were shown in pain response timing views (Figure S21B) but excluded from the calculations of total inflammatory pain response in Figure 6G and Figure S21C.

##### Horizontal Ladder Test

Mice were placed on a horizontal ladder (Maze Engineers, Item # 5103, 60 cm) with irregularly placed rungs (1-4 cm) and videotaped from the side. Forelimb and hindlimb slips, and the use of corridor walls (chimneying) were quantified. The experimenter was blinded to the genotypes while doing the experiment and analysis.

#### Calcium imaging of DRG Neurons

Lumbar DRG ganglia (L2-L6) were dissected from Ptges3-Flox or Ptges3-cKO mice at P18/P19 postnatal stages and placed in neuron medium (DMEM and F12 [Gibco^TM^, 10565018], 10% fetal calf serum [Gibco^TM^, 10437028], 100 U/mL +penicillin and 0.1 mg/mlL streptomycin [Gibco^TM^, 15140122]) with 30 μM ActinomyosinD to inhibit new transcription due to dissections). Dissected DRG ganglia were placed in a collagenase/dispase cocktail composed of 1 mg/mL collagenase (Roche, 10103578001) and 2.9 mg/mL dispase (Roche, 101103578001) in 1X HEPES buffered saline solution (Sigma-Aldrich, 51558) containing 40 U of papain (Worthington LS003126), 100 μl DNAse (0.02 mg/mL) and 15 μM ActinomyosinD and placed in 37 °C for 25 minutes. DRG ganglia were dissociated in Lo-ovomucoid inhibitor solution in (DMEM and F12 [Gibco^TM^, 10565018]) with 3 μM actinomycin-D using a glass Pasteur pipette. Dissociated cells were plated in MatTek dishes (MatTek Corporation P35G-1.5-20-C) coated with 0.01 mg/mL PDL in water 30 minutes at room temperature and 1 mg/mL laminin (R&D Systems, 3400-010-02) diluted 1:200 in Neurobasal media (Thermo Fisher Scientific, 21103049) at 37 °C for 4 to 24 hours. Cells were allowed to adhere to MatTek dishes for 1 hour. Subsequently, media was replaced with FluoroBrite DRG media (Fisher Scientific A1896701) that contains all DRG media ingredients mentioned above except for DMEM and Neurobasal. Next, 5 μM cell permeable calcium indicator, Fluo-4-AM (Invitrogen, F14201) was added onto cells for 20 minutes. Media was changed with fresh FluoroBrite DRG media and cells were further incubated for 20 minutes. Cells were imaged on an Opterra II Multipoint Swept Field Confocal outfitted with a humidified, temperature-controlled microscope enclosure (Okolab microscope enclosure, H201-Temperature Unit, CO2 controller, HM-Active Vibration Free Humidity Controller with humidity sensor and temperature-controlled tube). Imaging was performed using the 20x/.75NA objective and Perfect Focus to prevent z-plane drift during imaging. Images were acquired every second in 488 channel with 70 μM slit and 100 ms exposure time with 15% laser power. Cells were imaged at baseline for approximately 160 frames and then an equal half volume of 10 μM veratridine in FluoroBrite DRG media (5 μM final concentration) was applied to cells and imaged for 240 further frames. Calcium imaging data was analyzed using Fiji to select regions of interest (cell bodies from first time point, blinded to genotype and calcium response) and MATLAB scripts to calculate max f/f0 for each cell.

#### Dissociation of DRG ganglia for single-cell RNA-seq (scRNA-seq)

DRG ganglia from all axial levels were dissected from Ptges3-Flox or Ptges3-cKO mice at P4 postnatal stage and placed in Neurobasal medium supplemented with 12.5 mM _D_-glucose and 100 U/mL +penicillin and 0.1 mg/mL streptomycin [Gibco^TM^, 15140122]). Next, dissected DRG ganglia were placed in a collagenase/dispase cocktail composed of 1 mg/mL collagenase (Roche, 10103578001), 2.9 mg/mL dispase (Roche, 101103578001), 40 U of papain (Worthington LS003126), 100 μl DNAse (0.02 mg/mL) in 1X HEPES buffered saline solution (Sigma-Aldrich, 51558) and placed in 37 °C for 20 minutes. DRG ganglia were dissociated in Lo-ovomucoid inhibitor solution in PBS. Cells were counter-stained with trypan blue counted under a hemocytometer to assess cell density and health (cell viability was measured as greater than 92%), and subsequently resuspended in Neurobasal medium supplemented with 12.5 mM _D_-glucose and 100 U/mL +penicillin and 0.1 mg/mL streptomycin [Gibco^TM^, 15140122]) as 1000 cells per μl.

#### Single-cell library preparation, sequencing and analysis

Libraries were prepared using the Single Cell 3ʹ v.3.1 according to the manufacturer’s protocol (10x Genomics), targeting 10,000 cells per sample. 12 cycles of cDNA amplification were done for all samples. Individual libraries were quality checked on an Agilent 4200 Tapestation 827 using D5000 screen tape. Next, KAPA library quantification kit (#KK4923) was used for qPCR on a BioRad CFX96 RT PCR thermal cycler. Libraries were sequenced on NextSeq 550 High Output platform. Initial gene expression tables for barcodes were generated using CellRanger version 5.0.1 (10X genomics) using default parameters. Gene expression tables were than imported to Phyton and analyzed using Scanpy ^28^.

#### Single-cell RNAseq Quality Controls and Exclusion

Cells containing fewer than 1000 genes and genes that were detected in less than 3 cells were removed from the analysis. The remaining 8248 cells from Ptges3-cKO and 7366 cells from Ptges3-Flox mice were used for further analysis. Sensory neurons were identified by expression of the sensory neuron marker *Avil* (expression level >0.5), revealing 1507 individual DRG neurons in Ptges3-cKO mice (18.2%) and 2079 individual DRG neurons in Ptges3-Flox mice (36.77%). A dictionary of marker genes for each DRG subtype was generated based on previously published studies ^29–31^ and used to identify DRG subclusters identified in Ptges3-Flox mice. DRG neurons from Ptges3-cKO mice were integrated with the embeddings and annotations of Ptges3-Flox neurons using Scanpy’s ingest function. Differential gene expression analysis were done with Seaborn and Statannotations using t-test for independent samples ^32^.

#### Bulk RNAseq library preparation, sequencing and analysis

Total RNA from DIV14 immunopanned DRG neurons (untreated [n=3], SCCM [n=4] or PGE_2_ [n=4] treated) was purified using RNeasy Plus mini kit (Cat # 74134). Genomic DNA was eliminated using eliminator spin columns provided by the manufacturer. RNA quality and quantity were assessed using Agilent 2100 Bioanalyzer system, samples with high RNA integrity numbers (>8) were used for further analysis. KAPA/Roche Stranded mRNA-Seq kit KK8420 (Cat # 07962193001) was used to prepare poly-A enriched mRNA libraries. Samples were sequenced using Illumina Hiseq 2 x 75 platform at Stanford Genomics Center. Sequenced reads were aligned to the rat reference genome (Rnor_6.0) using Geneious Prime 2021.0.1 (https://www.geneious.com). Expression levels were compared using DEseq2 parametric fit type^33^. Ingenuity Pathway Analysis (IPA) was performed through QIAGEN IPA (QIAGEN Inc., https://digitalinsights.qiagen.com/IPA). Biological replicates with evidence of cell death were excluded from the subsequent analysis.

#### Compound Action Potential Measurement

The ex vivo electrophysiology setup and methods used in this study have been detailed elsewhere^34, 35^. In summary, the ex vivo electrophysiology system was composed of a high current stimulus isolator (Isostim A385R, World Precision Instruments, Sarasota, Florida, USA), a custom designed (TinkerCAD https://www.tinkercad.com/, Waltham, Massachusetts, USA) and 3-D printed electrophysiology (FormLab Form 2 3-D printer, Somerville, Massachusetts, USA) chamber with two suction electrodes, an extracellular amplifier/recording device (SYS-DAM80, World Precision Instruments, Sarasota, Florida, USA), and software for stimulating, recording, and analyzing CAP signals (WinWCP V5.5.5 and WinEDR V3.9.1, https://spider.science.strath.ac.uk/sipbs/software_ses.htm, John Dempster, University of Strathclyde Glasgow).

Sciatic nerves were dissected immediately after euthanasia (decapitation with isofluorane anaesthesia) and transferred to 20-40 ml of “cold” (in a container of ice, temperature 0.7 °C - 2.0 °C) HEPES solution (136 millimolar (mM) NaCl, 11mM glucose, 4.7mM KCl, 2.4mM CaCl_2_, 1.1mM MgCl_2_, 1mM NaHCO_3_, 10mM HEPES, pH 7.2-7.4, oxygenated continuously with 100% O_2_). After ≥ 60 min in the cold HEPES solution, the nerve was transferred to the chamber along with 3 ml of “room temperature” (17.0 °C - 21.0 °C) HEPES solution; the proximal end of the nerve was pulled into the stimulating suction electrode, and the distal end of the nerve was pulled into the recording suction electrode. Stimulator settings for A-alpha, A-beta, and A-delta fibers were: stimulation pulse width 0.2 milliseconds (ms), recording interval of 50 ms, and repeated for 60 events. Stimulator settings for C fibers were: stimulation pulse width 0.5 ms, recording interval of 2 s and repeated for 6 events. Amplifier settings were as follows: 0.1 Hz low filter, 3 kHz high filter, 1000x gain.

Once placed in the chamber suction electrodes, the nerves were allowed to “rest” for 20 minutes before stimulation and recording occurred. The nerves were first stimulated with 0.1 milliamperes (mA) of current to check for the presence of a compound action potential (CAP). For A-fiber action potentials (A-alpha/A-beta, A-delta), the current was then successively increased (0.1-0.2 mA increments) and the CAP amplitude measured until less than a 5% change in peak amplitude was observed. For C-fiber action potentials, the amplitude was assessed at the following currents, 1, 2, 5, 10, 50, and 100 mA. Action potentials of all fiber types (A-alpha/A-beta, A-delta, C) were recorded and action potential components were analyzed (amplitude, latency to 100% peak, rate of rise, and area under the curve) using WinEDR V3.9.1(https://spider.science.strath.ac.uk/sipbs/software_ses.htm, John Dempster, University of Strathclyde Glasgow).

#### Statistical analysis

All statistical analyses were done using GraphPad Prism 9.10 (216). We pre-determined that using 3–5 biological replicates for culture or in vivo experiments were sufficient based on previously published studies ^10, 36^. In all electrophysiology and immunohistochemistry (culture) experiments, cells were compared in a two-tailed unpaired t-test when there were two groups or in a one-way ANOVA followed by Tukey’s multiple comparisons when there were three or more groups. A one-tailed ratio paired t-test was used to compare the fold change increases in response to SCCM or PGE_2_ treatments in Figures 1G–I and Figure 2D. In Figure 2E, due to the lack of some biological replicate pairs (e.g., due to cells washing off the coverslips during RNAscope assays), a mixed effect analysis with Tukey’s multiple comparisons was performed since ANOVA cannot handle missing values. In Figure S4 and Figure S9, cells were compared in one-way ANOVA followed by Tukey’s multiple comparisons. In in vivo experiments, experimental groups were compared in a one-tailed unpaired t-test when there were two groups; or in a one-way ANOVA followed with Tukey’s multiple comparisons when there were three or more groups. Mann Whitney test was used for non-parametric comparisons of two groups where distributions are unmatched. Percent frequency of axon diameter was compared in a two-way ANOVA followed with Tukey’s multiple comparisons in Figure S16D. For behavioral experiments, data were analyzed using ROUT test to identify outliers with Q = 1% power (exclusion criterion established *a priori)*. Two outliers in the control group from the Hargreaves test (Figure 6F) and two outliers in the Ptges3 cKO group from the rotarod test (Figure 6H) were identified and excluded from the data analysis; there were no other outliers/excluded mice in any other experiment.

#### Resource Availability

##### Lead contact

Further information and requests for resources and reagents should be directed to and will be fulfilled by the lead contact, J. Bradley Zuchero.

##### Materials and data availability

Requests of plasmids and mouse lines described in this study will be available upon request from the lead contact, and will be deposited at Addgene (plasmids) and Charles Rivers (mouse lines). Raw fastq files resulting from the bulk RNA-seq studies were deposited in the National Center for Biotechnology Information (NCBI) Gene Expression Omnibus (GEO, accession number GSE177037). Single-cell RNA-seq data will be deposited at NCBI GEO prior to publication. All raw data used in the manuscript will be deposited at Dryad at the time of acceptance for permanent storage and free access.

## SUPPLEMENTAL FIGURES AND FIGURE LEGENDS

**Figure S1.**
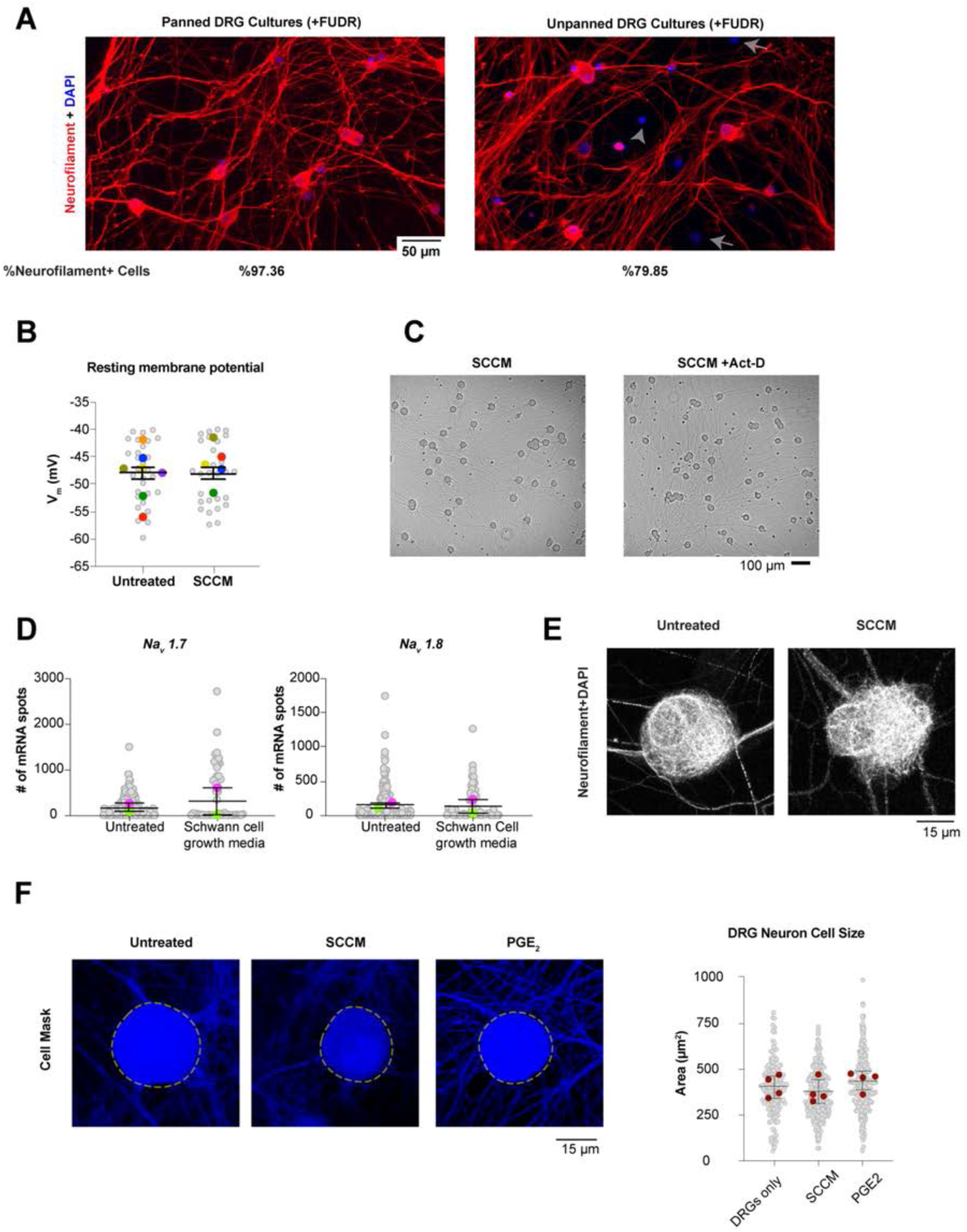
Immunopanned DRG neuron cultures are highly pure, containing more than 97% neurons. (**A**) Images show Neurofilament and DAPI staining in immunopanned DRG neurons as described in Figure 1A or unpanned DRG neurons plated after dissociation without the panning steps. Even with the use of the mitotic inhibitor FUDR, unpanned DRG neuron cultures had significant contamination with non-neural cells, negatively stained for Neurofilament. (**B**) Untreated and SCCM treated DRG neurons exhibited statistically similar resting membrane potentials. *n* = 29 total cells from 5–7 distinct biological replicates per treatment group. (**C**) Bright field images indicate that DRG neuronal health is unaffected by Act-D treatment, consistent with prior studies. (**D**) The addition of Schwann cell growth media to DRG neurons was insufficient to increase expression of *Na_v_1.7* and *Na_v_1.8* transcripts, suggesting the effects of SCCM are specific to a Schwann cell-secreted molecule(s). Gray circles, mRNA number of an individual DRG neuron for the indicated genes; colored circles, the average # of mRNAs per cell in each biological replicate. Mean ± SEM is shown for the biological replicates (not significantly different in a paired t-test). *n* = 2 distinct biological replicates per group. (**E**) Immunopanned DRG neurons showed normal cell growth and neurofilament staining, all unaffected by SCCM treatment. (**F**) DRG neuron cell size, measured with Cell Mask, was unaffected by SCCM or PGE_2_ treatments. Mean ± SEM is shown for the biological replicates (not significantly different in an one-way ANOVA and Tukey test). *n* = 4 distinct biological replicates per group.

**Figure S2.**
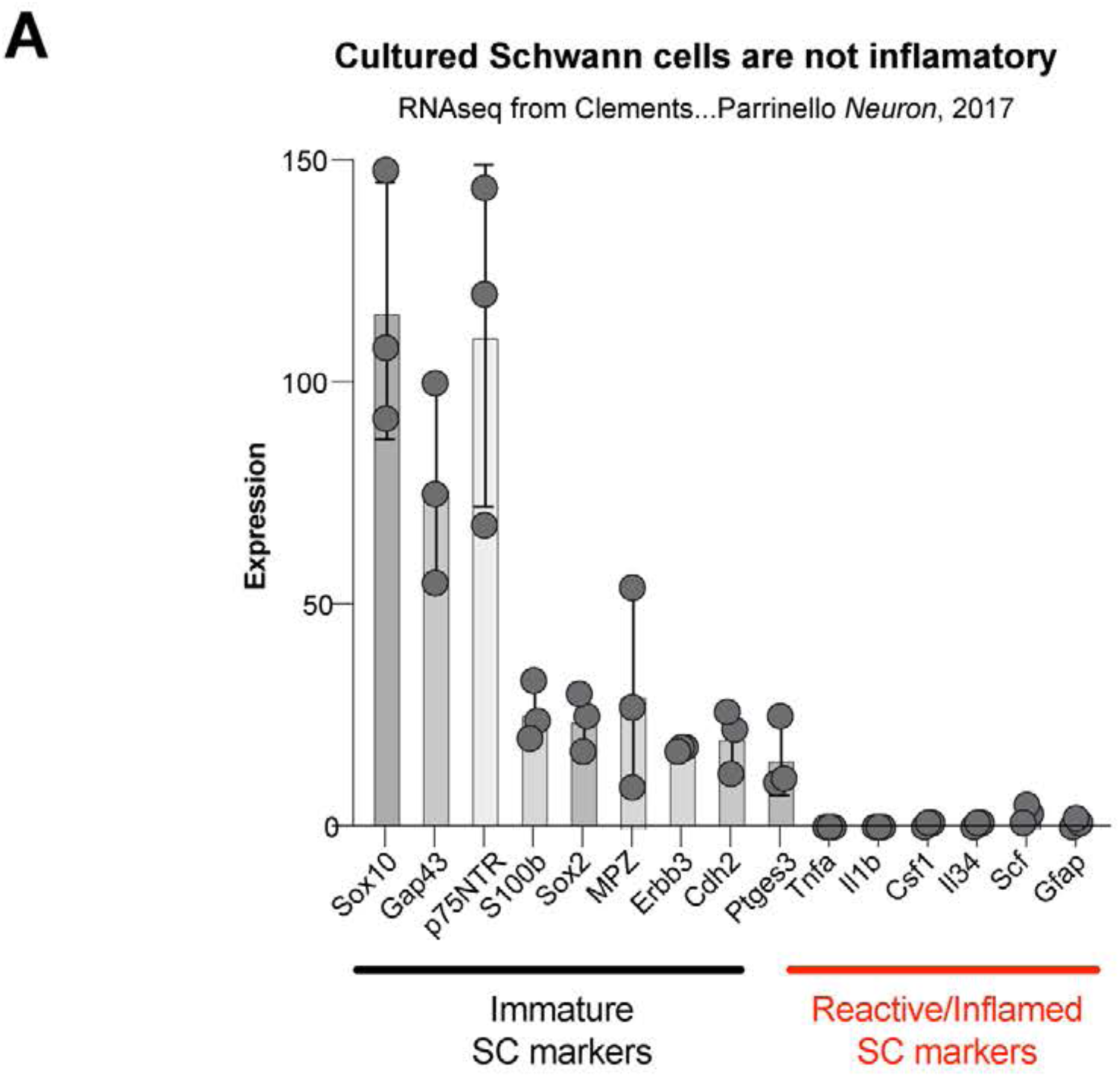
Isolated Schwann cells are not inflammatory. (**A**) Data from a published RNAseq study^37^ shows that cultured Schwann cells isolated from postnatal rats express immature Schwann cell markers but not inflammatory or reactive markers associated with injury or disease states^38, 39^.

**Figure S3.**
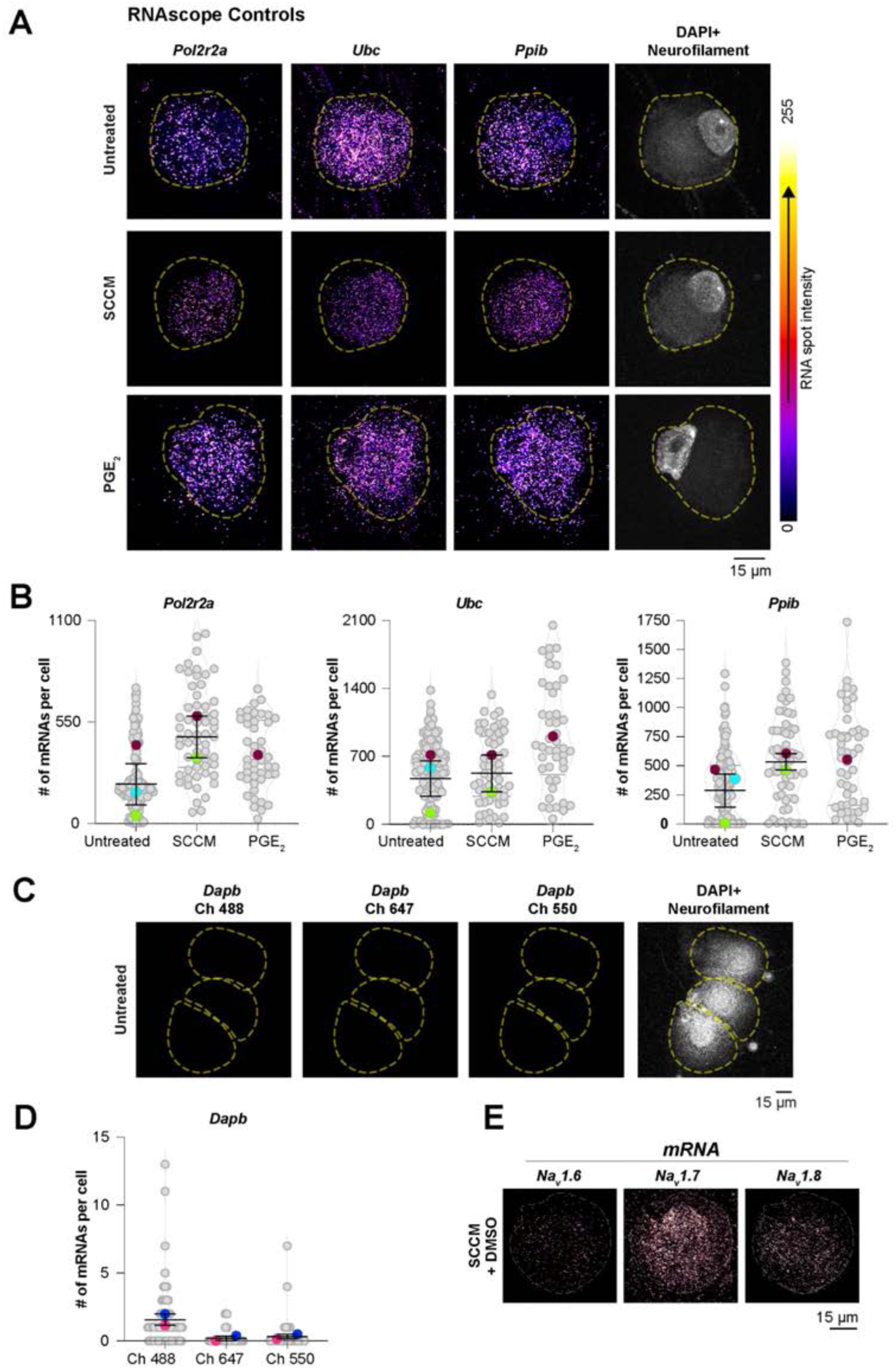
Specificity controls for RNAscope. (**A**) DRG neurons expressed control housekeeping genes (*Pol2r2a*, *Ubc*, and *Ppib*) at similar levels when cultured alone or treated with SCCM or PGE_2._ (**B**) Quantification of mRNA spots using FishQuant for indicated genes and conditions. *n* = 3, 2 and 1 biological replicates per group, respectively. (**C**) Negligible background in RNAscope experiments detected using the negative control probe designed for the bacterial gene *dapB*. (**D**) Quantification of mRNA spots using FishQuant for the *dapB* gene detected in three different channels. (B and D) Gray circles, mRNA number of an individual DRG neuron for the indicated genes; colored circles, the average # of mRNAs per cell in each biological replicate. Mean ± SEM is shown for the biological replicates. *n* = 2 distinct biological replicates. (**E**) DMSO+ SCCM increases DRG Na_v_ expression similar to SCCM treatment alone.

**Figure S4.**
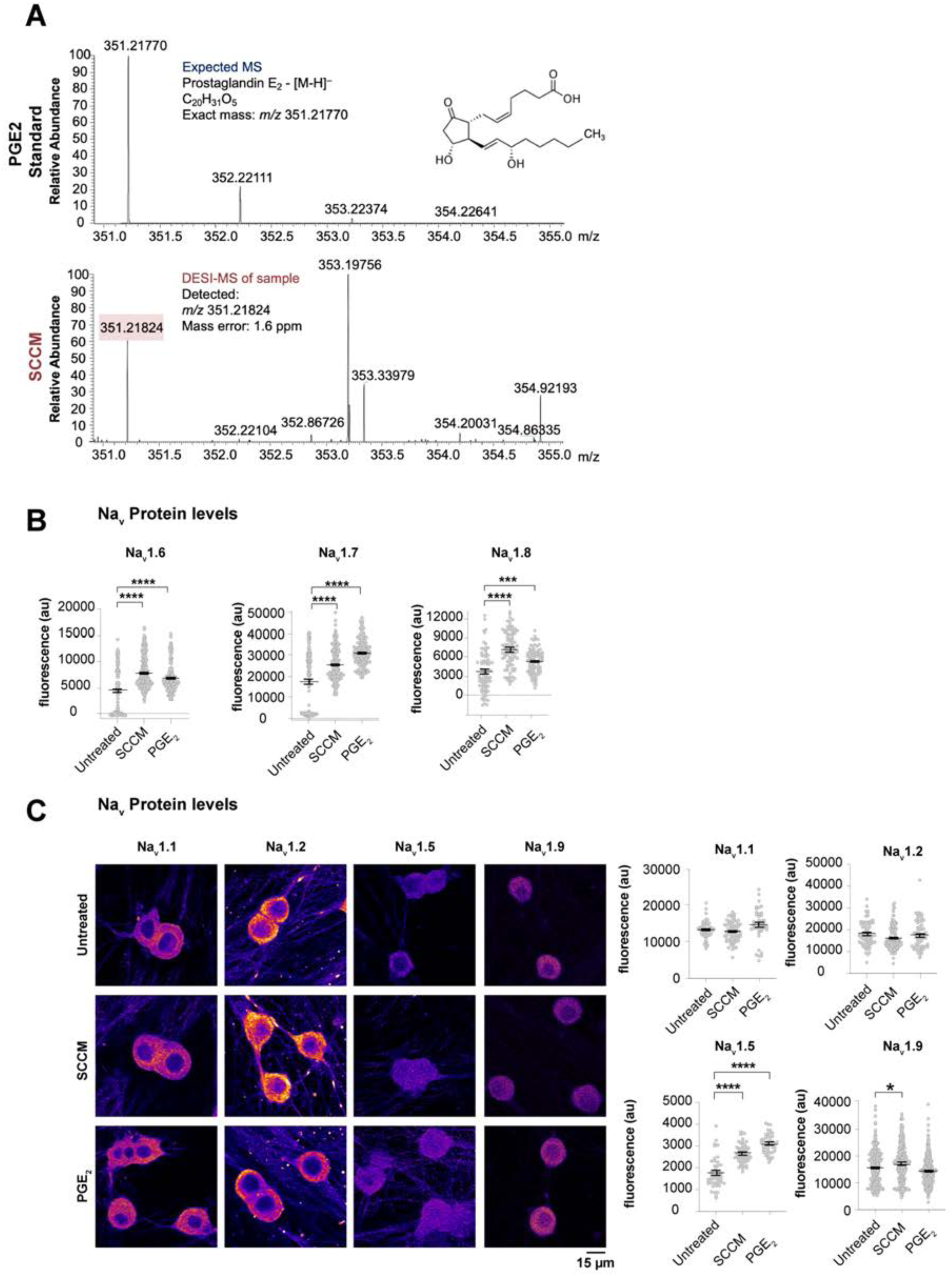
SCCM mass spectrometry and additional immunostaining data, supporting Figures 1 and 2. (**A**) DESI-MS analysis of SCCM. The expected m/z of PGE_2_ is 351.21770 and a m/z of 351.21824 ± 1.6 ppm was detected in the SCCM sample. (**B**) SCCM and PGE_2_ treatments enhanced Na_V_1.6, Na_v_1.7 and Na_v_1.8 protein levels in DRG neurons. *n* = 80 to 328 cells from 1-4 distinct biological replicates per treatment group. Mean ± SEM is shown for cells. Untreated and SCCM conditions are replotted from Figures 1K-1M, as these experiments were done concurrently. (**C**) SCCM and PGE_2_ treatments did not cause a major change in protein expression of indicated genes, except a significant change in Na_v_1.5 protein levels. Fluorescence levels were quantified using cellpose. Gray circles, fluorescent intensity of an individual DRG neuron for the indicated genes. *n* = 70 to 331 cells from 2-4 distinct biological replicates per treatment group. Mean ± SEM is shown for cells. p values compare cells in a one-way ANOVA and Tukey test, statistically significant increases compared to untreated DRG neuron condition are indicated in brackets. *p<0.05, **p<0.01, ***p<0.001, ****p<0.0001.

**Figure S5.**
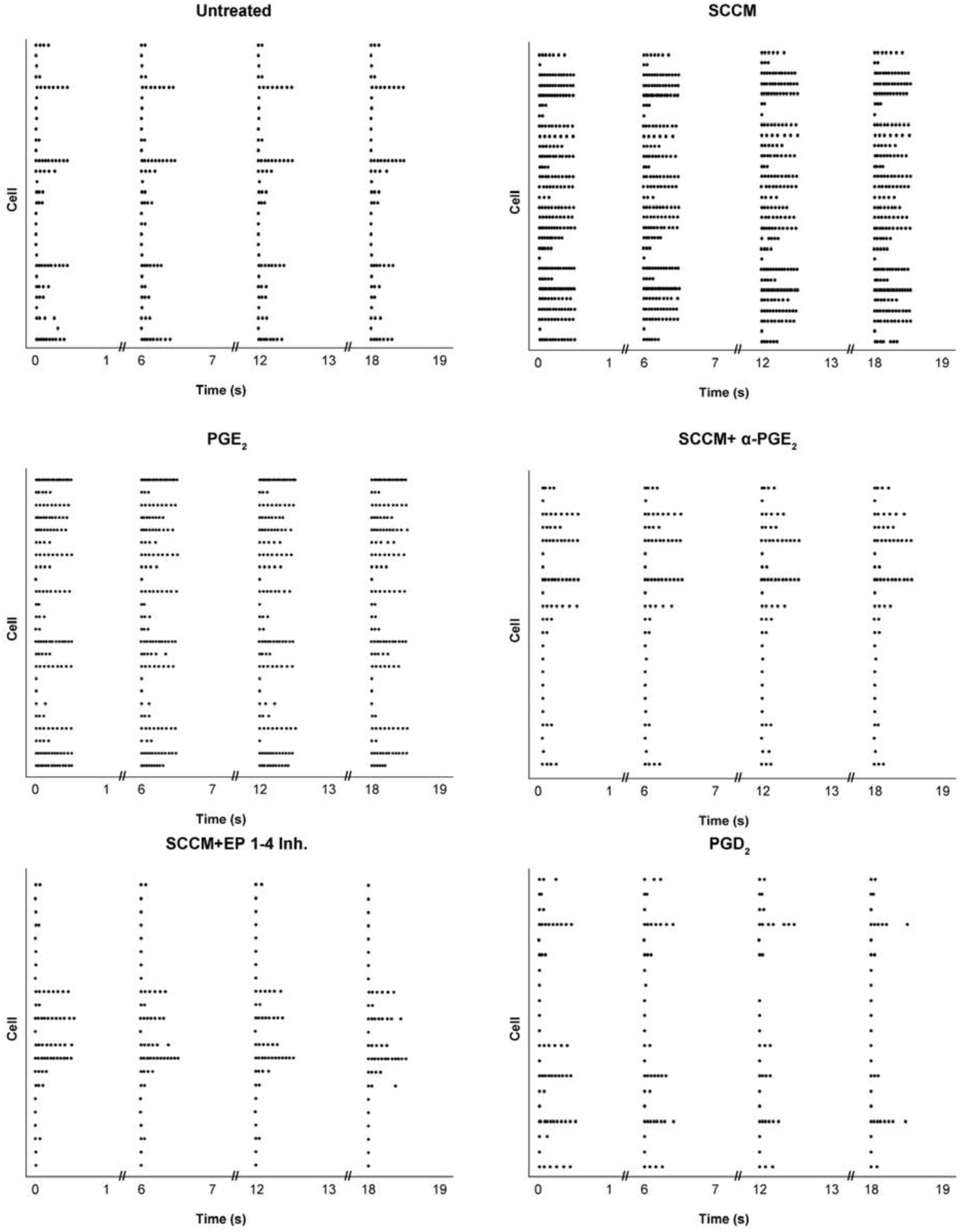
PGE_2_ is necessary and sufficient to induce excitability of DRG neurons. Raster plots of DRG neurons treated with different conditions (electrophysiology experiments shown in Figures 1 and 2). Each row on the y-axis represents a recording from a single DRG neuron. Each dot represents one action potential fired by a single DRG neuron. Raster plots show the excitability of DRG neurons treated under different conditions. SCCM and PGE_2_ treatment markedly increased the number of action potentials fired by DRG neurons. PGE_2_ function-blocking antibodies or EP1–4 inhibitor cocktail blocked the excitability-inducing effect of SCCM. PGD_2_, a structural isomer of PGE_2_, did not show the excitability-inducing effect.

**Figure S6.**
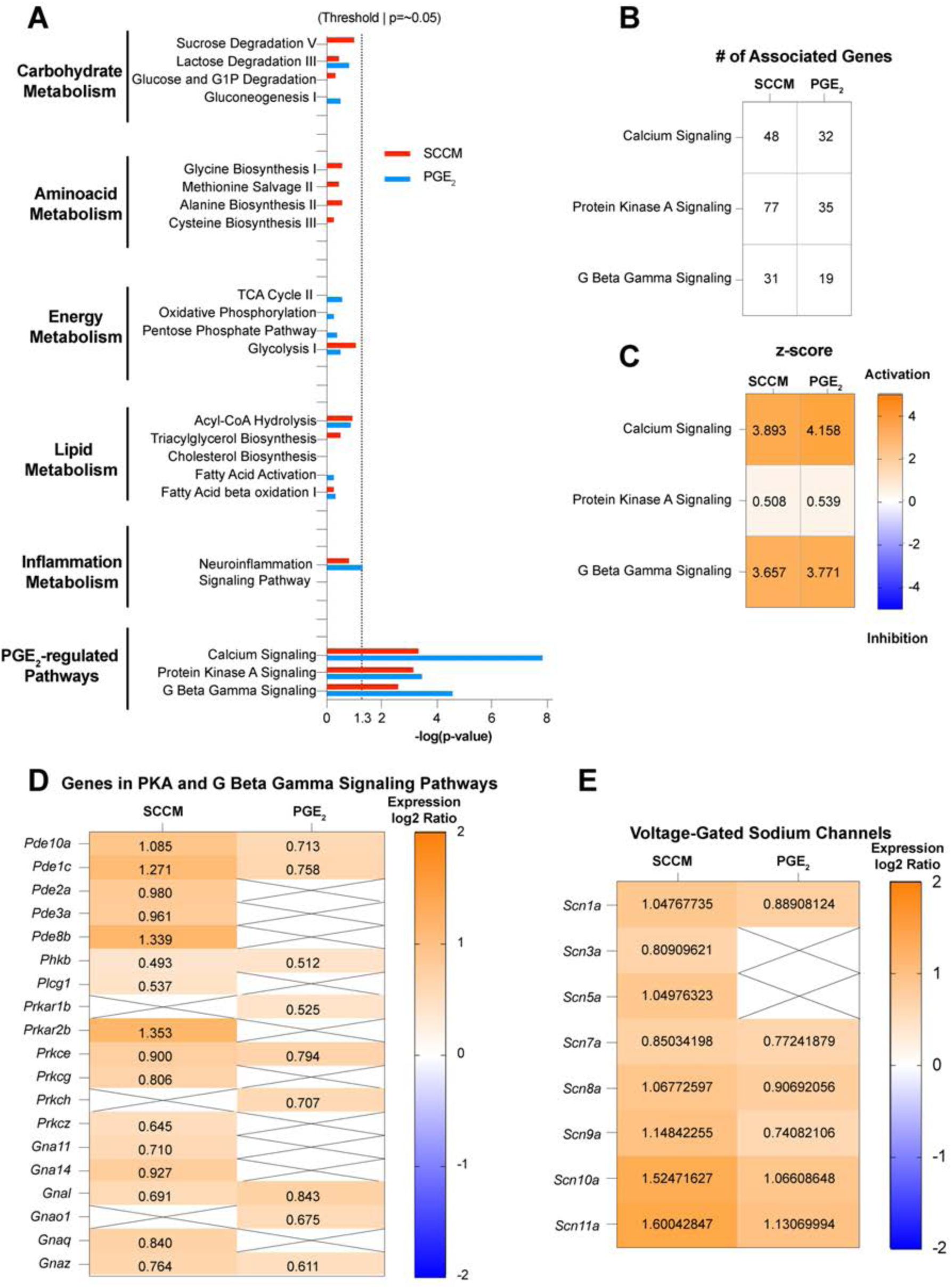
Schwann cell-secreted factors or PGE_2_ treatment increases PGE_2_ downstream pathways without changing inflammatory or metabolic pathways. (**A**) We analyzed activated pathways in SCCM or PGE_2_-treated DRG neurons using IPA. IPA results show a significant correlation to possible PGE_2_ downstream signaling pathways but no correlation to metabolic or inflammatory pathways. (**B**) PGE_2_ or SCCM treatment causes significant changes in genes associated with the indicated pathways. (**C**) z-score analysis suggests activation of the indicated pathways in SCCM or PGE_2_-treated neurons. (**D** and **E**) Diagrams show upregulation of genes in PKA or G beta Gamma signaling pathways, and Na_v_s in SCCM or PGE_2_-treated neurons.

**Figure S7.**
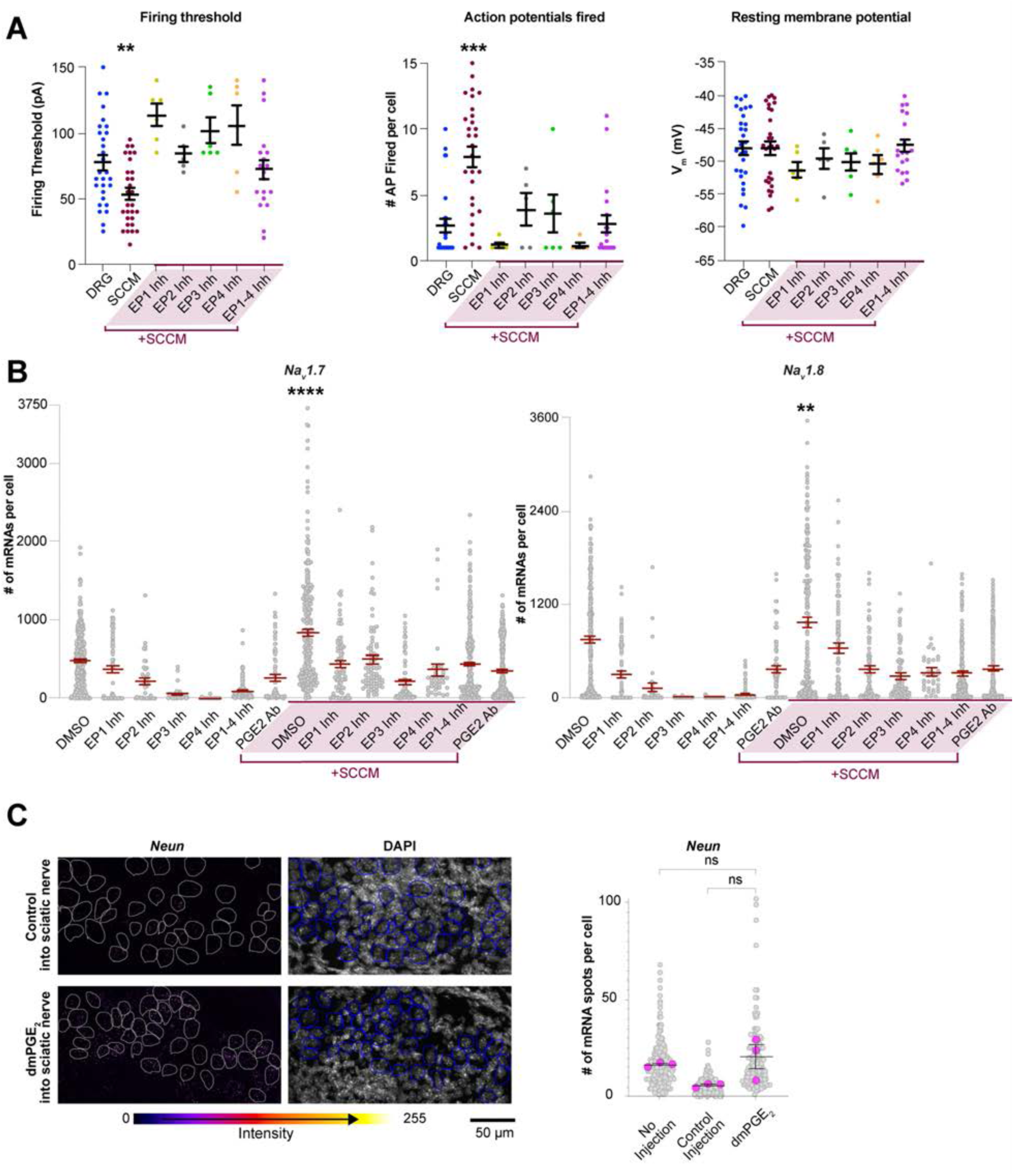
Inhibiting PGE_2_ signaling blocks the ability of SCCM to increase the excitability of DRG neurons. (**A**) Individual EP receptor inhibitors (ONO-8711 [EP1 inhibitor], PF-04418948 [EP2 inhibitor], L-798106 [EP3 inhibitor], and AH 23848 [EP4 inhibitor]) reduced the ability of SCCM to lower the firing threshold or increase the number of fired action potentials in DRG neurons. Untreated DRG neurons and DRG neurons treated with SCCM+EP1–4 inhibitors exhibited statistically similar resting membrane potentials. Mean ± SEM is shown for cells, p values compare cells in a one-way ANOVA and Tukey test. *n* = 6–29 DRG neurons per group. (**B**) Individual EP receptor inhibitors blocked the transcriptional increase of Na_V_s following SCCM treatment. Gray circles, mRNA number of an individual DRG neuron for the indicated genes. Mean ± SEM of all cells, p values compare cells in a one-way ANOVA and Tukey test. Only the DMSO+SCCM condition exhibited a statistically significant increase in mRNA counts compared to DMSO controls. *n* = 17-321 DRG neurons from 2–4 biological replicates per group. p<0.05, **p<0.01, ***p<0.001, ****p<0.0001. (**C**) RNAscope shows *NeuN* expression and DAPI in lumbar DRG neurons at P0 following control or dmPGE2 injections, the anatomy of DRG ganglia and *Neun* expression was unchanged (related to Figures 2H and 2I).

**Figure S8.**
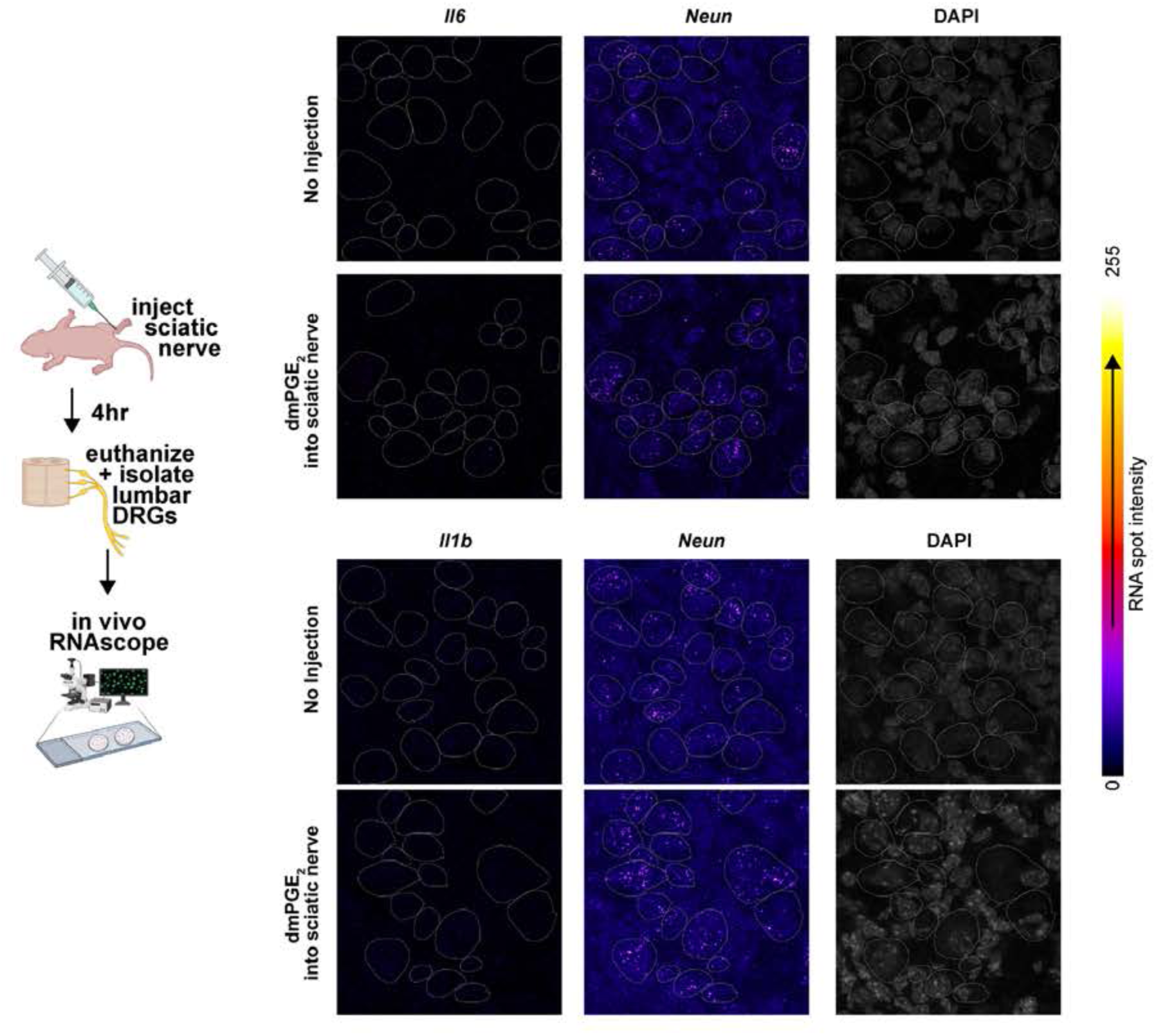
PGE_2_ injection into the sciatic nerve does not cause upregulation of inflammatory markers. RNAscope shows that injection of PGE_2_ into the sciatic nerve does not cause upregulation of inflammatory markers *Il6* and *Il1b* in DRG neurons. These results indicate that PGE_2_ injection causes upregulation of *Na_v_ 1.7* and *Na_v_ 1.8* (See Figures 2H and 2I) without causing neuronal inflammation.

**Figure S9.**
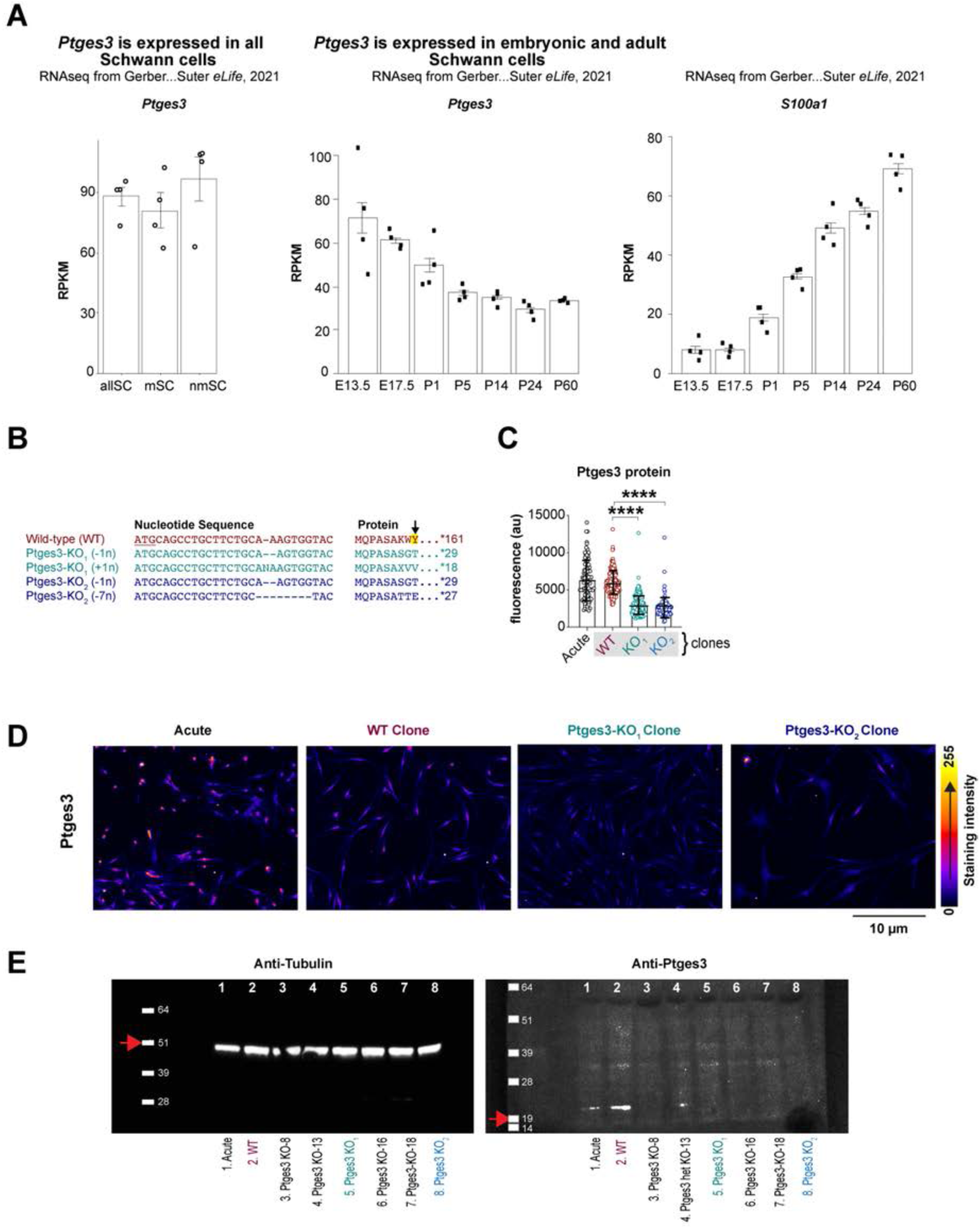
Ptges3 is required for Schwann cell PGE_2_ synthesis. (**A**) Published RNAseq study^40^ shows that both myelinating and nonmyelinating Schwann cells express *Ptges3*. *Ptges3* expression levels are comparable to Schwann cell-specific marker *S100a1*. (**B**) DNA sequencing predicted biallelic mutations resulting in loss of the catalytic residue (arrow) and premature stop codons in *Ptges3* in both KO colonies. (**C**) Ptges3 protein was reduced in Ptges3-KO colonies, *n* = 97, 172, 212 and 100 cells per group respectively. (**D**) Ptges3 expression was lost from *Ptges3* knockout rat Schwann cell colonies. A wild-type colony created in parallel showed similar Ptges3 expression to that of acutely purified Schwann cells (WT). (**E**) Western blot shows Tubulin (51 kDa) and Ptges3 (19 kDa) expression (indicated by red arrows) in acutely purified (WT) or wild-type/KO Schwann cell colonies. Tubulin expression was similar in all collected samples. Ptges3 expression was similar to acutely isolated WT Schwann cells (Acute) and an isolated WT colony (WT), and reduced in an isolated *Ptges3* heterozygous mutant colony (Ptges3 het KO-13). Ptges3 expression was lost in five other isolated colonies that were confirmed to carry biallelic, loss-of-function lesions in *Ptges3* locus. Two *Ptges3* knockout colonies (KO_1_ and KO_2_) were propagated for further studies.

**Figure S10.**
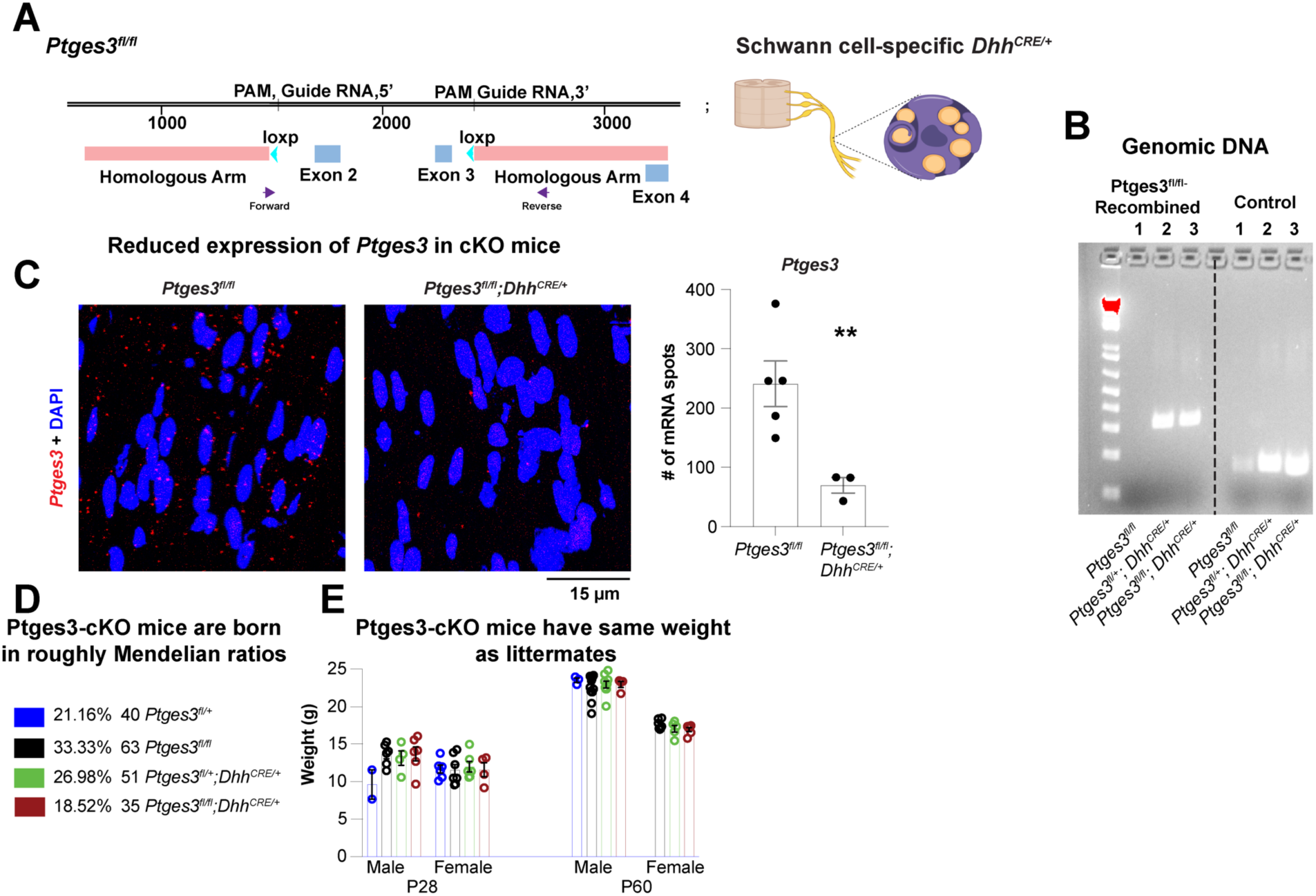
Development of *Ptges3* conditional knockout mice. (**A**) *Ptges3* locus in Ptges3^fl/fl^ mice. Loxp sequences were inserted using CRISPR/Cas9 editing to exclude exons 2 and 3 of the *Ptges3* locus. Target sites for the guide RNAs and homologous arms used for the insertion of loxp sites are indicated. *Ptges3^fl/fl^* mice were crossed to *Dhh^CRE/+^* mice to induce *Ptges3* exon 2 and 3 knockout specifically in myelinating and nonmyelinating Schwann cells. (**B**) PCR amplification results showed the recombined band in the *Ptges3* locus using the forward and reverse primers indicated in (A). PCR only amplified the recombined locus in *Ptges3^fl/fl^;Dhh^CRE^* and *Ptges3^fl/+^;Dhh^CRE/+^* mice but not in control (*Ptges3^fl/fl^*) mice. A control PCR experiment targeting the *Mbp* locus showed amplification in all DNA samples. (**C**) RNAscope shows *Ptges3* expression in *Ptges3* Schwann cell conditional knockout and control mice. *Ptges3* expression was significantly reduced. The graph shows the number of *Ptges3* mRNA per 10^4^ µm^2^ (area of the micrograph) quantified using FishQuant. Mean ± SEM is shown for biological replicates. p values compare the biological replicates in an unpaired t-test. *n* = 5 and 3 mice. (**D**) *Ptges3* conditional knockout mice were generated by crossing *Ptges3^fl/+^;Dhh^CRE/+^* mice to *Ptges3^fl/fl^* mice with 25% of expected Mendelian ratios for each indicated genotype. Chi-square test showed a significant deviation from 25% expected ratios, Ptges3 conditional knockout mice were observed slightly less frequently, at 18.5% (*n* = 189 total, p = 0.02 in Chi-square test). (**E**) Weights of *Ptges3* conditional knockout mice were not significantly different than that of the control littermates at P28 or P60 when compared in a one-way ANOVA and Tukey test performed among each sex and age group. *n* = 4–13 mice per group. Mean ± SEM is shown for biological replicates, compared in a one-way ANOVA and Tukey test. *p<0.05, **p<0.01, ***p<0.001, ****p<0.0001.

**Figure S11.**
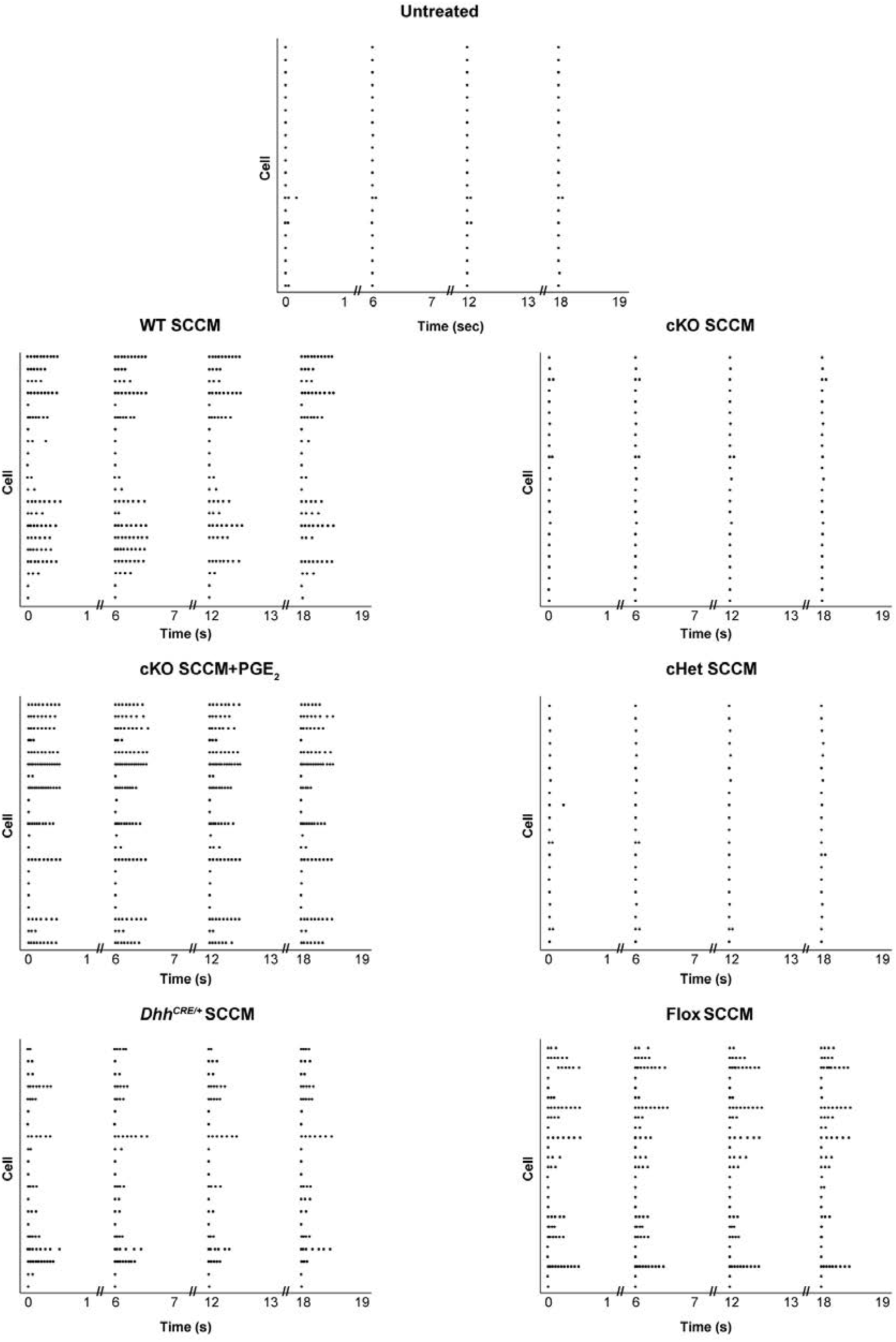
Ptges3 loss-of-function reduces the ability of Schwann cells to induce neuronal excitability. Raster plots of DRG neurons treated with SCCM collected from Schwann cells of indicated genotypes (electrophysiology experiments shown in Figures 3H to 3J). Each row on the y-axis represents a recording from a single DRG neuron. Each dot represents one action potential fired by a DRG neuron. SCCM from control Schwann cells (*Dhh^CRE/+^*, Flox [*Ptges3^fl/fl^*] or WT) increased the number of action potentials fired by DRG neurons. cKO (*Ptges3^fl/fl^;Dhh^CRE/+^*) or cHET (*Ptges3^fl/+^;Dhh^CRE/+^*) SCCM did not induce this effect, suggesting that *Ptges3* heterozygous or complete loss-of-function impair the ability of Schwann cells to induce DRG neuron excitability.

**Figure S12.**
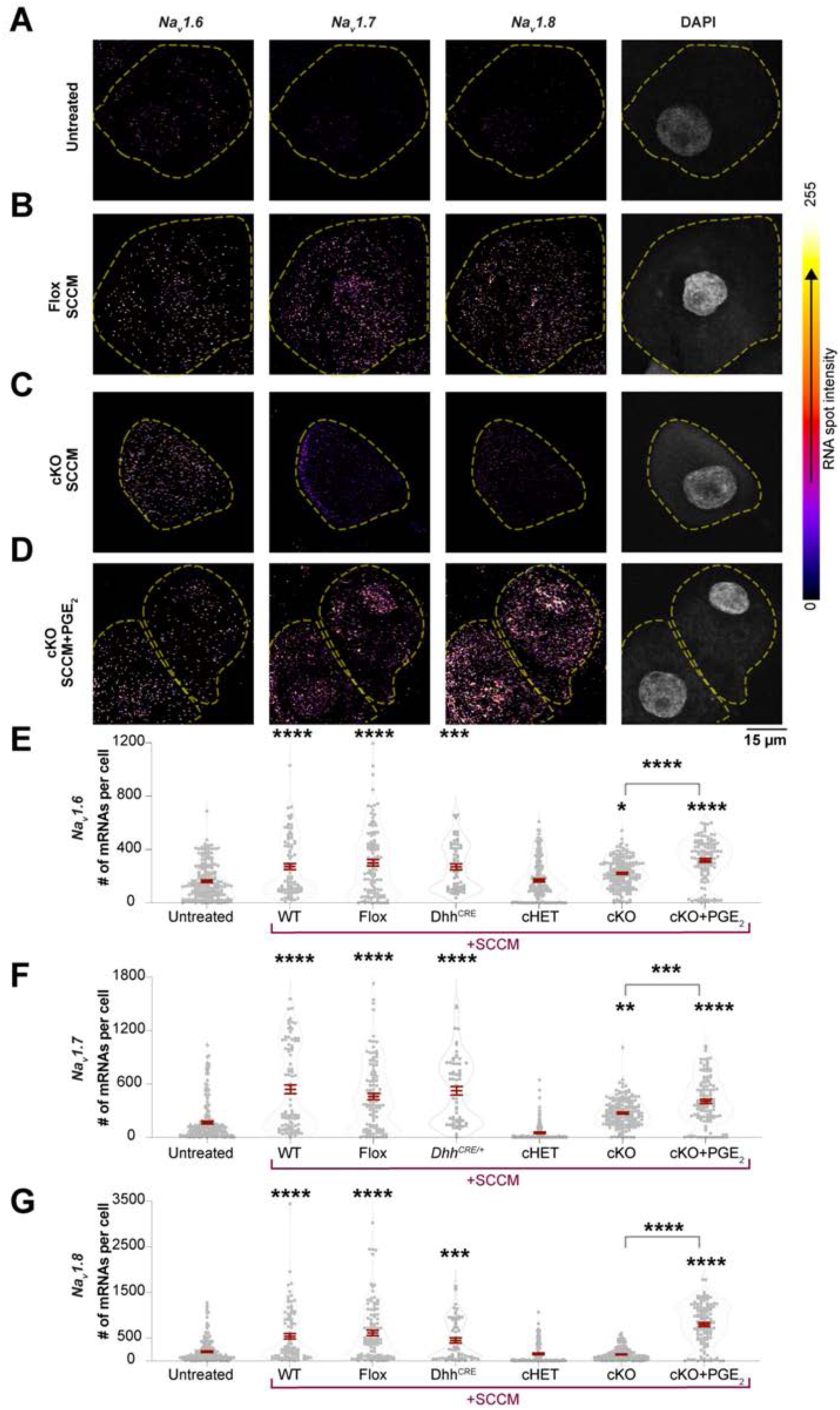
Ptges3 loss-of-function reduces the ability of Schwann cells to increase the expression of Na_V_s in cultured DRG neurons. (**A** to **D**) RNAscope shows expression of *Na_V_1.6*, *Na_V_1.7* and *Na_V_1.8* transcripts in DRG neurons treated with SCCM from Flox (*Ptges3^fl/fl^*) or cKO (*Ptges3^fl/fl^;Dhh^CRE/+^*) mice. Micrographs of SCCM treatment from cHET (*Ptges3^fl/+^;Dhh^CRE/+^*), *Dhh^CRE/+^* and WT mice are not shown but are also quantified in (E to G). (**E** to **G**) Conditioned media from *Ptges3* conditional knockout Schwann cells (cKO) failed to enhance neuronal *Na_V_1.8* expression, contrary to Schwann cells collected from controls (Flox, Dhh^CRE/+^, or WT). We detected a slight but significant change in *Na_V_1.6* and *Na_V_1.7* expression in DRG neurons following the addition of cKO media; however, this effect was lower than the fold increase observed with WT, Flox, or Dhh^CRE/+^ SCCM addition. The addition of PGE_2_ to cKO media rescued the loss of Na_v_ expression-inducing effect in cKO SCCM. Gray circles, mRNA number of an individual DRG neuron for the indicated genes. Mean ± SEM of all cells; p values compare cells in a one-way ANOVA and Tukey test. Significant increases in comparison to untreated DRG neurons or between groups indicated by brackets are shown. *n* = 61–195 DRG neurons from 2–4 biological replicates per group. p<0.05, **p<0.01, ***p<0.001, ****p<0.0001.

**Figure S13.**
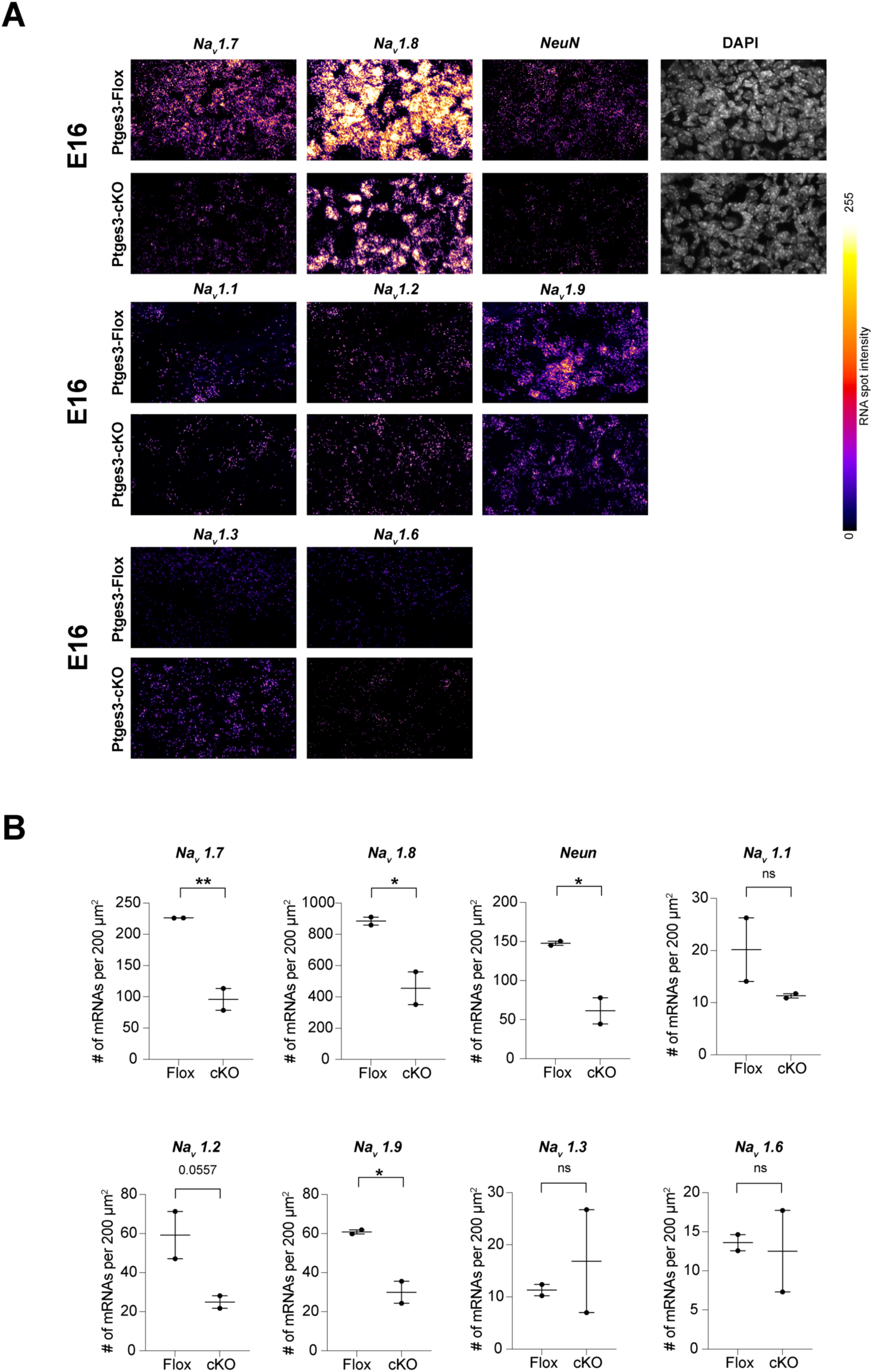
Schwann cells promote neuronal maturation and Na_v_ expression during embryonic development. (**A**) RNAscope shows expression of the indicated Na_v_ transcripts in lumbar DRG neurons at embryonic day 16 (E16). *n* = 2 mice per group. (**B**) Number of *Na_v_* mRNAs per 200 μm^2^ was quantified using FishQuant. Mean ± SEM is shown for biological replicates (mice). p values compare biological replicates in an unpaired t-test. *p<0.05, **p<0.01, ***p<0.001, ****p<0.0001.

**Figure S14.**
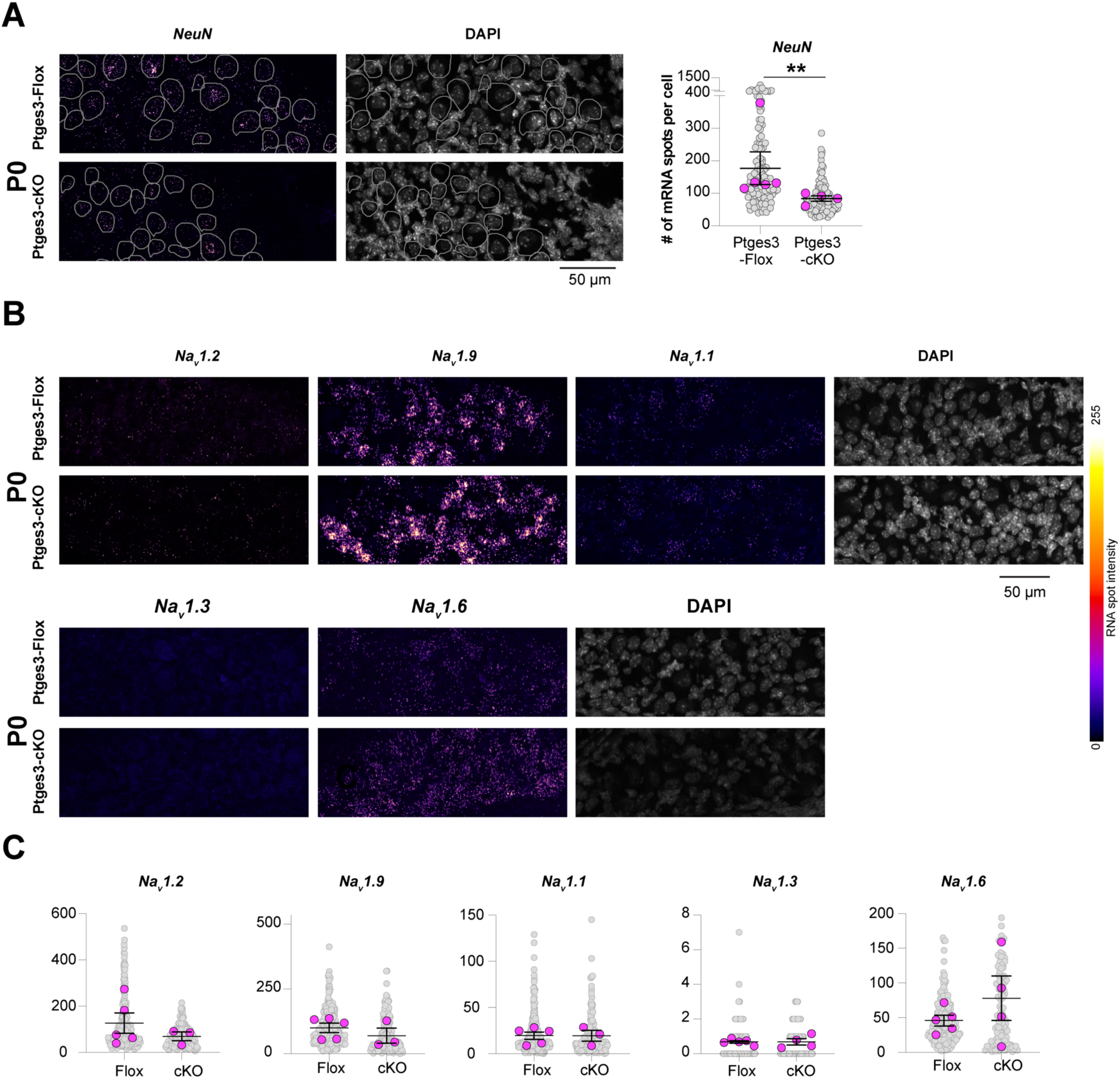
Schwann cells promote neuronal maturation and Na_v_ expression during development. (**A**) RNAscope shows expression of *NeuN* and DAPI in lumbar DRG neurons at the day of birth (P0). *n* = 4-5 mice per group. Number of *NeuN* mRNAs per cell was quantified using FishQuant. Mean ± SEM is shown for biological replicates (mice). p values compare biological replicates in Mann-Whitney test. (**B**) RNAscope shows expression of the indicated *Na_v_* transcripts in lumbar DRG neurons at P0. *n* = 3-5 mice per group. (**C**) Number of *Na_v_* mRNAs cell was quantified using FishQuant. Mean ± SEM is shown for biological replicates (mice). p values compare biological replicates in an unpaired t-test.*p<0.05, **p<0.01, ***p<0.001, ****p<0.0001.

**Figure S15.**
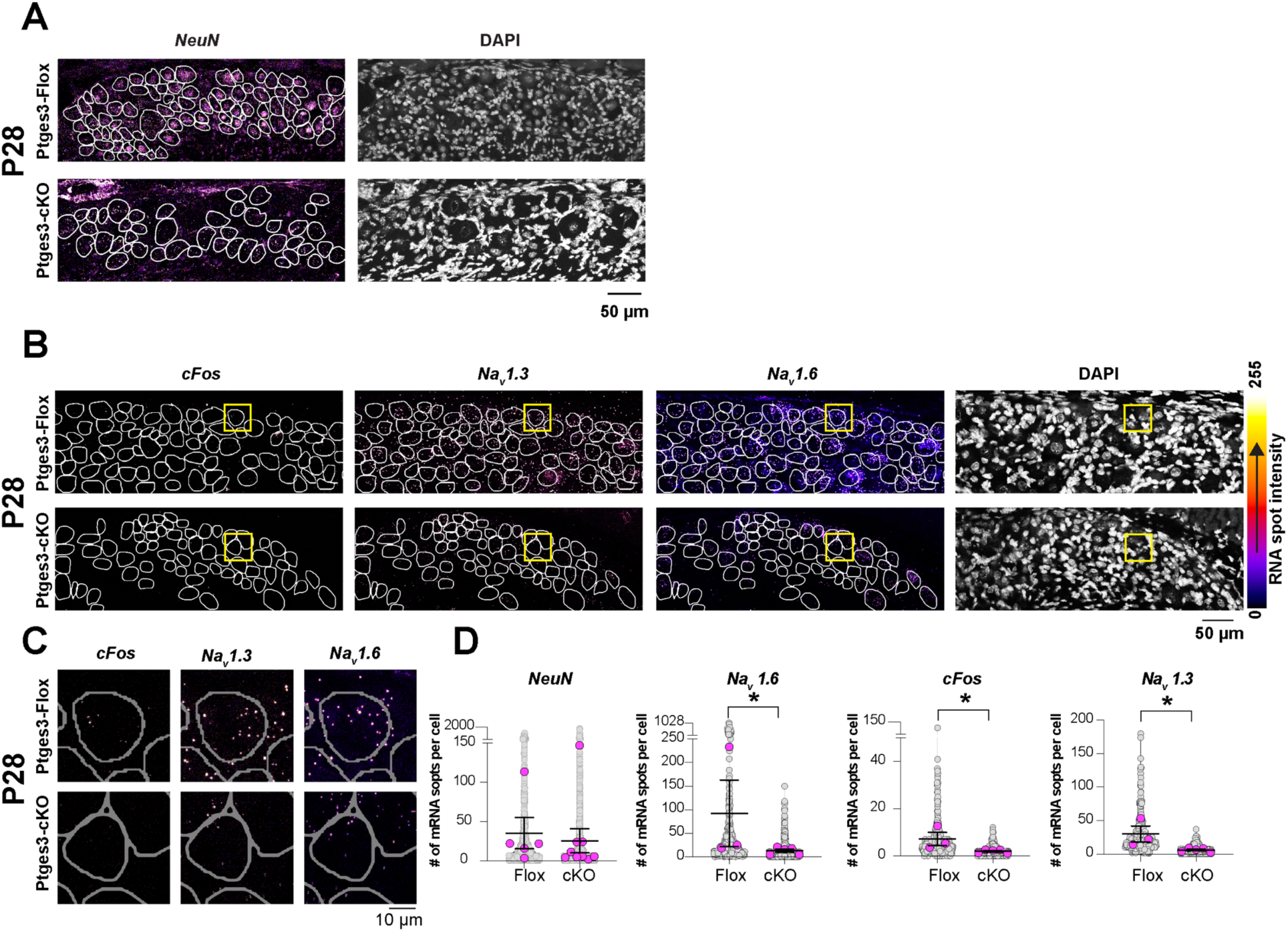
Schwann cells promote neuronal maturation and Na_v_ expression at late postnatal stages. **(A** and **B)** RNAscope shows expression of *NeuN*, indicated *Na_v_* transcripts and *cFos* in lumbar DRG neurons at P28. (**C** and **D**) Insets highlighted with yellow rectangles (B) are magnified for visualization. *n* = 3-9 mice per group. Number of *Na_v_* mRNAs cell was quantified using FishQuant. Mean ± SEM is shown for biological replicates (mice). p values compare biological replicates in Mann-Whitney test. *p<0.05, **p<0.01, ***p<0.001, ****p<0.0001.

**Figure S16.**
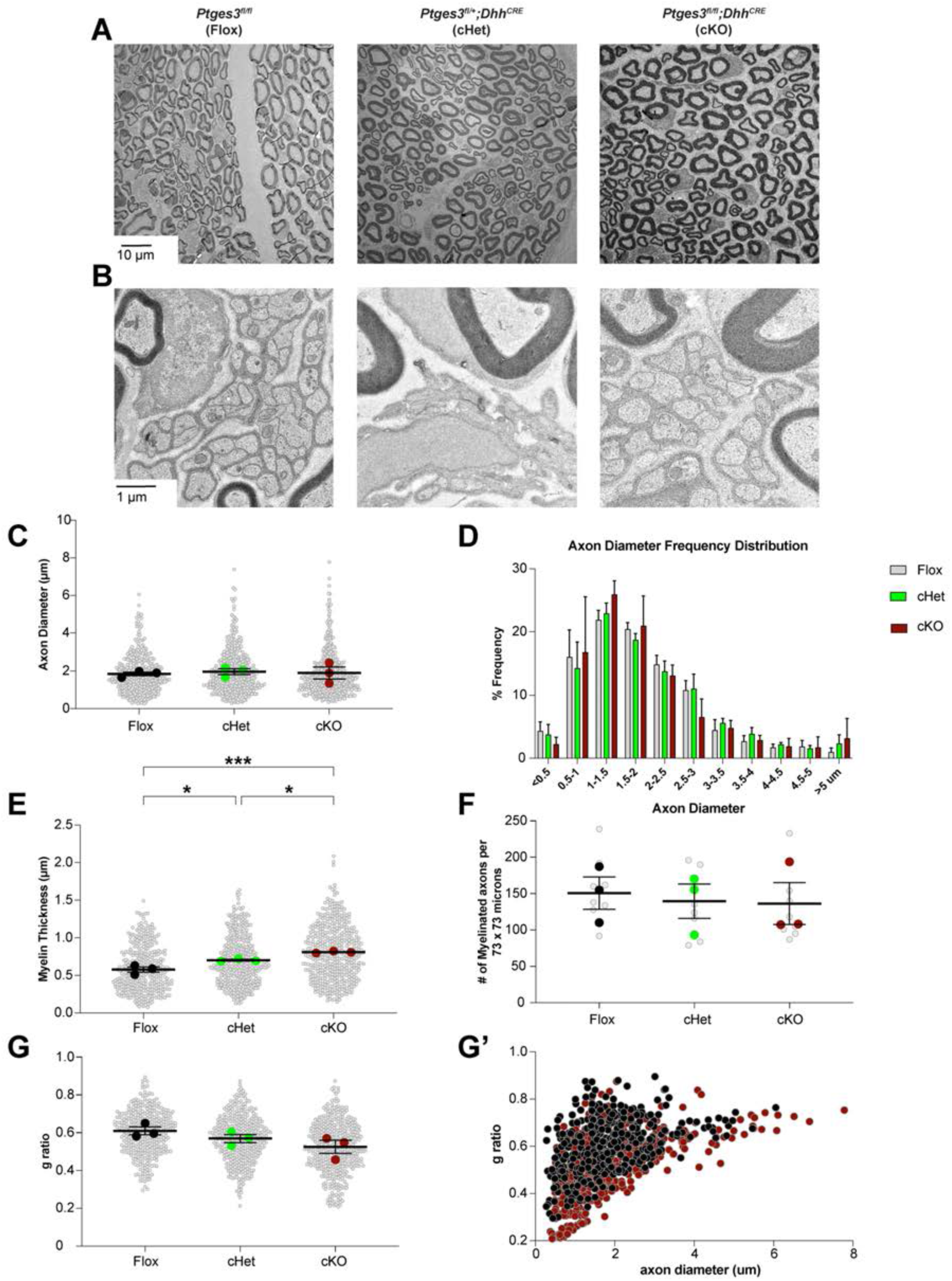
Sciatic nerve anatomy is grossly normal in *Ptges3* conditional knockout mice. (**A**) Images show a cross-sectional area of the sciatic nerve in mice with indicated genotypes at P28 using transmission electron microscopy. The number of myelinated axons per field of view (73 x 73 µm^2^) was statistically similar in *Ptges3* conditional full, conditional heterozygous mice compared to controls. (**B**) Images show that Remak bundles formed normally in all groups (**C, E,** and **G**) Quantification of axon diameter, myelin thickness, and *g*-ratio (the ratio of the inner diameter of an axon to its total outer diameter). Gray circles, measurement of a single axon; colored circles, the average for each biological replicate (1 mouse). Mean ± SEM is shown for biological replicates. p values compare biological replicates in a one-way ANOVA and Tukey test, *p<0.05, **p<0.01, ***p<0.001. (**D**) Percent frequency of the diameter of myelinated axons in the sciatic nerve. No significant differences were detected in a two-way-ANOVA and Tukey test. (**F**) Number of myelinated axons per 73 x 73 µm^2^ (area of the micrographs in (A)) was not statistically different between genotypes. Gray circles, the number of myelinated axons per micrograph; colored circles, the average of the quantified micrographs in each biological replicate (mouse). Mean ± SEM is shown for biological replicates compared in a one-way ANOVA and Tukey test. (**G** and **G’**) *g*-ratio of axons in controls and *Ptges3* conditional full knockout mice (cKO). 90–170 axons per mouse were analyzed, *n* = 3 mice per genotype.

**Figure S17.**
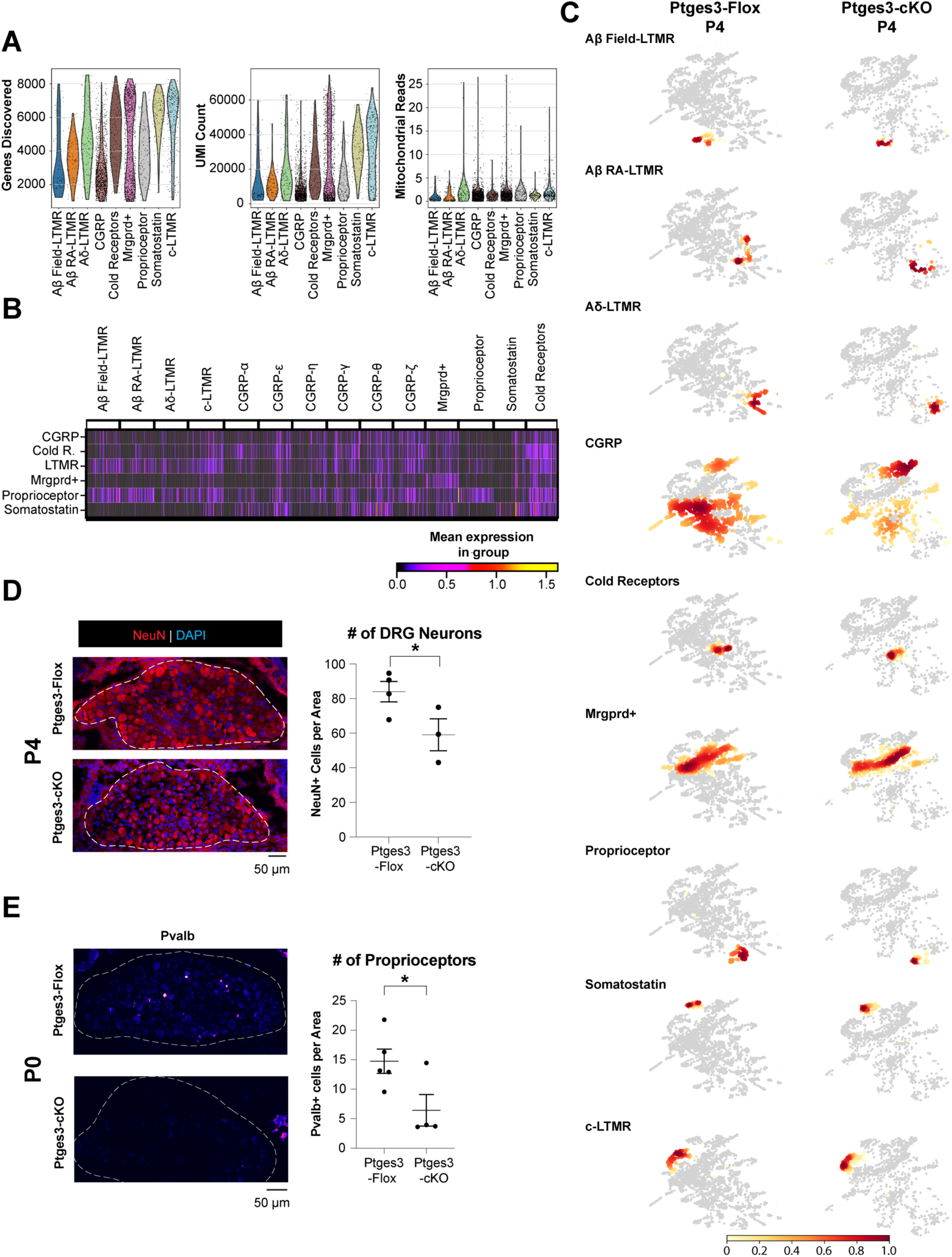
Quality control, clustering and cell density distribution of scRNA-seq data. (**A**) Number of discovered genes and unique molecules in each cell, and percent mitochondrial reads are shown in each population of classified sensory neurons at P4. (**B**) Matrix plot indicates expression of genes that aided classification of the identified DRG neuron subclusters. (**C**) Cell identity composition heatmaps of DRG neuron subtype populations. Dark red indicates high relative cell density. CGRP and proprioceptor subtype population heatmaps shown in Figure 5C are also indicated here to aid comparison to other subtypes. (**D**) Immunohistochemistry indicated a ∼30% decrease in NeuN+ cells in DRG ganglia at P4. N=4 and 3 mice per group. (**E**) Parvalbumin (Pvalb)+ neurons were also decreased by 56%. n=5 and 4 mice per group. Mean ± SEM is shown for biological replicates. p values compare biological replicates in an unpaired t-test. *p<0.05, **p<0.01, ***p<0.001, ****p<0.0001. Graphs show number of NeuN+ (D) or Pval+ (E) cells per 40^3^ μm^2^.

**Figure S18.**
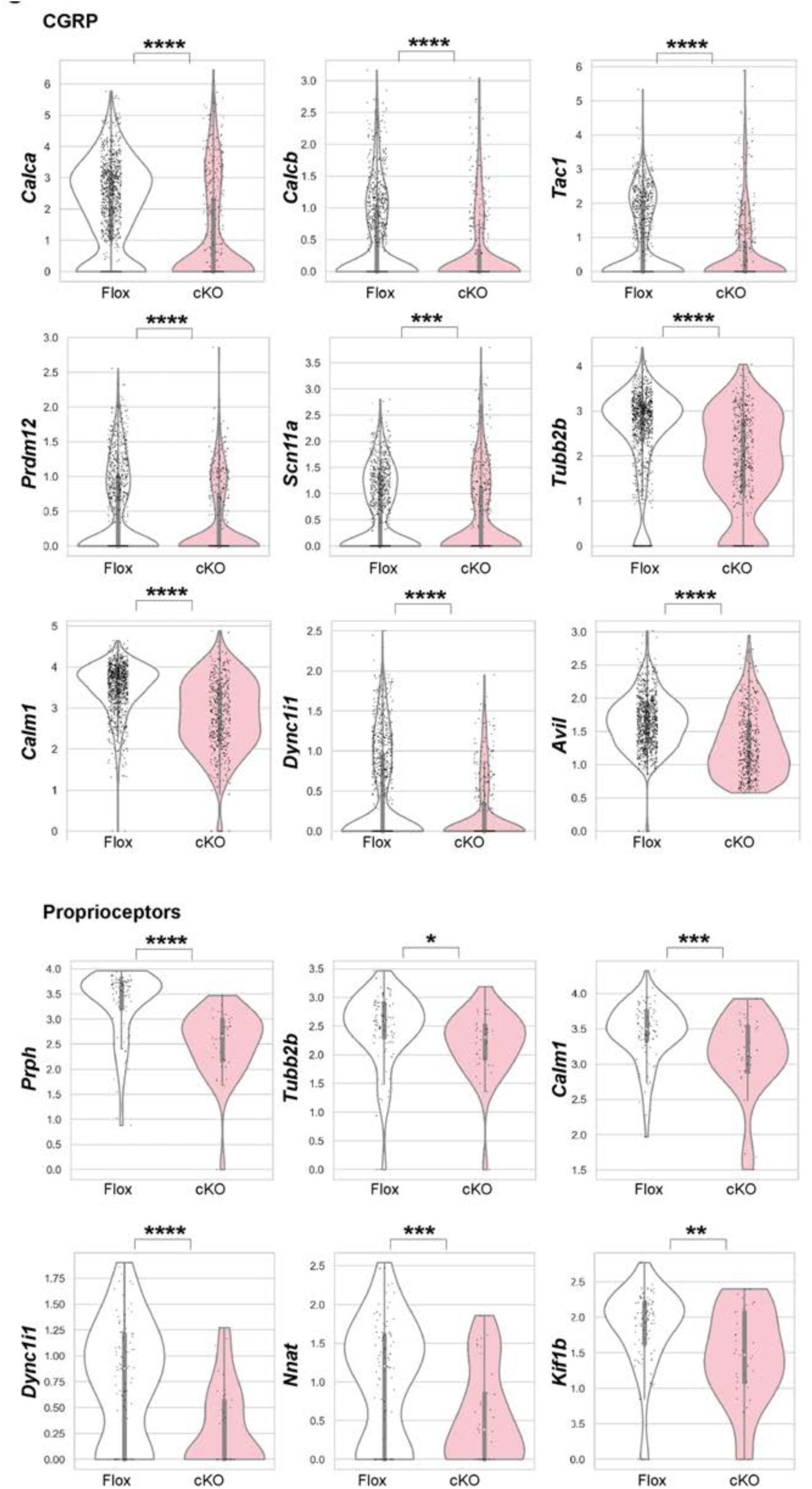
Transcriptomic analysis indicates impaired functional maturation of CGRP and proprioceptor DRG neuron subtypes. Violin plots indicate expression of example genes marking neuronal maturation in CGRP or proprioceptor DRG subpopulations. p values compare cells in an unpaired t-test. *p<0.05, **p<0.01, ***p<0.001, ****p<0.0001.

**Figure S19.**
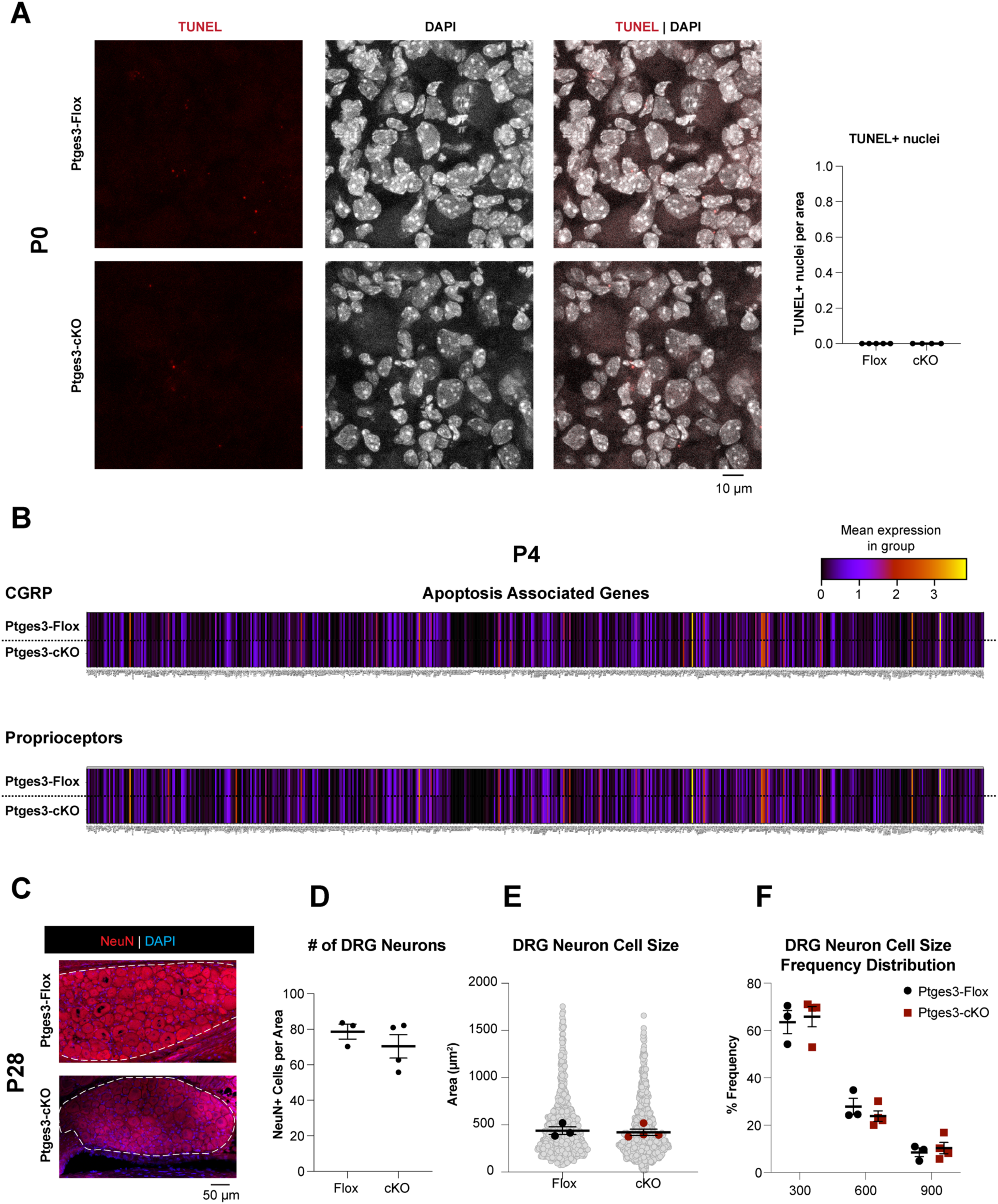
CGRP and proprioceptor DRG neuron subtypes do not express cell death markers in Ptges3-cKO mice. (**A**) TUNEL assay shows no cell death in lumbar DRG ganglia of Ptges3-cKO mice at P0. *n* = 5, 4 mice per group. (**B**) Matrix plot shows the expression of apoptosis gene markers (GO:0097190) in CGRP and proprioceptor DRG subtype cell populations at P4 and suggests no elevation of apoptotic signals in Ptges3-CKO mice. (**C** and **D**) Immunohistochemistry did not indicate a significant decrease in Neun+ cells in DRG ganglia at P28, suggesting that the earlier deficiency in Neun+ cells at P4 (See Figure S17D) likely resulted from a maturation defect rather than cell death. n=3 and 4 mice per group. (**E**) Area of the DRG neuron cell bodies was also not significantly different. Mean ± SEM is shown for biological replicates. p values were not significant in comparing biological replicates in an unpaired t-test. (**F**) Frequency distribution histogram of DRG neuron cell body size revealed no difference in the percentage of small (<300 μm^2^), medium (>300 μm^2^, <600 μm^2^), and large (>900 μm^2^) DRG axons in Ptges3-cKO mice in 2way ANOVA.

**Figure S20.**
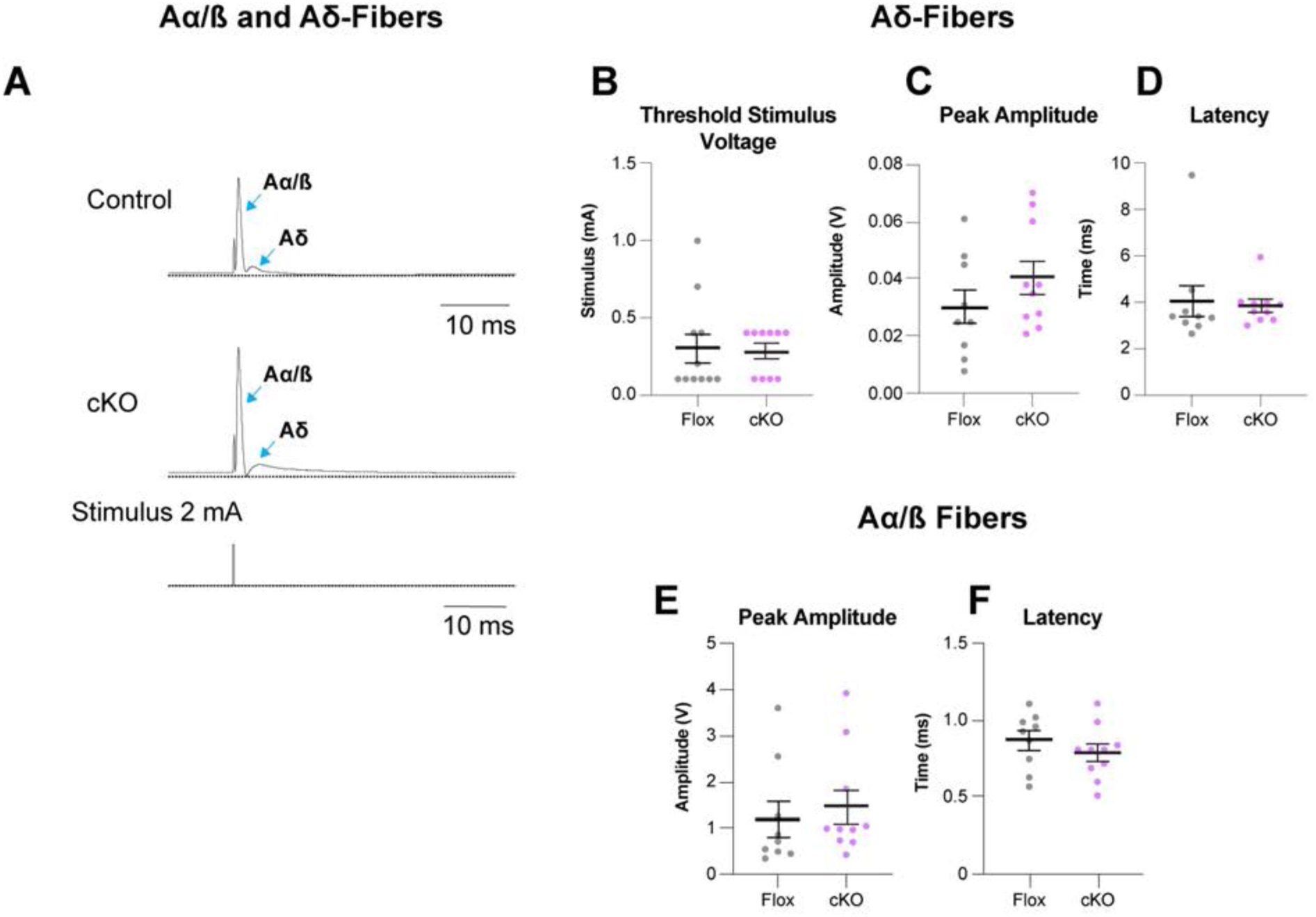
Aα/μ and Aδ fiber excitability is normal in Ptges3-cice. (**A**) Aα/μ and Aδ fiber compound action potentials in Ptges3-Flox and Ptges3-cKO sciatic nerve in response to 2 mA stimulus application. (**B** to **D**) Threshold stimulus voltage, peak amplitude and latency of Aδ fibers are not significantly different in Ptges3-cKO sciatic nerve. (**E** and **F**) Peak amplitude and the latency of the Aα/μ lfibers were not significantly different between Ptges3-cKO and Ptges3-Flox mice. n=10 sciatic nerves from 5 or 6 male mice for each group. p values compare sciatic nerves in Mann-Whitney test.

**Figure S21.**
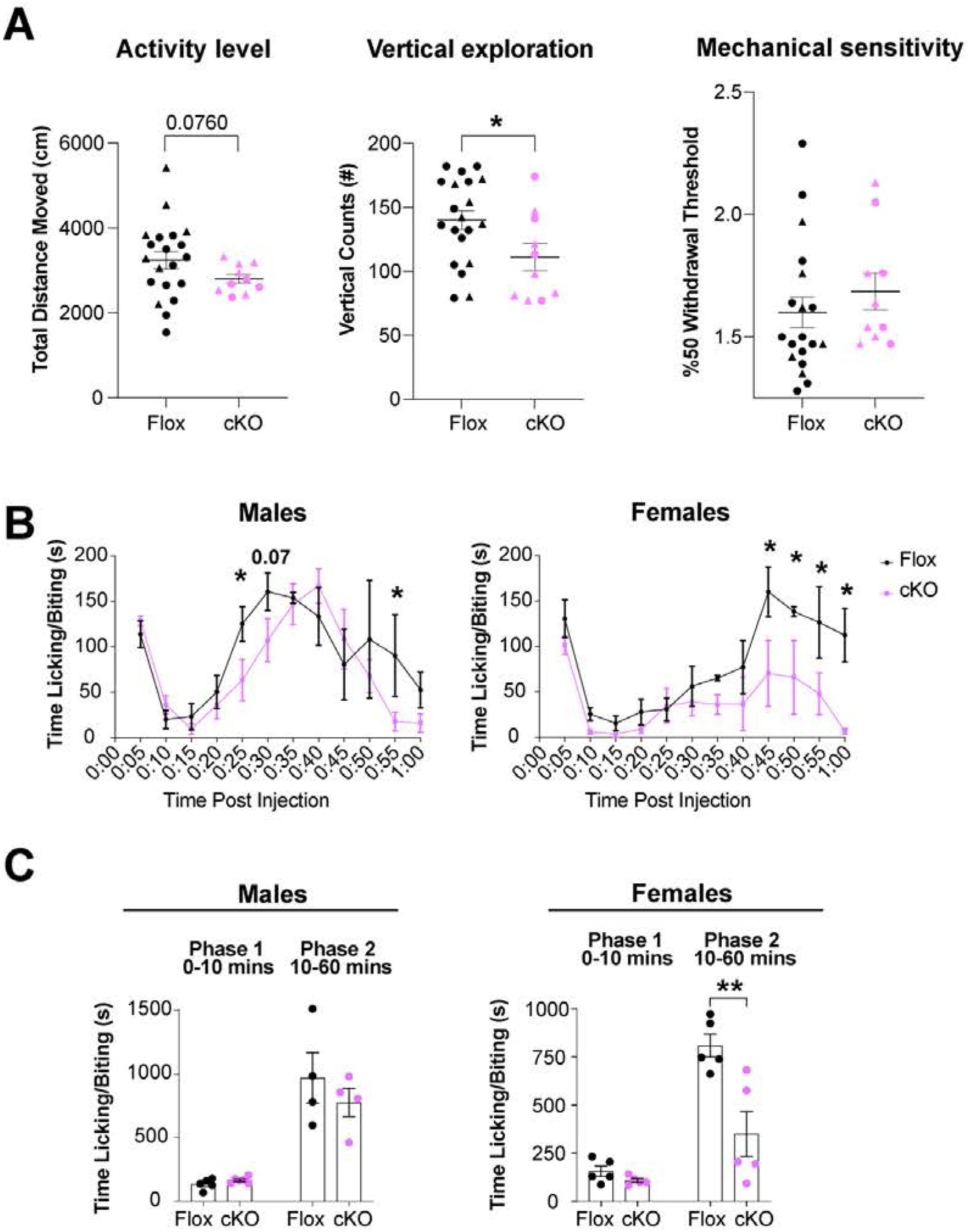
Loss of Schwann cell-secreted PGE_2_ reduces pain behavior. (**A**) Activity chamber assay showed a trend for decreased locomotion and a significant decrease in vertical activity in Schwann cell conditional *Ptges3* knockouts (cKO). n= 20 and 10 mice. In contrast to pain responses, Von Frey test showed no significant changes in mechanical sensitivity between cKO mice and controls. n= 19 and 10 mice (triangles-females, circles-males). (**B** and **C**) Pain response after 1% formalin injection to hind paw is shown in males and females. n= 4-6 males and 5 females per group. Mean ± SEM is shown for each sex at indicated time points, p values indicate statistical differences in a two-way ANOVA test. In all other graphs, p values compare biological replicates in an unpaired t-test. *p<0.05, **p<0.01, ***p<0.001, ****p<0.0001.

**Figure S22.**
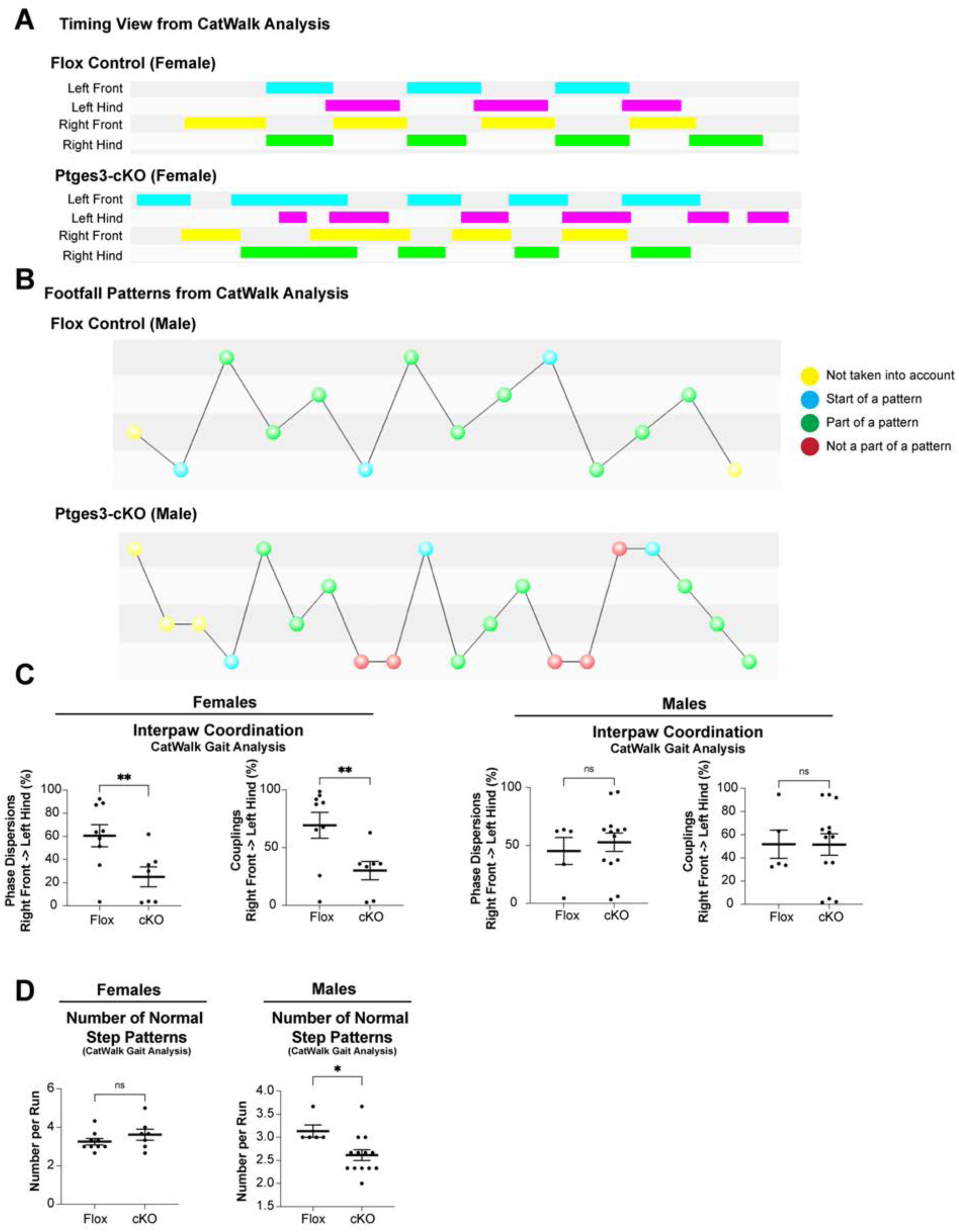
Loss of Schwann cell-secreted PGE_2_ impairs gait and coordination. (**A**) Timing view display shows gait diagrams in a representative control and Ptges3-cKO female mice, and indicates irregular timing of paw placements, suggesting impaired gait. (**B**) Footfall pattern diagram shows abnormal footfall patterns in male Ptges3-cKO mice, suggesting impaired coordination. (**C** and **D**) Graphs indicate impaired interpaw phase dispersion and couplings in female Ptges3-cKO mice and a decrease in normal footfall patterns in male Ptges3-cKO mice. n= 9 females and 5 males for Ptges3-Flox, 7 females and 13 males for Ptges3-cKO mice. p values compare biological replicates in an unpaired t-test. *p<0.05, **p<0.01, ***p<0.001, ****p<0.0001.

**Figure S23.**
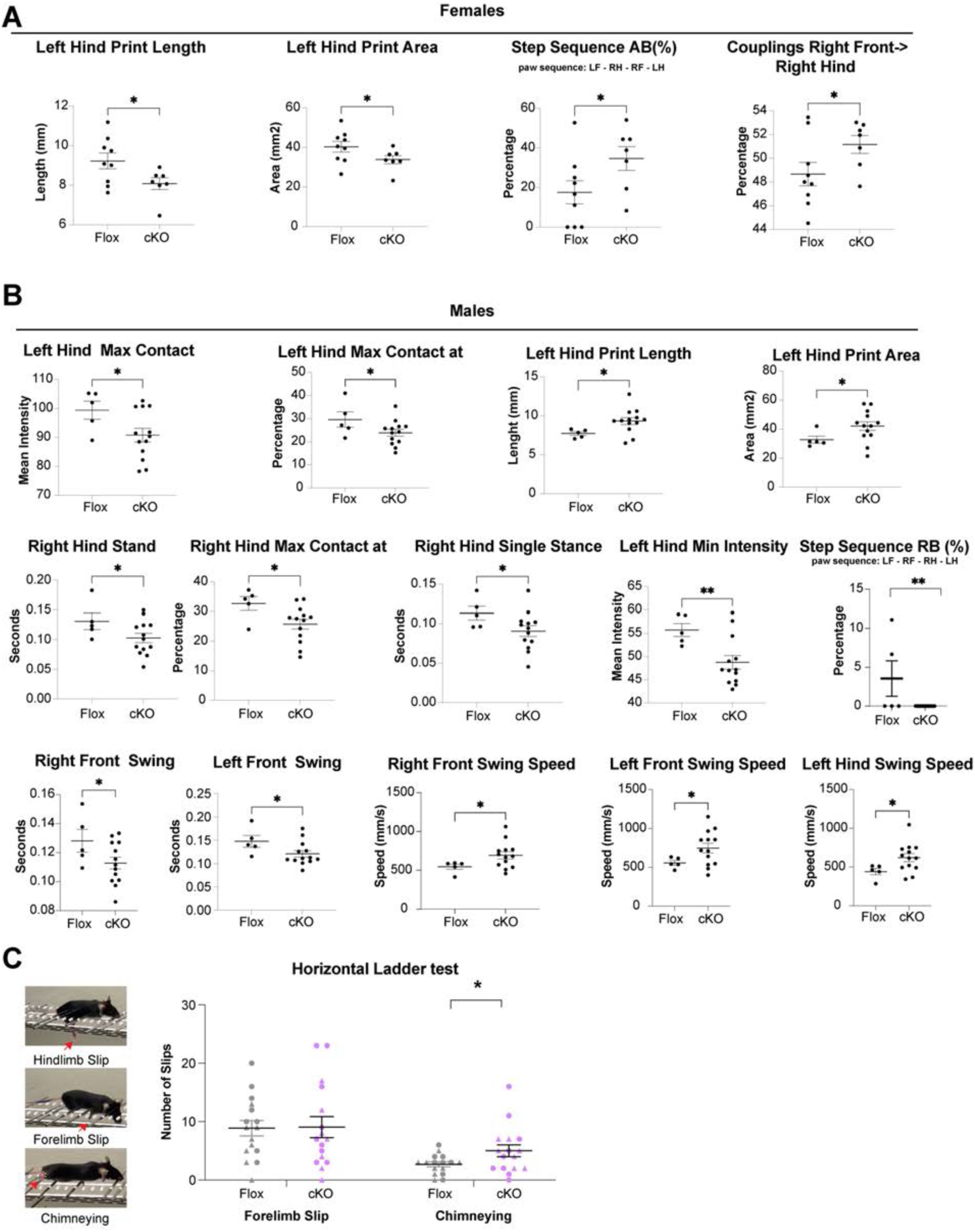
Loss of Schwann cell-secreted PGE_2_ impairs gait parameters and proprioceptive sensory behavior. (**A** and **B**) Graphs indicate additional impaired gait parameters measured in video-based CatWalk XT system (v9.1, Noldus). n= 9 females and 5 males for Ptges3-Flox, 7 females and 13 males for Ptges3-cKO mice. (**C)** Horizontal ladder test with unevenly placed rungs indicated a 92% increase in incorrect hindlimb placement in Ptges3-cKO mice suggesting impaired proprioceptive behavior (See Figure 6I). Forelimb placements were normal. Ptges3-cKO mice also frequently used side transparent bars to progress through the corridor instead of placing limbs on rungs (chimneying). n=16 mice for each group (triangles-females, circles-males). p values compare biological replicates in an unpaired t-test. *p<0.05, **p<0.01, ***p<0.001, ****p<0.0001.

